# Agentic campaign control for high-throughput de novo binder design

**DOI:** 10.64898/2026.09.22.753604

**Authors:** Minkyu Jeon, Jinyeop Song, Jina Kim, Ellen D. Zhong

## Abstract

Progress in artificial intelligence has produced a rapidly growing ecosystem of methods for *de novo* protein design. With access to many specialized and often complementary tools, how does one use them effectively, especially with a finite compute budget? Here, we introduce **Target-adaptive Rescue–Explore–eXploit (T-REX)**, an agentic protein binder design framework that orchestrates multiple state-of-the-art protein generative models and structure evaluators. Over the course of a design campaign, T-REX leverages large language model (LLM) agents to reason over accumulating outcomes and decide whether to rescue promising candidates, explore alternative settings, or exploit productive routes, while a deterministic controller validates and schedules proposed actions. Across seven targets, we find that T-REX adaptively allocates its compute across methods in a target-dependent manner and achieves the highest throughput of structurally distinct hits compared to all baselines. These results suggest that adaptive orchestration can complement increasingly powerful protein design models, and we openly release T-REX to support compute-efficient binder design workflows.

## 1. Introduction

De novo protein design is emerging as a powerful paradigm to create novel proteins of desired structures and molecular functions, with opportunities in therapeutics, catalysis and molecular engineering (Chu et al., 2024; Listov et al., 2024; Bennett et al., 2023; Watson et al., 2023). Recent advances in artificial intelligence have introduced a growing suite of computational tools that are revolutionizing various stages of the protein design pipeline (Yang et al., 2026), including generative models of backbone structures (Watson et al., 2023; Ingraham et al., 2023), inverse folding for sequence design (Ingraham et al., 2019; Dauparas et al., 2022; Hsu et al., 2022), and folding models for structure prediction and evaluation (Jumper et al., 2021; Lin et al., 2023; Abramson et al., 2024; Chai Discovery et al., 2024; Wohlwend et al., 2025; Protenix Team et al., 2026). Joint structure–sequence generative models (Stark et al., 2025; Didi et al., 2026b; Hayes et al., 2025) and integrated hallucinate-and-filter pipelines such as BindCraft (Pacesa et al., 2025) provide yet additional paths to contemporary *de novo* protein binder design.

As the set of individual tools becomes increasingly capable and diverse, a new question emerges: *which methods should be run, with which settings, and how should a finite compute budget be allocated*, especially as the protein design campaign progresses? Here, a campaign denotes the sequence of computational generation, redesign and evaluation jobs carried out for a single target. Notably, these specialized tools can differ in productivity and computational cost (Liu et al., 2026), both between targets and throughout the course of a design campaign. At different stages, candidate designs can miss *in silico* metrics or duplicate existing structures, and tool calls can fail. Additionally, designs that do not initially qualify can still provide promising starting points for subsequent structural or sequence refinement (Vázquez Torres et al., 2024; Bennett et al., 2023). Effective orchestration of available methods therefore requires interpreting these outcomes to decide whether to refine promising designs, continue productive approaches, or test alternative paths.

Recent progress in scientific agents suggests that large language models (LLMs) can provide the contextual reasoning capabilities needed to interpret heterogeneous outcomes and autonomously strategize follow-up actions (Ding et al., 2025; Ghafarollahi and Buehler, 2024; Ouyang et al., 2026; Swanson et al., 2025; Teneggi et al., 2026; Xu et al., 2026). In protein design, LLM agents have begun to coordinate multi-stage computational workflows and iterative design procedures (Ge et al., 2026; Swanson et al., 2025), and recent approaches have shown autonomous protein design albeit with large compute and LLM token budgets (Claude Science and Shanehsazzadeh, 2026). A central control challenge remains in adaptively allocating limited compute over the course of a design campaign. In particular, protein generation, redesign and structure evaluation are computationally intensive, whereas LLM reasoning over compact summaries of campaign outcomes can be comparatively fast. A relatively brief reasoning step can therefore guide more expensive computation by specialized scientific tools. Concurrent jobs also finish at different times, providing new evidence while other computations remain in progress. These differences in computational cost and runtime motivate a supervisory campaign controller that continually interprets incoming evidence and decides which methods and settings to run next while other jobs continue asynchronously.

Here, we introduce **Target-adaptive Rescue–Explore–eXploit (T-REX)**, an agentic campaign controller for high-throughput de novo binder design. In T-REX, LLM agents interpret accumulating campaign evidence to propose and prioritize work according to three strategies: **Rescue**, which directs follow-up computation toward an identified weakness in a promising design or route; **Explore**, which tests underused methods or alternative configurations; and **eXploit**, which allocates additional computation to productive routes. A deterministic execution layer then validates the proposed jobs, enforces resource constraints, and manages asynchronous execution. Across seven de novo binder-design targets, T-REX achieved the highest throughput of structurally distinct hits on every target, outperforming a non-agentic tree-search controller by a geometric mean of 2.43-fold and the strongest generator-only baseline for each target by 2.48-fold under matched compute budgets. Analysis of the campaign records shows that T-REX uses different follow-up jobs in response to distinct failure modes and shifts its **Rescue-Explore-eXploit** priorities as campaign conditions change. These results support evidence-guided, adaptive orchestration as a practical approach to allocating compute across complementary protein-design methods during a design campaign. T-REX is available fully open source at https://github.com/ml-struct-bio/T-REX.

## 2. Results

### 2.1. Overview of the T-REX campaign controller

T-REX is designed to adaptively orchestrate diverse computational protein binder design tools to maximize structurally distinct hits under a fixed compute budget (Figure 1). The toolset includes six generation tools – four Proteína-Complexa variants (beam search, best-of-*n*, Feynman-Kac steering, and Monte Carlo Tree Search (MCTS)), BindCraft, and BoltzGen (Stark et al., 2025; Didi et al., 2026b; Pacesa et al., 2025) – along with ProteinMPNN (Dauparas et al., 2022) for sequence redesign and AlphaFold2 (AF2) refolding for structure evaluation (Jumper et al., 2021) (Figure 1a). We formulate a *de novo* binder-design campaign as an online allocation problem over computational jobs where each job specifies an action family with a particular tool, hyperparameter configuration, and, when applicable, a previously generated parent design (Supplementary Table S5). Designed binders must satisfy pre-specified *in silico* metrics to qualify as hits (e.g., pLDDT, iPAE, scRMSD thresholds), and the accumulating pool of designed binders is structurally clustered by Foldseek (Van Kempen et al., 2024) to count unique hits (Methods Section 4.2).

**Figure 1.**
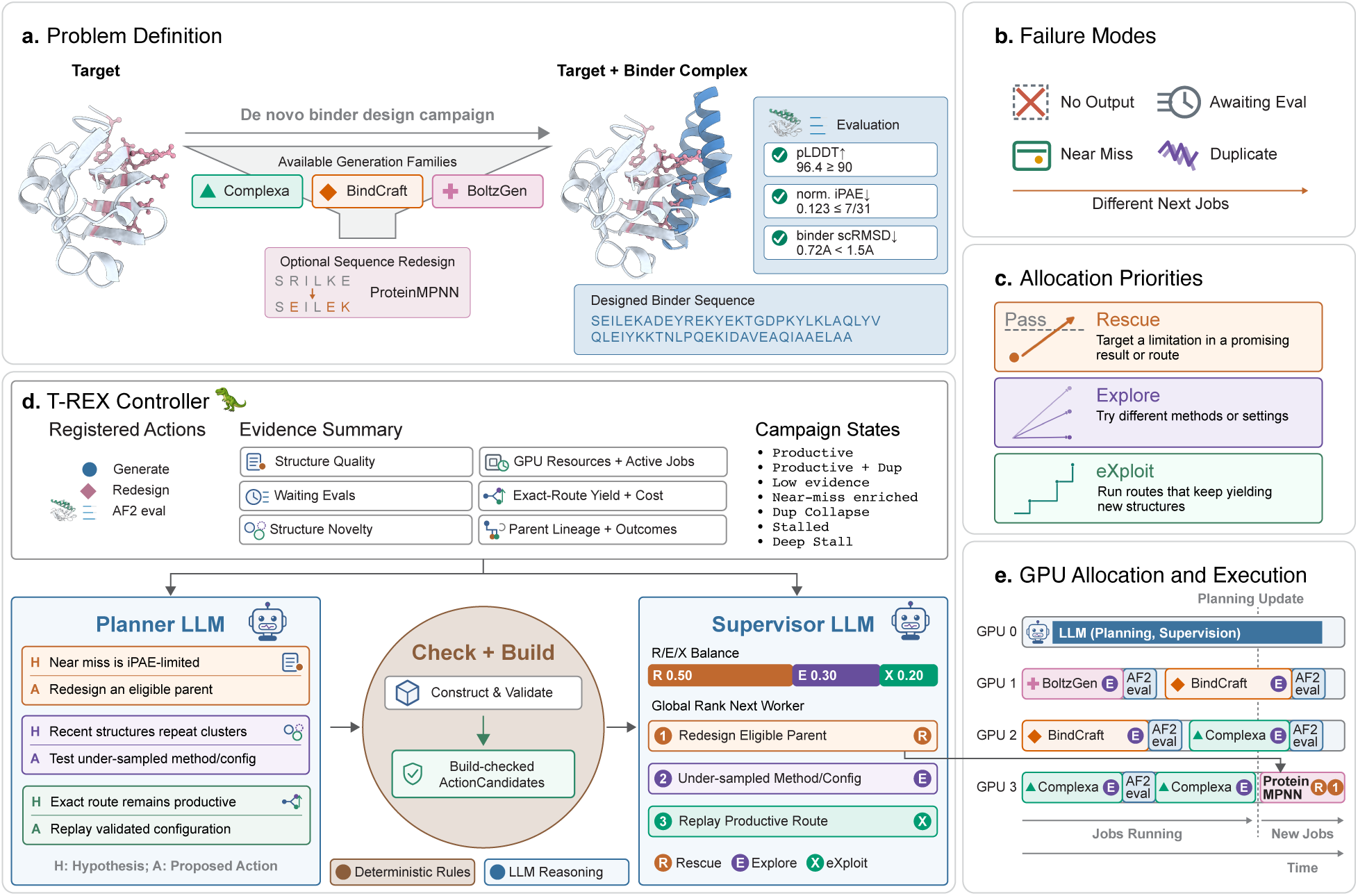
Overview of T-REX. **a.** T-REX is an agentic framework that adaptively orchestrates computational tools for *de novo* protein binder design under a fixed compute budget. **b,c.** Design outcomes are organized into failure modes that motivate distinct priorities for subsequent actions: **Rescue** promising designs, **Explore** alternative methods or settings, or **eXploit** productive routes. **d.** Evidence from completed and ongoing design jobs is organized into a structured representation of the evolving campaign state and interpreted by two LLM agents. The Planner proposes evidence-guided hypotheses and follow-up actions, while the Supervisor prioritizes validated actions and proposes their allocation across Rescue, Explore and eXploit. A deterministic controller orchestrates job execution given the available compute resources. **e.** Proposed actions are asynchronously executed across available GPUs, with completed results providing new evidence for subsequent planning and allocation.

During a campaign, hit production can slow or stall, but simply considering a scalar hit count does not reveal where progress is being lost or what type of follow-up is appropriate. In T-REX, we distinguish four operational failure modes (Figure 1b). These correspond to different points where ongoing or completed work has not yet produced a new structural hit: generation may produce no usable output, evaluation may still be pending, a candidate may narrowly miss the hit qualification criteria, or a qualified design may duplicate a structure already found. Because these failure modes arise for different reasons, they can require different responses. To organize these responses, T-REX defines three priorities for follow-up actions: **Rescue**, **Explore**, and **eXploit** (Figure 1c). **Rescue** addresses an observed limitation in a promising design or route, **Explore** tests alternative methods or settings, and **eXploit** allocates further compute to routes that continue to produce new structural hits.

Rather than relying only on a scalar hit count, T-REX uses a structured summary of outcomes (i.e. the *EvidenceSummary*) to represent the broader campaign context (Figure 1d). The *EvidenceSummary* combines completed results, ongoing work, and available resources, summarizing evidence about structure quality and novelty, route-specific yield and cost, pending evaluations, design lineage, and active compute (Figure 1d; Methods Section 4.3; Supplementary Section C.4 and Supplementary Figure S1). The *EvidenceSummary* is used to assign one of seven campaign states, which provide a high-level description of the campaign’s current condition, such as whether evidence is still sparse, new structures are being produced, near misses are accumulating, or progress has stalled (Figure 1d and Supplementary Section C.4.5). Because the relevance of the evidence depends on the current campaign state and history, choosing what to try next requires contextual interpretation. T-REX therefore uses LLM agents to reason over the campaign state, underlying evidence, and history. The Planner LLM formulates new evidence-linked hypotheses and follow-up tests as *HypothesisCards*. The proposed follow-up jobs then pass a deterministic “Check + Build” stage for validation. Finally, the Supervisor LLM assigns each valid candidate job to **Rescue**, **Explore**, or **eXploit**, ranks candidate jobs, and proposes an R/E/X allocation specifying the desired proportions of subsequent jobs devoted to each priority (Figure 1d). The deterministic controller uses these priorities and rankings to select jobs for execution as worker capacity becomes available, subject to feasibility and resource constraints.

Generation, redesign, and evaluation jobs can differ substantially in runtime and resource requirements, so waiting for all ongoing work to finish before planning the next job would delay the use of newly available evidence from a recently completed job. T-REX therefore uses asynchronous execution to incorporate completed results into decisions about which methods and settings to run next (Figure 1e). When a job finishes and its GPU becomes available, the controller can assign a new proposed job to that GPU while jobs on other GPUs continue. As each job finishes, its results are added to the campaign archive, a JSON-style record of campaign results and decisions (Methods Section 4.3). These results are summarized in the *EvidenceSummary*, which is used primarily as evidence for deciding which job to run next. This closes the loop between execution and evidence-guided planning (Figure 1e; Methods Section 4.5). In this work, the Planner and Supervisor use Qwen3.6-27B-FP8, served with vLLM (Team, 2026; Kwon et al., 2023).

### 2.2. T-REX achieves high-throughput in silico design across diverse protein targets

We compare T-REX against three generator-only baselines and two non-agentic adaptive controllers under matched worker-compute budgets across seven targets selected from the Proteína-Complexa (Complexa) (Didi et al., 2026a) benchmark: CD45, BetV1, CbAgo, HER2-AAV, SC2RBD, PDL1 and IL7RA (Figure 2; Table 1). The generator-only baselines were Complexa, BindCraft (Pacesa et al., 2025) and BoltzGen (Stark et al., 2025); the adaptive comparators were non-agentic tree-search controller PUCT (Aygün et al., 2026; Silver et al., 2017; Rosin, 2011) and *ε*-greedy (Vermorel and Mohri, 2005). PUCT and *ε*-greedy operated over the same action space and execution and evaluation framework as T-REX, but used scalar reward-based allocation without LLM-based contextual reasoning (Supplementary Section B.5.1).

**Figure 2.**
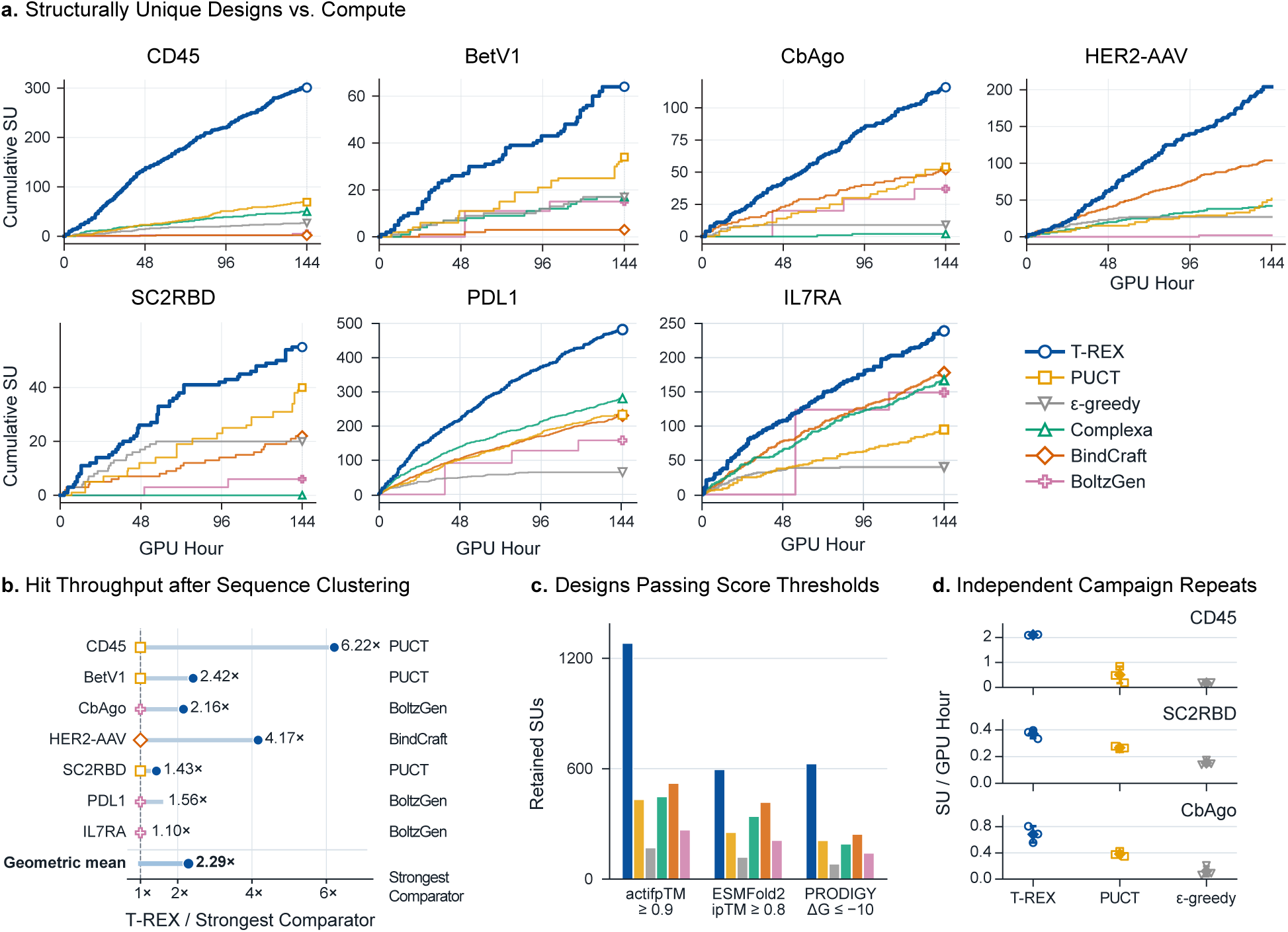
T-REX produces more structurally unique hits than baseline methods over the course of a design campaign. **a.** Cumulative number of structurally unique hits versus GPU-hours across seven targets for T-REX, non-agentic adaptive approaches and single-generator baselines. **b.** Hit throughput under MMseqs2 clustering at 70% sequence identity, shown relative to the highest-throughput alternative for each target. **c.** Number of structurally unique hits retained under alternative post-hoc quality thresholds, pooled across seven targets. actifpTM and ESMFold2 ipTM measure predicted interface confidence, and PRODIGY estimates binding free energy (Δ*G*, kcal mol*^−^*^1^). Each threshold is applied separately to the same set of TM0.6 cluster representatives. **d.** Hit throughput across three independently initialized campaigns for T-REX, PUCT and *ε*-greedy on CD45, SC2RBD and CbAgo. Points show individual campaigns; diamonds and error bars show the mean and s.d.

**Table 1.**
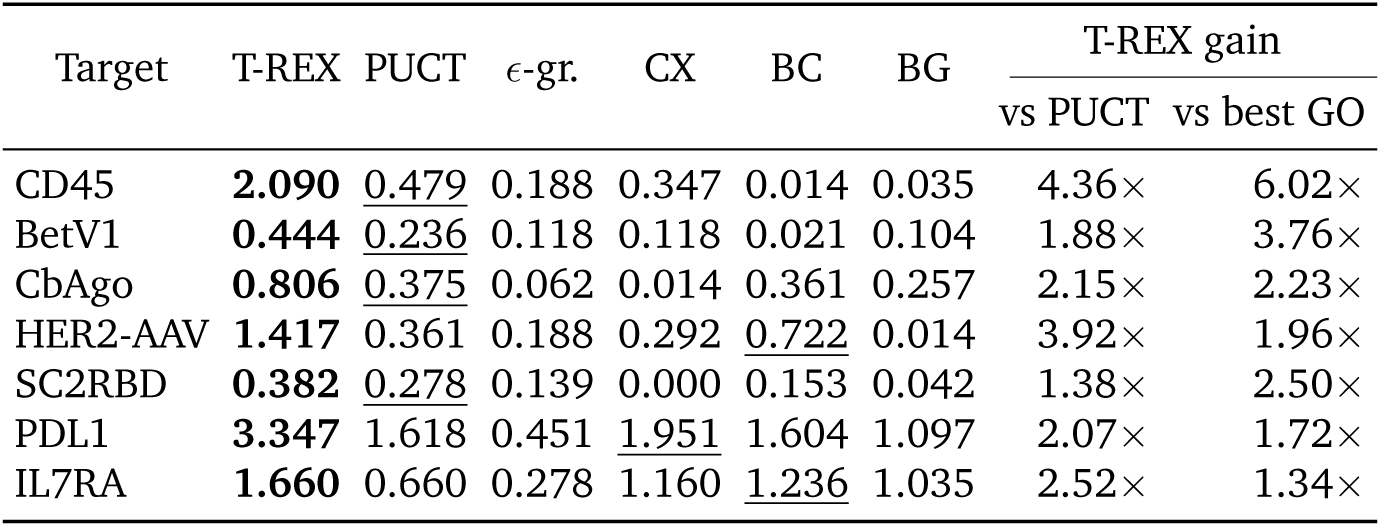
Protein binder design hit throughput for T-REX and baseline methods. Hit throughput is reported as the number of structurally unique hits per GPU-hour under a total 144 H100 worker GPU-hour budget. Hits are designs that pass the prespecified *in silico* criteria and are clustered with Foldseek at a TM-score threshold of 0.6. Gain columns report the T-REX ratio relative to PUCT and the highest-throughput generator-only (GO) baseline for each target. *ε*-gr., *ε*-greedy; CX, Complexa-only; BC, BindCraft-only; BG, BoltzGen-only. Bold and underline indicate the highest and second-highest throughput, respectively, for each target.

Across the seven targets, T-REX produced 1,461 structurally unique hits (SUs) clustered at a TM threshold of 0.6 with each design campaign spanning 48 hours on a 4-H100 GPU node (144 worker GPU-hours). The final SU count of T-REX for each target exceeded all five baselines (Figure 2a and Table 1). No single generator-only method was strongest across all targets, supporting the use of target-adaptive tool orchestration. T-REX throughput was 1.38–4.36-fold that of PUCT across targets, with geometric-mean gains of 2.43-fold over PUCT and 2.48-fold over the strongest observed generator-only baseline for each target (Figure 2a and Table 1). Comparisons at 10, 50, 100 and 144 GPU-hours further quantify the throughput trajectories in Figure 2a (Supplementary Section D.1.2 and Supplementary Table S10).

The throughput gain of T-REX was robust to alternative definitions of structural diversity and design quality (Figure 2b, c). T-REX led on every target under structural clustering at TM-score thresholds of 0.5, 0.6 and 0.8 and sequence clustering at 70% identity (Figure 2b and Supplementary Tables S14 and S13). The throughput gain was also retained as hit qualification criteria were tightened (Supplementary Figure S4). Across AF2, actifpTM, PRODIGY, ESMFold2 and Boltz-2 diagnostics, T-REX retained the largest total number of structurally distinct hits passing each score threshold (Figure 2c and Supplementary Figure S6). This indicates that the throughput gain of T-REX is not limited to AF2-based assessment.

We ran independently initialized replicate campaigns for three targets (CD45, SC2RBD, and CbAgo) and observed that T-REX consistently outperformed both PUCT and *ε*-greedy baselines (Figure 2d). T-REX also had higher throughput than a no-LLM SMAC3 Bayesian optimizer on all seven targets, extending the comparison beyond PUCT and *ε*-greedy (Supplementary Table S11) (Lindauer et al., 2022). Together, these analyses show that the throughput gain was not specific to the primary diversity or quality criteria, a single controller initialization, or the particular adaptive comparators used in the primary benchmark.

### 2.3. T-REX integrates campaign evidence to propose follow-up jobs

To qualitatively investigate how T-REX uses campaign context to interpret failures and select follow-up jobs, we retrospectively examined three illustrative campaign trajectories. In HER2-AAV, a pLDDT-reward test yielded no qualified designs (0/32), leading to a revised hypothesis about sequence–backbone compatibility and a sequence refinement job that yielded 5/32 qualified designs (Figure 3a). In CbAgo, five designs with iPAE values above the cutoff motivated broader Feynman–Kac (FK) steering (Singhal et al., 2025; Didi et al., 2026b), which yielded 4/16 qualified designs (Figure 3b). In PDL1, a parent that failed only pLDDT was selected for fixed-backbone ProteinMPNN redesign, yielding 14/16 qualified descendants and one new SU (Figure 3c). For each case, the campaign archive links the earlier evidence to the proposed follow-up test, confirmed job start, and resulting measurements; the PDL1 example additionally retains the direct parent–descendant relationship (Supplementary Section D.2.3). These examples illustrate how T-REX reused criterion- and route-specific evidence to propose different follow-up computations.

**Figure 3.**
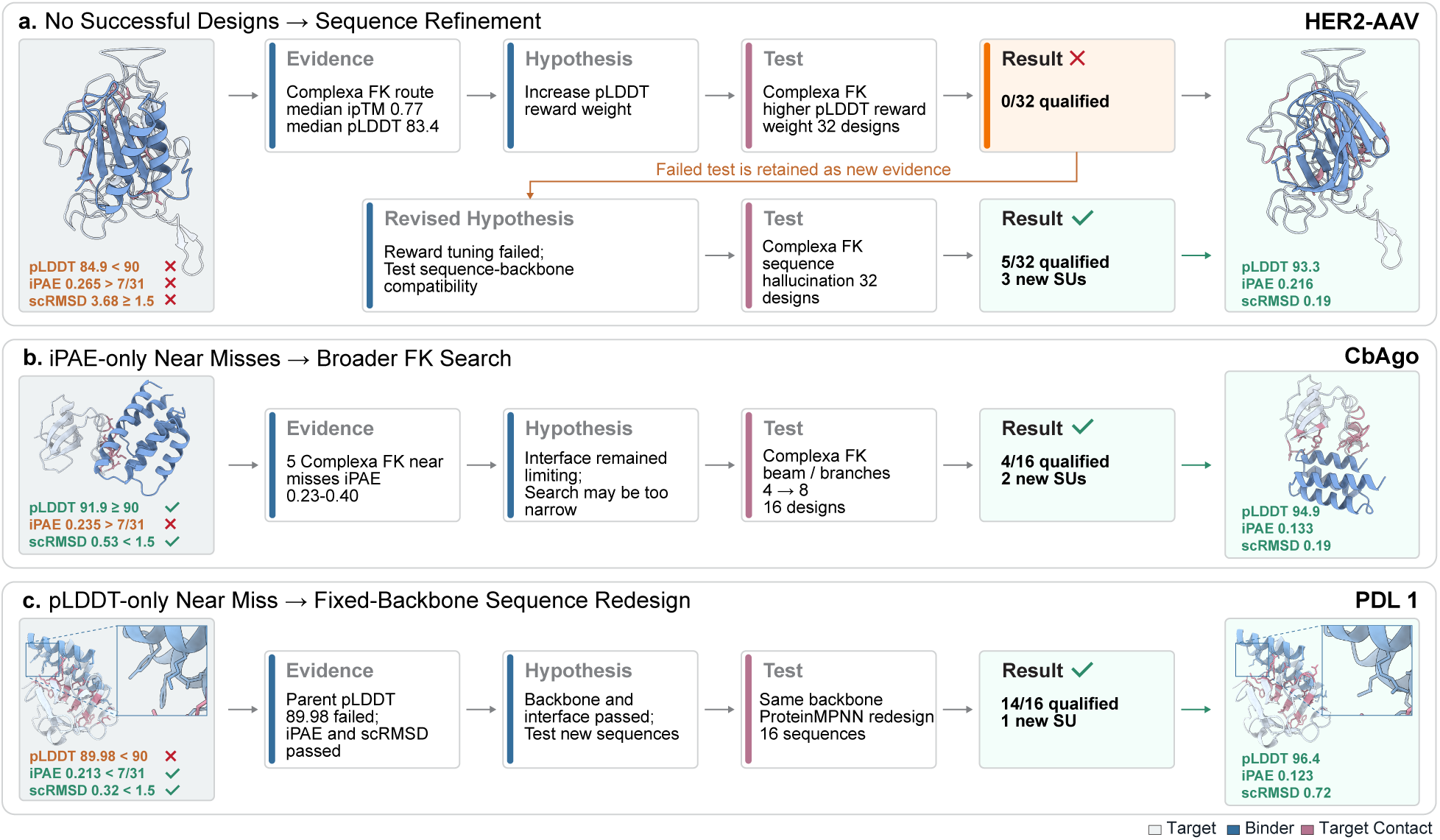
T-REX uses campaign outcomes to propose follow-up design strategies. Three retrospective examples illustrate how T-REX uses observed outcomes to propose subsequent computational actions. **a.** For HER2-AAV, a batch optimizing pLDDT produced no successful designs, motivating sequence refinement that produced 5/32 successful designs and three new structurally unique hits. **b.** For CbAgo, repeated designs failing only the iPAE criterion motivated broader Feynman–Kac (FK) search, producing 4/16 successful designs and two new structurally unique hits. **c.** For PDL1, a design failing only the pLDDT criterion motivated fixed-backbone ProteinMPNN sequence redesign, producing 14/16 successful designs and one new structurally unique hit. Orange and green labels indicate failed and passed design criteria, respectively.

### 2.4. T-REX adapts allocation across campaign states and prioritizes stall recovery

We next investigated how T-REX adapts its allocation by examining shifts in **Rescue**, **Explore**, and **eXploit** priorities across campaign states. Campaign states were assigned from archived campaign evidence using our seven-state classification (Figure 1d and Methods Section 4.3). Among 1,454 confirmed generation and redesign starts, **Explore** accounted for 74% of starts when little evidence was available, **Rescue** for 51% during “stall” or “deep stall”, and **eXploit** for 82% in “productive-with-duplication” states (Figure 4a). Thus, T-REX allocated work differently across campaign states, favoring **Explore** when evidence was sparse, **Rescue** during stalls and **eXploit** when productive routes continued to yield new structures despite frequent structural duplication.

**Figure 4.**
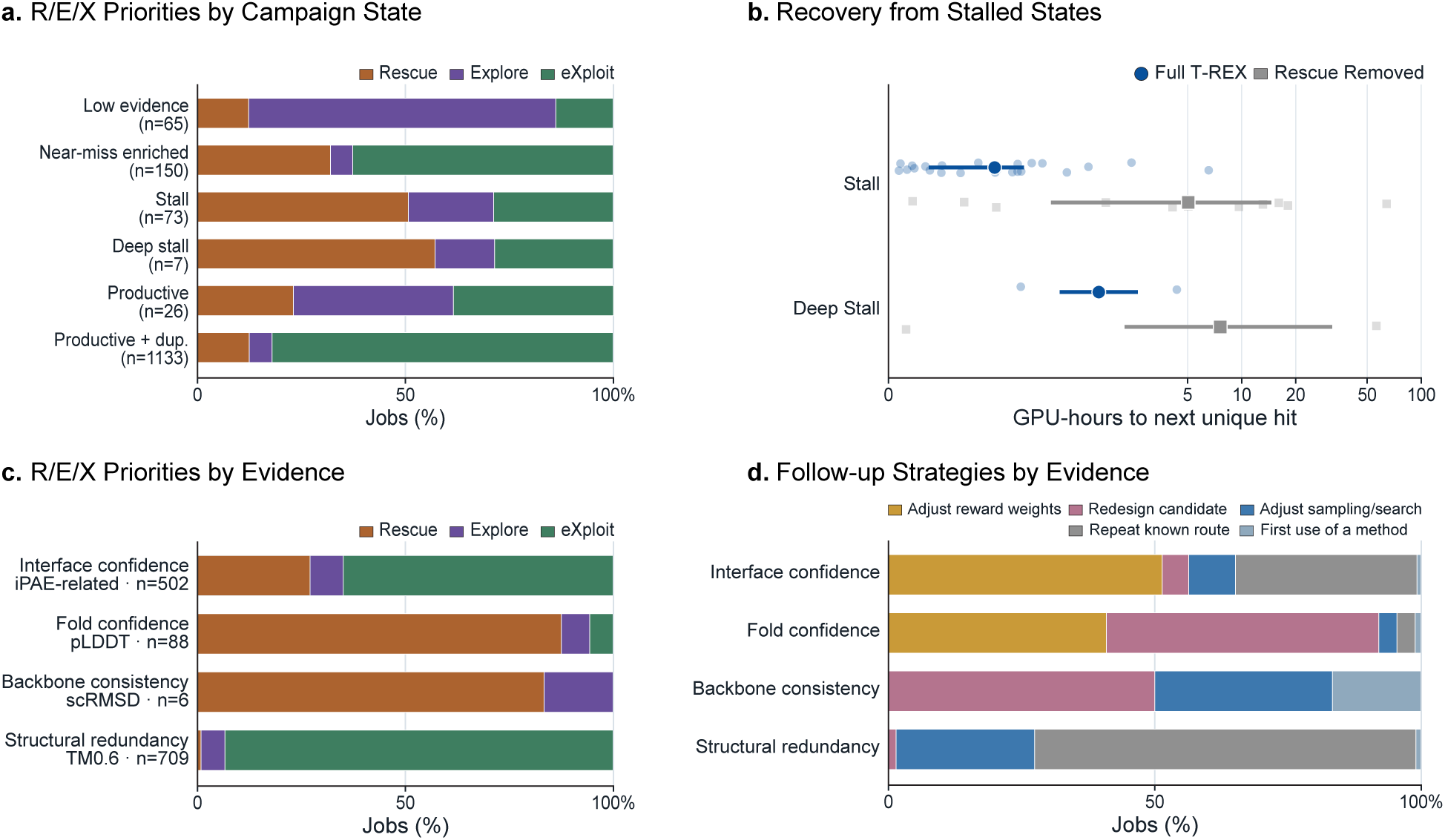
T-REX adapts design strategies to campaign state and accumulated evidence. **a.** Rescue– Explore–eXploit (R/E/X) priorities across 1,454 generation and redesign jobs, grouped by campaign state. **b.** GPU-hours from entry into a stalled campaign state to the next structurally unique hit for T-REX and T-REX without an explicit Rescue priority. Points show individual recovery events; larger markers and bars show medians and interquartile ranges. **c.** R/E/X priorities across 1,305 generation and redesign jobs, grouped by the primary evidence informing each decision. **d.** Follow-up design strategies for the same jobs, grouped by primary evidence as in **c**. compositions, including more *β*-rich clusters than the other methods tested.

We tested the role of **Rescue** prioritization in stall recovery by removing this priority while retaining the same action space. Over the first 100 GPU-hours, recovery was measured from entry into a stall or deep-stall campaign state until the next new TM 0.6 SU (Figure 4b; Supplementary Section D.3.3). Full T-REX recovered in 26 of 27 such entries, compared with 14 of 20 under the Rescue removed intervention. Among entries followed by recovery, the median time to the next SU was shorter with full T-REX for both “stall” (1.77 versus 5.04 GPU-hours) and “deep stall” (3.52 versus 7.60 GPU-hours); full T-REX also spent less time in “stall” or “deep-stall” states than the Rescue removed intervention (3.8% versus 27.7%). These results support a role for **Rescue** prioritization in recovering from stalled campaign states.

Finally, we examined how different job outcomes and structural duplication were associated with allocation priorities and follow-up jobs. Among 1,305 confirmed generation and redesign starts with a retrospectively identified primary evidence focus, interface confidence (iPAE) and structural duplication were most often associated with **eXploit** (65% and 93%, respectively), while fold confidence (pLDDT) was most often associated with **Rescue** (88%; Figure 4c). Follow-up jobs also differed by evidence: confidence-related evidence commonly led to changes in generator scoring weights or sequence redesign, while structural duplication was most often associated with repeating known routes, followed by changes in sampling or search settings (Figure 4d; Supplementary Section D.3.2).

### 2.5. Designed proteins span diverse secondary structures and predicted binding positions

To qualitatively characterize the repertoire of binder structures produced by T-REX, we examined the secondary-structure composition of the structurally distinct hits. Across the seven primary target campaigns, we calculated *α*-helix and *β*-strand fractions for all 1,461 final TM 0.6 representatives using simplified three-state Dictionary of Protein Secondary Structure (DSSP) assignments (Kabsch and Sander, 1983; McGibbon et al., 2015). Representative *α*-helix-rich, mixed helix/strand and *β*-strand-rich binders are shown in Figure 5a, with their secondary-structure composition shown among all representatives in Figure 5b. To compare *β*-strand-rich designs across methods, we selected all qualified designs with at least 20% of binder residues assigned as a *β*-strand before structural clustering. Across the seven targets, T-REX yielded 308 TM 0.6 clusters, compared with 95 for BoltzGen-only, the highest count among the other methods (Supplementary Table S23). These results show that structurally distinct hits produced by T-REX span different secondary-structure

**Figure 5.**
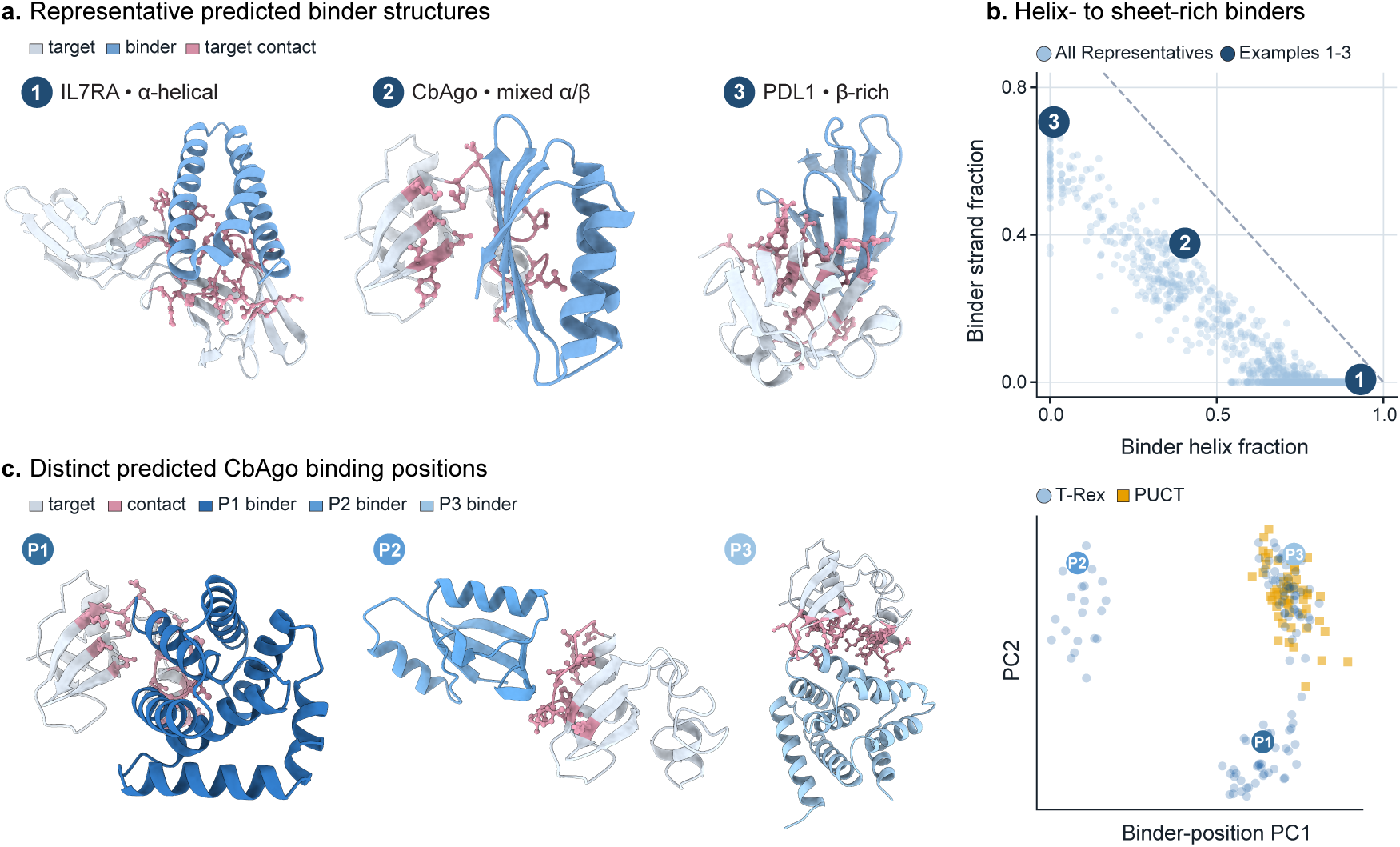
T-REX produces binders with diverse structures and predicted binding positions. **a.** Representative predicted complexes showing an *α*-helix-rich IL7RA binder, a mixed *α*-helix/*β*-strand CbAgo binder and a *β*-rich PDL1 binder. **b.** Secondary-structure fractions of T-REX hits (*n* = 1, 461), calculated using DSSP. Numbered points correspond to the examples in **a**. **c.** Representative T-REX hits at distinct predicted binding positions on CbAgo and principal component analysis (PCA) of target-relative binder positions for hits from T-REX (*n* = 116) and PUCT (*n* = 54).

We then analyzed predicted target-relative binder positions for CbAgo to assess structural variation around the target. After aligning each predicted complex on the target, we obtained each binder center as the mean position of its C*α* atoms. Joint principal component analysis (PCA) showed that T-REX binder centers occupied a broader region than those from PUCT (Figure 5c). Across 1,000 sample-size-matched resamples of 30 representatives per method, the median pairwise separation between binder centers averaged 31.2 Å for T-REX and 9.97 Å for PUCT. Related interface-contact analyses for all seven targets are reported in Supplementary Table S24. Together, these results show additional structural variation among T-REX hits, including different secondary-structure compositions and, for CbAgo, a broad range of predicted target-relative binder positions.

## 3. Discussion

We propose **Target-adaptive Rescue–Explore–eXploit (T-REX)**, an agentic campaign controller that achieves high-throughput *de novo* binder design by orchestrating multiple specialized protein structure generators and evaluators. By leveraging multiple design tools and strategies, T-REX achieved higher hit throughput than both single-generator baselines and non-agentic adaptive controllers under matched compute budgets (Figure 2). These results highlight how campaign-level orchestration is emerging as a complementary direction to continued improvements in specialized protein-design tools, consistent with general trends towards higher-level abstractions for generative design in biology (Merchant et al., 2026).

Central to T-REX is its combination of adaptive decision-making driven by LLM reasoning, a structured classification of outcomes and failure modes tailored to protein-design campaigns, and deterministic control for efficient resource allocation. Rather than asking an LLM to reason directly over an unstructured history of design attempts, T-REX organizes campaign outcomes into interpretable failure modes and campaign states that provide context for subsequent decisions (Figure 1). Two LLM agents then operate over this structured evidence: a Planner proposes evidence-linked hypotheses and follow-up actions, while a Supervisor prioritizes candidate actions and proposes how subsequent computation should be distributed among **Rescue**, **Explore**, and **eXploit**. These priorities distinguish between repairing promising but unsuccessful designs or routes, exploring alternative strategies, and extending strategies that are already productive (Figures 3,4). Together, these design choices provide structure to autonomous decision-making while retaining the flexibility of LLM reasoning.

Here, we solely evaluated T-REX in a computational setting using established *in silico* metrics to score “successful” and “unique” hits. While experimental validation is ultimately required, we note that the individual generators within T-REX (Proteína-Complexa, BindCraft and BoltzGen) have each produced experimentally validated binders in their original studies (Didi et al., 2026a; Pacesa et al., 2025; Stark et al., 2025). Real-world *de novo* design and development also require assessment of binding affinity (Passaro et al., 2025; Xue et al., 2016), specificity and developability (Listov et al., 2024), followed by iterative lead optimization, molecular simulation and mutation screening (Barletta et al., 2024; Chaudhury et al., 2010). Extending T-REX to these stages would require adding the corresponding computational tools and experimental operations to the action space and adapting the *EvidenceSummary*, campaign states, decision criteria, and deterministic constraints to their outcomes and costs. Emerging techniques in robotics and autonomous laboratories could further connect this control loop to experimental validation, enabling cycles of design, testing and refinement (Boiko et al., 2023).

More generally, T-REX complements recent scientific-agent systems, including *Autoresearch* and *The AI Scientist*, that use feedback to guide extended sequences of computational experiments (Karpathy, 2026; Lu et al., 2026; Yamada et al., 2025). A potentially transferable design principle is the separation of structured campaign-state representation, LLM-based proposal and prioritization, and deterministic validation and execution for allocating limited resources. Future work could examine whether these principles can improve resource allocation in other scientific campaigns involving heterogeneous tools, delayed outcomes, and fixed budgets, using domain-specific campaign states and Rescue, Explore, and eXploit priorities. To facilitate adoption and further development, we openly release T-REX together with its agent prompts and campaign-control framework. T-REX uses Qwen3.6-27B-FP8, an openly available LLM that can be served locally, and is available at https://github.com/ml-struct-bio/T-REX.

## 4. Methods

### 4.1. Campaign specification and shared action space

**Campaign setup and controllers.** T-REX adaptively selects computational jobs to maximize the number of structurally distinct binder-design hits obtained within a fixed compute budget. A campaign comprises generation, redesign, and evaluation jobs carried out for a single target using a fixed compute budget and multiple GPUs. For each target, campaigns used a prespecified target structure and binding-site hotspots, along with a default binder-length range (Supplementary Table S1). Structurally distinct hits were counted using the common in silico qualification criteria and structural clustering described in Section 4.2. T-REX, PUCT (Aygün et al., 2026; Silver et al., 2017; Rosin, 2011) and *ε*-greedy (Vermorel and Mohri, 2005) served as campaign-level controllers and operated over the same action space, with shared parameter bounds and a common execution and evaluation framework. PUCT and *ε*-greedy used scalar reward-based allocation without using LLM-based contextual reasoning (Supplementary Section B.5.1).

**Shared action space.** The shared action space, defined as the action families and configurations permitted within a campaign, comprised eight action families: de novo generation using four Complexa search variants—beam search, best-of-*n*, Feynman–Kac (FK) steering and Monte Carlo tree search (MCTS)—as well as BindCraft and BoltzGen; sequence redesign using ProteinMPNN; and evaluation using AF2 (Singhal et al., 2025; Didi et al., 2026b; Pacesa et al., 2025; Stark et al., 2025; Dauparas et al., 2022). Within this action space, an action family identifies a computational method, a route groups generation or redesign jobs that share an action family and configuration, and a job specifies the concrete computation to execute, including its permitted settings and, when applicable, an eligible parent design. Controllers could adjust family-specific configuration settings and the weights of Complexa reward terms or BindCraft design loss terms within prespecified bounds. The pretrained model weights, generator implementations, and permitted action space remained fixed throughout each campaign, while available jobs changed as controllers selected permitted configurations and newly generated designs became eligible for redesign.

**Campaign execution.** All three adaptive controllers began with prespecified Complexa Beam, BoltzGen and BindCraft jobs using common initial settings. PUCT and *ε*-greedy additionally started a Complexa best-of-*n* job on their fourth worker. As results became available, subsequent job selections used the accumulated campaign evidence, including job and evaluation outcomes, design quality and novelty, route productivity and cost, and resource state (Section 4.3, Supplementary Section C.4). Worker use was counted per job and qualification per design. For controller statistics, routes served as the unit for aggregating job outcomes and compute use. Redesign routes retained the upstream generation context. Operationally, each campaign followed a repeated evidence, planning, and execution loop: completed jobs updated campaign evidence; the controller used that evidence to propose and prioritize follow-up work; validated jobs were admitted to the shared worker queue; and their outcomes became evidence for subsequent decisions. Sections 4.3–4.5 describe each stage of this loop.

### 4.2. Design qualification and throughput measurement

**Design qualification.** Designs were evaluated in two stages: they first had to satisfy the common qualification criteria, after which qualified binder chains were clustered to identify structurally distinct hits (SUs). Designs qualified only if binder pLDDT ≥ 90, normalized iPAE ≤ 7/31 and binder scRMSD < 1.5 Å, using thresholds adapted from the Proteina-Complexa evaluation protocol (Didi et al., 2026b) and measurements from the common ColabDesign-derived AF2-Multimer protocol (Didi et al., 2026b; Evans et al., 2021; Ovchinnikov et al., 2025). Normalized iPAE was the mean of the two directional binder-to-target and target-to-binder PAE means, divided by 31. Binder scRMSD was the C*α* root-mean-square deviation between generated and AF2-predicted binder structures after Kabsch alignment of the corresponding target C*α* atoms (Kabsch, 1976). Complexa normally produced all three qualification measurements during generation. Outputs from BindCraft, BoltzGen, and ProteinMPNN redesign generally required separate standardized AF2 evaluation, subject to the admission rules described in Section 4.5. Generator-native scores (e.g., BoltzGen design ipTM, interface ipTM, structure confidence and native-refold RMSD) and labels (e.g., BindCraft accept/reject labels indicating whether a design passed BindCraft’s native filters) could guide planning but did not determine qualification.

**Structural novelty.** A qualified design could still add no new SU if its binder chain clustered with a structure already represented in the campaign. Foldseek clustered qualified binder chains separately within each campaign at TM0.6, a threshold previously used with Foldseek to quantify structural diversity in generated protein backbones (Jendrusch and Korbel, 2025). TM0.5 and TM0.8 were additionally evaluated as sensitivity analyses (Supplementary Table S14). Each final cluster counted as one structure-unique unit (SU) (Van Kempen et al., 2024), quantifying structural similarity and not biological fold or epitope identity. Protocol and clustering details are given in Supplementary Sections B.1–B.2 and Table S4.

**Throughput measurement.** Final SU counts for campaigns controlled adaptively by T-REX, PUCT or *ε*-greedy were recomputed from the finalized campaign records rather than copied from the last *EvidenceSummary* generated during campaign execution (see Section 4.3 for details on the *EvidenceSummary*). Each SU was credited to the generation or redesign job that produced its selected representative, not to an evaluation job; rules for selecting one representative design from each cluster are given in Supplementary Section B.2. Throughput is defined as *N*_SU_/*H*, where *N*_SU_ is the final cluster count and *H* = 144 H100 worker GPU-hours for each method–target combination in the primary benchmark. Adaptive campaigns counted generation, redesign and standardized AF2 evaluation toward this budget because these computations occurred during campaign execution and could inform subsequent allocation. For generator-only baselines, generation and search counted toward the budget, whereas standardized AF2 evaluation performed only after the generation cutoff was excluded because it could not affect which designs were generated. For T-REX, LLM serving and post-hoc analyses were excluded from *H*, which is defined to capture worker compute used during the design campaign. Alternative accounting and campaign-level statistics are specified in Supplementary Sections D.1.3 and D.1.1.

### 4.3. Campaign archive and evidence construction

**Campaign archive.** To make the evidence–planning–execution loop traceable, each campaign stored results and controller decisions in an archive that added new records without overwriting earlier ones. Three record types captured the main scientific results and planning information: (1) a *ResultRecord* recorded an output or failure from job execution; (2) an *EvidenceSummary* summarized accumulated evidence, unfinished work and available resources at each planning update; and (3) a *HypothesisCard* represented an evidence-linked hypothesis and proposed follow-up test. A *ResultRecord* linked each output or failure to its action family and route and, when applicable, to its parent design. It also stored available measurements, worker compute, execution status and links to structure, score and log files. A job could produce multiple *ResultRecords*, and individual outputs could be recorded before the job finished. If a job produced no usable design, its *ResultRecord* could still record the failure status and worker compute, even though no design measurements were available. In addition to these three record types, the archive stored candidate jobs, Supervisor rankings and allocation proportions, queue-admission decisions, confirmed worker starts, start failures, and LLM-call audits (Supplementary Section C.3).

**EvidenceSummary construction.** Before the Planner proposed follow-up tests and candidate jobs were selected, prespecified deterministic code constructed an *EvidenceSummary* from archived results, unfinished work and available resources. The *EvidenceSummary* provided the shared campaign context for the Planner and Supervisor; a reduced view of the summary was supplied to their prompts. As illustrated in Figure 1d, the summary organized evidence about (1) Structure Quality: qualification measurements, outcomes and margins; (2) Structure Novelty: new SUs and structural duplication; (3) Exact-Route Yield + Cost: route-level outcomes, recent new SUs per GPU-hour and compute use; (4) Waiting Evals: designs awaiting standardized AF2 evaluation; (5) Parent Lineage + Outcomes: parent designs and the outcomes of their descendants; and (6) GPU Resources + Active Jobs: queued and running jobs and available worker capacity. It also retained near-miss evidence and source-specific auxiliary measurements. A near miss had complete qualification measurements, failed qualification and satisfied the prespecified near-miss rule. Auxiliary measurements were source-specific diagnostic scores not used to determine qualification, including ipTM, ipSAE, interface energy and shape complementarity. The summary kept these measurements associated with their sources, and missing values were not treated as failures or zeros. The full *EvidenceSummary* schema, recent-evidence windows defined by *ResultRecord* count or recorded worker compute, and quality calculations are specified in Supplementary Sections C.4.1–C.4.3; the reduction applied before LLM input is described in Supplementary Section C.5.2.

**Campaign-state assignment.** Campaign-state labels provided context for LLM reasoning and deterministic allocation rules without prescribing a particular job (Figure 1d). As part of this evidence-construction step, a deterministic classifier assigned one of seven campaign-state labels: *low evidence*, *structural duplicate collapse*, *productive*, *productive with duplication*, *near-miss-enriched*, *stalled* or *deep stall*. The classifier used evidence of productivity, structural duplication, near misses and worker compute without a new SU. When the evidence satisfied the conditions for more than one campaign state, a prespecified decision order determined which label was assigned; *low evidence* served as the default when no condition matched. The assigned label was included in the *EvidenceSummary*. State definitions and thresholds are given in Supplementary Section C.4.5.

### 4.4. Evidence-guided planning and hypothesis feedback

**LLM agent roles.** T-REX used two LLM-based controller roles, the **Planner** and **Supervisor** (Figure 1d). Both were instantiated using the same locally served Qwen3.6-27B-FP8 model with vLLM, but used separate prompts and structured output schemas (Team, 2026; Kwon et al., 2023). We selected this open-weight model to enable local deployment with a fixed, versioned model snapshot and prespecified serving configuration. The model was served on a single H100 GPU. Across the seven primary campaigns, 7 of 3,125 LLM calls (0.22%) produced responses that failed structured-output validation; these responses were rejected before launch approval and triggered deterministic fallback paths (Supplementary Table S15). Functionally, the Planner proposed what to test next and why, whereas the Supervisor prioritized which validated candidates should receive future worker starts. Neither LLM directly launched jobs; candidate construction, validation, admission, dispatch, and execution were handled by deterministic controller logic (Section 4.5; Supplementary Section C.8.2).

**LLM decision guidance.** Both prompts provided guidance on whether follow-up work should extend a productive route, address an identified weakness, or test an alternative route or configuration. Route productivity was measured as the recent number of new TM0.6 SUs per GPU-hour charged to the route, including required evaluation and ancestor jobs. To determine which routes to prioritize for further computation, the prompts instructed both LLM roles to use recent route productivity as the primary quantitative criterion. Two routes were considered to have comparable productivity when the recent new TM0.6 SUs per GPU-hour of the less productive route was at least 85% of that of the more productive route, although both prompts advised caution when this comparison was based on few observations or little compute. For such routes, the LLM roles were instructed to prefer the route with the higher 25th percentile of its qualified-design quality scores, using the median as the next comparison (Supplementary Sections C.5.3, C.7 and C.4.3).

**Planning.** The Planner received the *EvidenceSummary* and campaign-state context, together with up to eight active or recent *HypothesisCards*, available action families and their roles, and permitted settings and per-job workload limits. The prompt requested one to four new *HypothesisCards*, each describing an evidence-linked hypothesis and a proposed follow-up computational test. Each new card cited supporting evidence, suggested action families and settings, and indicated the test’s relevance to Rescue, Explore, and eXploit. It also predicted how selected qualification measurements should change and, when applicable, identified other measurements that should not deteriorate beyond a specified limit. Serving settings, prompt details, and call conditions are specified in Supplementary Sections C.5 and C.5.4.

**Check + Build and candidate validation.** Planner proposals were not directly executable. Fixed Check + Build logic converted new or retained *HypothesisCards* into concrete *ActionCandidates* and validated the requested operation, registered action family, permitted parameter and workload bounds, required input files and, where applicable, parent eligibility. Permitted deterministic repairs were applied when defined; otherwise invalid candidates were rejected. This validation enforced the fixed action space, preventing the LLM from introducing unregistered action families, exceeding prespecified parameter bounds, or altering fixed campaign definitions such as the target, qualification criteria, structural-novelty rule, or worker budget. Only build-checked candidates were eligible for Supervisor prioritization and subsequent admission. Passing Check + Build established construction-time validity but did not guarantee execution; candidates remained subject to final admission rules and live dispatch-time checks described in Section 4.5.

**Supervisor prioritization and allocation.** The Supervisor received the *EvidenceSummary* and campaign-state context, together with hypotheses from the Planner and eligible candidate jobs. It recommended candidate jobs for execution, assigning each recommended job to **Rescue**, **Explore** or **eXploit** and ranking them across all three priorities. It also returned a normalized **R/E/X** mixture specifying the desired proportions of jobs to be executed under each priority. Detailed input and output specifications, ranking guidance and allocation rules are given in Supplementary Sections C.5.2 and C.7.

**Hypothesis feedback.** After selecting candidate jobs, the controller used prespecified rules to evaluate results from follow-up tests proposed earlier by the Planner. Results linked to each *HypothesisCard* were compared with a previously observed reference result and classified as supporting, contradictory or neutral. Accumulated results determined whether a card was supported, refuted or no longer active after its specified lifetime. These statuses informed later Planner and Supervisor decisions but did not affect design qualification or SU credit or validate the proposed causal explanation. Detailed evaluation and status-update rules are given in Supplementary Section C.9, with a worked example in Supplementary Section D.2.2.

### 4.5. Asynchronous job allocation and execution

**Candidate job selection.** Candidate selection operated on build-checked jobs described in Section 4.4. The Supervisor assigned each recommended candidate job to **Rescue**, **Explore** or **eXploit** and ranked them across the three priorities. The controller considered these jobs in rank order and used the **R/E/X** mixture to guide how subsequent generation and redesign starts were distributed, subject to prespecified feasibility and resource constraints. If additional jobs could be admitted to the shared worker queue, the controller selected them from the remaining eligible candidates using deterministic rules based on recent starts by action family, route productivity and estimated job cost. Selected jobs entered the shared worker queue, but only confirmed generation and redesign starts contributed to the realized **R/E/X allocation**. Detailed candidate ordering, allocation, fallback and dispatch rules are given in Supplementary Section C.7 and Table S8.

**Worker scheduling and asynchronous execution.** T-REX used three H100 workers and a separate H100 for LLM serving; PUCT and *ε*-greedy used four workers. T-REX had a reporting budget of 144 worker GPU-hours, corresponding to three worker slots over 48 h (3 × 48 = 144 worker GPU-hours). When a worker became free, the controller rechecked the next job’s inputs, parent eligibility and resource availability before starting or deferring it. Completed results were incorporated into campaign evidence as soon as they became available, so subsequent planning and job selection could use them while other jobs remained in progress. Detailed rules for queue admission, worker starts, concurrency limits and failure handling are given in Supplementary Section C.8, with the controller workflow summarized in Algorithm S1.

**AF2 evaluation admission.** Standardized AF2 evaluation used the same three worker GPUs as generation and redesign but followed a separate deterministic admission policy. Designs with usable structures were eligible for this evaluation when one or more of the three required qualification measurements had not been computed during generation (Figure 1e). Prespecified deterministic controller rules determined which eligible designs were evaluated and in what order. For each route requiring AF2 evaluation, up to four designs could initially be evaluated. After results from all evaluations within the current allowance were available, the cumulative allowance could increase successively to 8, 16, 32 and 64 evaluations and then continue doubling, but only if the route showed evidence of improvement relative to the preceding allowance boundary. Improvement was defined as an increase in route-attributed new SUs, near-miss count, the number of qualified structural clusters, the 25th percentile or median of the qualified-design quality scores, or the source-specific auxiliary diagnostic score. Qualified-design quality scores were calculated from the three normalized pass margins while retaining only the highest-scoring design per structural cluster. Exact score definitions, comparison rules, capacity limits and the spare-capacity exception are specified in Supplementary Sections C.4.3 and C.8.1.

**Campaign termination and final endpoint reconstruction.** The evidence–planning–execution loop ran until the prespecified 48-h campaign time limit; campaigns did not terminate based on campaign state, lack of new SUs or the number of SUs already obtained. The controller checked the time limit before each pass through the loop; a pass already in progress was allowed to finish. At shutdown, each running job was allowed up to 600 s to complete. Results from completed jobs were parsed. After execution stopped, the qualification criteria were reapplied to the final archived *ResultRecord* entries for outputs with a usable structure and all three required qualification measurements, including any such outputs recovered during shutdown. Foldseek TM0.6 clustering was then recomputed for the designs that qualified. Detailed shutdown, endpoint-inclusion and clustering rules are given in Supplementary Sections C.8.3, B.3 and B.2.

## Data availability

Design target inputs are listed in Supplementary Table S1. HER2-AAV, SC2RBD, PDL1 and IL7RA derive from Protein Data Bank (PDB) entries 1N8Z, 6M0J, 5O45 and 3DI3, respectively, with cropping or repacking recorded in Supplementary Table S1 (Cho et al., 2003; Lan et al., 2020; Magiera-Mularz et al., 2017; McElroy et al., 2009).

## Code availability

The T-REX source code is available at https://github.com/ml-struct-bio/T-REX. The repository includes installation instructions, example configurations and tools for running campaigns and analyzing campaign archives.

## Acknowledgments

The authors acknowledge the use of computing resources at Princeton Research Computing, a consortium of groups led by the Princeton Institute for Computational Science and Engineering (PICSciE) and Office of Information Technology’s Research Computing. E.D.Z. is grateful for support from the Chan Zuckerberg Initiative DAF (grant number 2025-358484), an advised fund of Silicon Valley Community Foundation, the National Institutes of Health (grant number DP2GM164606), the AI2050 program at Schmidt Sciences (grant number G-25-69788), the Princeton Catalysis Initiative, Princeton School of Engineering and Applied Sciences, Janssen Pharmaceuticals, and Generate Biomedicines.

Competing interests

E.D.Z. holds equity in Generate Biomedicines and serves on the Scientific Advisory Board of Deep Apple Therapeutics and the Chan Zuckerberg Imaging Institute.

## Supplementary Information

**Reading guide.** Section A specifies target inputs. Section B defines evaluation and benchmark comparisons. Section C describes the controller from stored observations through evidence, proposals, selection, execution and feedback. Section D groups supporting analyses for main Figures 2–5, including archived decisions and post-hoc characterization analyses. Monospace text denotes schema fields, settings or explicitly labeled example identifiers.

### A. Data and target specifications

#### A.1. Primary benchmark inputs

Unless otherwise stated, qualification, outcome and post-hoc diagnostic tables use the fixed seven-target set: CD45, BetV1, CbAgo, HER2-AAV, SC2RBD, PDL1 and IL7RA. These targets were selected from the benchmark assembled by Proteina-Complexa, and we retained its target structures, chain and residue selections, hotspot definitions and default binder-length ranges (Didi et al., 2026b). Table S1 lists the exact frozen specifications used here.

**Table S1.**
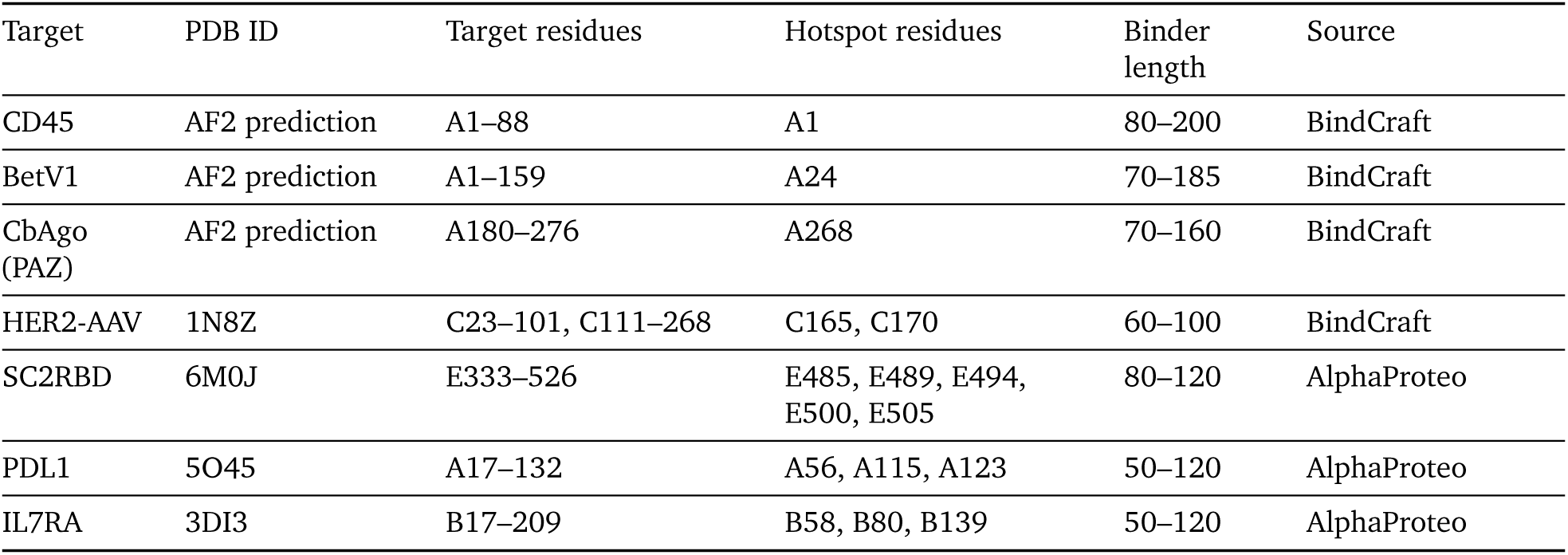
Fixed target inputs and default binder-length ranges. The table shows the frozen target specifications used in the primary benchmark. Entries without a PDB accession are identified as AF2 predictions. Residue and hotspot numbering follows each input structure. The binder-length column gives the target-specific default range; adaptive-controller BindCraft actions could vary these design bounds within the prespecified range without changing the target definition. Source denotes the benchmark from which the specification originated.

### B. Evaluation metrics and benchmark design

#### B.1. Qualification criteria and evaluation

Qualification required binder pLDDT ≥ 90, normalized iPAE ≤ 7/31 and binder scRMSD < 1.5 Å under the common AF2 protocol, with thresholds adapted from Proteina-Complexa (Didi et al., 2026b). pLDDT is the mean confidence over binder residues in the predicted complex, reported on a 0–100 scale (90 equals 0.90 on the 0–1 scale). Normalized iPAE averages the two directional binder–target PAE means and divides by 31 Å; the dimensionless cutoff 7/31 corresponds to an unnormalized mean of 7 Å. Binder scRMSD compares generated and AF2-predicted binder C*α* coordinates after Kabsch alignment of the corresponding target C*α* coordinates.

AlphaFold2 evaluation (Jumper et al., 2021; Evans et al., 2021) used model_1_multimer_v3, three recycles, no dropout, generated-complex initial coordinates and a binder template with inter-chain template contacts removed. Adaptive evaluation derived its seed from the first eight hexadecimal digits of SHA-256 over candidate_id|af2_refilter|s{RUN_SEED}, interpreted as an integer modulo 10^6^; generator-only evaluation used rank-index seeds. Complexa supplied the common AF2 measurements during generation when available, whereas outputs without a complete set required separate standardized AF2 evaluation. Harmonized AF2 was a separate post-hoc analysis and did not affect allocation, qualification or SU credit.

##### B.1.1. Outputs admitted to evaluation

An individual design was eligible for standardized AF2 evaluation when its output was recorded as successful and its structure was available in Protein Data Bank (PDB), Crystallographic Information File (CIF) or macromolecular CIF (mmCIF) format. Evaluation could begin while the producing job was still running (Section C.8.2). The structure could be supplied directly or retrieved from its output directory.

**Complexa and BindCraft outputs.** A final sample from Complexa (Didi et al., 2026b) was admitted when a valid structure was available. Inline measurements from the common qualification protocol were used when present; otherwise the structure entered standardized AF2 evaluation. Final PDBs from BindCraft (Pacesa et al., 2025) were eligible for standardized AF2 evaluation regardless of whether they passed BindCraft’s own filters.

**BoltzGen outputs.** For BoltzGen (Stark et al., 2025), the fixed reference admitted individual designs listed in BoltzGen’s metrics table and excluded aggregate root structures. CIF files were converted to PDB, and up to 10,000 designs per target were admitted in ascending order of BoltzGen’s final rank before qualification outcomes were inspected. Adaptive BoltzGen admitted completed individual designs with resolvable per-design structures, again excluding aggregate root structures. BoltzGen’s own measurements could inform allocation, but qualification required all three measurements from the common AF2 protocol. Only a subset of adaptive BoltzGen designs underwent this evaluation during the campaign.

**ProteinMPNN outputs.** For ProteinMPNN (Dauparas et al., 2022), a redesigned child was admitted when its generated sequence, from a FASTA-formatted row beginning with sample=, had been successfully threaded. The child entered standardized AF2 evaluation as a new design; evaluating its parent did not establish the child’s qualification.

#### B.2. Sequence and structural clustering

Foldseek clustered qualified binder chains, not full complexes. Sorted inputs were processed with easy-cluster, minimum sequence identity 0, minimum coverage 0, coverage mode 0, alignment type 1, alignment-length TM-score normalization (–tmscore-threshold-mode 0) and one thread. TM0.5 and TM0.8 changed only the primary TM0.6 threshold.

Representative-based set-cover clustering defines similarity to cluster representatives, not an all-pairs criterion or biological fold classes; higher thresholds generally give finer partitions (Van Kempen et al., 2024; Xu and Zhang, 2010; Jendrusch and Korbel, 2025; Liu et al., 2026). Sensitivity analyses re-clustered the same qualified pools without rerunning controllers. Native generator clusters did not determine cross-method SU credit.

MMseqs2 clustered qualified binder sequences before structural deduplication at 70% identity, 0.8 coverage and coverage mode 0 (Steinegger and Söding, 2017). Tables S13 and S14 report sequence throughput and structural-threshold sensitivity, respectively.

**Fixed representatives.** For every qualified design, normalized qualification margins were

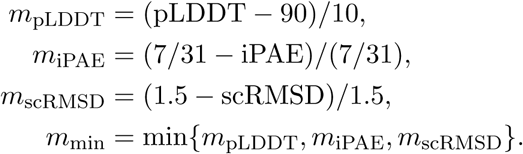

Each Foldseek TM0.6 cluster contributed the representative with the largest *m*_min_; ties were broken, in order, by larger mean margin, larger pLDDT margin, larger iPAE margin, larger scRMSD margin and finally a stable record identifier (lexicographically smaller for Complexa-only and larger for the other methods). These fixed representatives were used for final generation/redesign attribution, post-hoc analyses and qualification-margin sensitivity. During campaigns, the controller used separate route-specific representatives and quality summaries, defined in Section C.4.3.

#### B.3. Compute budget and endpoint inclusion

The primary benchmark used a fixed reporting denominator of 144 H100 worker GPU-h per method–target pair. T-REX used three worker GPUs plus one GPU for LLM serving; PUCT and *ε*-greedy used four workers. LLM serving and post-hoc diagnostic predictions were excluded from this denominator. Adaptive campaigns charged generation, redesign and standardized AF2 evaluation to the worker budget. For generator-only baselines, separate AF2 evaluation was performed post hoc after the 144 worker GPU-h generation budget. The comparison therefore matched worker compute rather than wall-clock duration. Final adaptive counts included eligible outputs collected during shutdown, without adding shutdown time to the reporting denominator. Generator-only counts used the admitted generation cohort defined in Section B.1.1. Table S2 summarizes execution differences. Compute-accounting sensitivities are reported in Section D.1.3; runtime and shutdown procedures are specified in Section C.8.3.

**Table S2.**
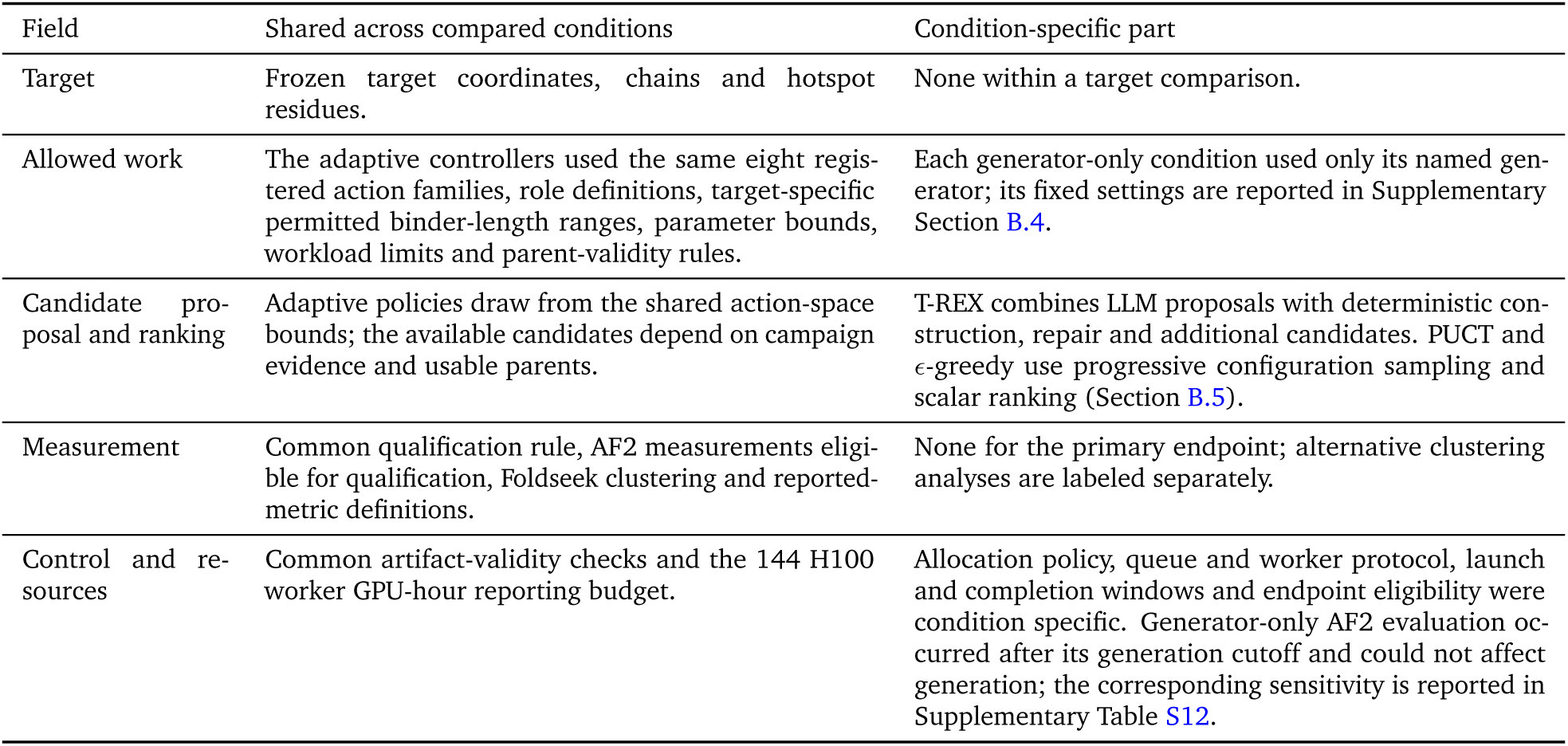
Shared and condition-specific fields in the primary benchmark. This table separates the common scientific task and endpoint from the allocation policy and its execution protocol.

#### B.4. Baseline setup and adaptive initialization

Each generator-only baseline repeatedly ran one prespecified workflow under the common 144 worker GPU-h generation budget: Proteina-Complexa’s Beam workflow, BindCraft’s default four-stage multimer workflow or BoltzGen’s published protein-anything recipe. Configurations were fixed across targets apart from the target inputs and binder-length intervals, and none was selected using T-REX outcomes. The adaptive controllers used the same generator implementations and learned checkpoints. Table S4 lists the versions, baseline settings and initial adaptive workloads; Table S1 specifies target inputs.

Adaptive campaigns began with prespecified Complexa Beam, BoltzGen and BindCraft jobs to obtain initial evidence. The initial Complexa settings matched Complexa-only; BindCraft and BoltzGen used smaller initial workloads to obtain earlier feedback, as specified in Table S4. Subsequent jobs varied within the bounds in Table S5. Successive Complexa runs used non-overlapping explicit seeds, BindCraft trajectories drew fresh stochastic seeds, and fixed BoltzGen runs were initialized from system entropy. Campaign replication is described in Section B.7.

**Development scope.** Controller settings were fixed rather than fitted online. Development runs used to calibrate the threshold for worker time without a new SU included some primary-panel targets. Here, target-independent means that the same runtime rules apply without target-identity branches, not that development used a disjoint held-out target set. The campaign-state definitions are given in Section C.4.5.

#### B.5. Adaptive comparator policies

The primary adaptive comparison shares the action-space bounds, scientific endpoint and required-evaluation machinery, while candidate proposal and ranking differ. T-REX combines LLM proposals with deterministic candidate construction, repair and supplementation (Section C.6); PUCT and *ε*-greedy use progressive configuration sampling and scalar ranking. Their shared AF2-admission policy can use diagnostic evidence (Section C.8.1) even though that evidence does not enter scalar ranking. These comparisons evaluate the complete allocation policies and do not isolate the contribution of LLM reasoning from the other policy differences.

##### B.5.1. Adaptive scalar-reward comparators

**PUCT.** Our PUCT comparator searched the same action space as T-REX without an LLM, using a prior-weighted exploration score adapted from tree-search allocation (Rosin, 2011; Silver et al., 2017; Aygün et al., 2026). It selected an action family and then a configuration or eligible parent-bound action within that family. At decision step *k*, the scalar reward was online new TM0.6 SU per charged worker GPU-h:

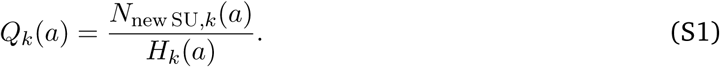

Here, *N*_new SU,*k*_(*a*) and *H_k_*(*a*) are the cumulative new-SU count credited to action *a* and its charged worker GPU-hours, with *Q_k_*(*a*) = 0 before any cost is recorded. Family rewards use the corresponding totals across their actions. Candidate configurations were progressively sampled within the shared parameter bounds.

At each level, the highest-scoring available child *a* of parent *p* was selected using

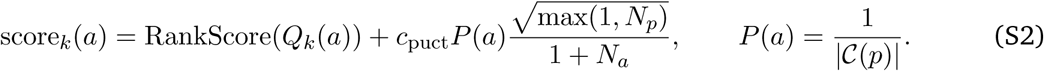

Here, C(*p*) is the available child set, *c*_puct_ = 1.0, and *N_p_*, *N_a_* are visit counts including queued, running and newly assigned work as virtual visits. For *m* distinct sibling rewards, the *j*th value in ascending order has RankScore = (*j* − 1)/(*m* − 1); it is zero when *m* = 1. This normalization puts the reward term on a 0–1 scale, while the second term favors less-visited actions.

*ε***-greedy.** The *ε*-greedy comparator used the same action space, configuration-sampling procedure and scalar reward *Q_k_*(*a*) in Eq. (S1), without an LLM (Vermorel and Mohri, 2005). It first prioritized unvisited feasible actions. Once none remained, it selected uniformly among available actions with probability *ε* = 0.10 and otherwise selected the action with the highest *Q_k_*(*a*). Both comparators used the shared validation, required AF2 evaluation, worker execution and qualification procedures.

##### B.5.2. Bayesian-optimization comparator

As an additional no-LLM comparator, SMAC3 searched the same action families and parameter bounds using a random-forest surrogate and expected-improvement acquisition (Lindauer et al., 2022). The objective was negative cumulative online new TM0.6 SU per route worker GPU-h. Initial sampling prioritized unobserved families and configurations; model fitting began after at least eight completed configuration observations with two distinct objective values.

The surrogate used action family and configuration, without scientific diagnostic fields or parent identity. Its acquisition score ranked feasible candidate jobs under the shared execution and qualification procedures. One campaign per target used four H100 workers for 25 h, with a reporting denominator of 100 worker GPU-h. Software versions and surrogate settings are listed in Table S4. The comparison with retrospective 100-worker-GPU-h prefixes is reported in Section D.1.2 and Table S11.

#### B.6. Related work: protein-design workflows and campaign control

**Protein-design workflows and adaptive execution.** ProteinDJ provides modular, parallel protein-design workflows and parameter sweeps (Silke et al., 2026); BinderFlow combines parallel binder-design batches with monitoring, hit-count-based stopping and candidate refinement (González-Rodríguez et al., 2025). Locuaz uses binding-affinity scores from MD-sampled complexes to retain or prune mutation lineages (Barletta et al., 2024), and BioPipelines demonstrates iterative redesign of small-molecule binding pockets using Boltz-2 predictions (Quargnali and Rivera-Fuentes, 2026). IMPRESS couples result-dependent ProteinMPNN–AlphaFold refinement with asynchronous execution and dynamic resource management (Alsaadi et al., 2025); Colmena provides more general infrastructure for result-guided scientific computation (Ward et al., 2025). These studies establish relevant precedents for workflow integration, feedback-directed optimization and adaptive execution. Our question concerns how to choose subsequent generation and redesign jobs across complementary methods as evidence accumulates within a finite-compute campaign.

**Scientific agents and binder-design campaigns.** ProtAgents combines LLM planning and critique with protein generation and physics-based analysis (Ghafarollahi and Buehler, 2024). The Virtual Lab develops a workflow for mutating existing nanobodies, with the resulting designs experimentally tested by human researchers (Swanson et al., 2025). ProteinMCP demonstrates autonomous protein-engineering workflows, including binder and nanobody design (Xu et al., 2026), while PDAgent uses template retrieval and conservation-aware directed mutation (Ouyang et al., 2026). The AutoBinder Agent preprint describes binder-workflow orchestration through the Model Context Protocol (MCP) and reports prompt-based evaluations of tool selection and execution order (Ge et al., 2026). The Claude binder-design technical report is a close precedent: agents choose methods and settings, manage GPU work within resource limits and deliver ranked designs for experimental testing (Claude Science and Shanehsazzadeh, 2026). It establishes that budget-aware multi-tool binder campaigns already form part of agentic protein design.

**Campaign control and evaluation in T-REX.** T-REX combines criterion-specific qualification outcomes, structural redundancy, computational cost and pending evaluations to guide evidence-linked follow-up tests and **Rescue**, **Explore** and **eXploit** priorities. LLM agents propose and prioritize work; a deterministic execution layer constructs and validates candidate jobs, enforces resource constraints and manages asynchronous execution while ongoing computations continue (Section C). We evaluate this controller by TM0.6 SU per worker GPU-h under common in silico qualification criteria across seven fixed targets, comparing it with generator-only workflows and implemented no-LLM allocation policies (Sections B and B.5). The related systems discussed above were not rerun here, so this evaluation establishes no performance advantage over them. The comparisons assess the complete controller and do not isolate the contribution of LLM reasoning.

#### B.7. Campaign replication

The campaign was the unit of replication. The primary seven-target benchmark included one campaign per method–target pair. For T-REX, PUCT and *ε*-greedy, two additional independently initialized campaigns on CD45, SC2RBD and CbAgo gave *n* = 3 per controller–target pair, using the same setup and prespecified denominator of 144 H100 worker GPU-h. In Table S9, Run 0, Run 1 and Run 2 correspond to controller seeds 0, 1 and 2, respectively.

### C. Campaign controller implementation

#### C.1. Controller workflow and agent roles

T-REX combines Planner and Supervisor LLMs with deterministic candidate construction, selection and execution. Table S3 distinguishes their inputs and responsibilities; Algorithm S1 specifies the event order. Newly recorded results inform later planning updates (Figure S2b), while qualification and SU credit follow fixed evaluation rules.

**Table S3.**
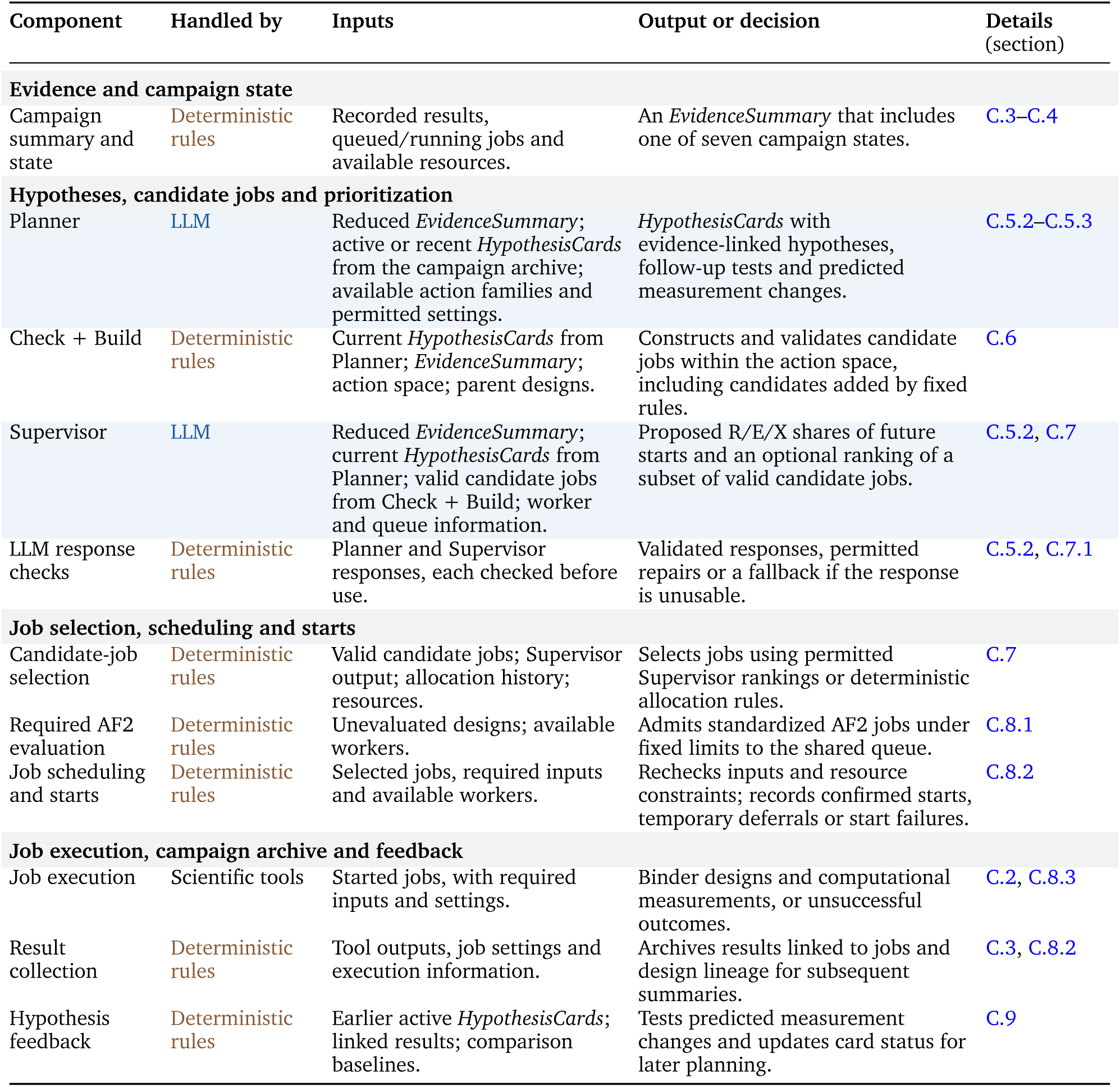
Controller responsibilities, inputs and outputs. Brown and blue distinguish deterministic rules and LLM reasoning (Figure 1d). Rows group responsibilities rather than execution order. Current *HypothesisCards* can include new or retained Planner proposals. Worked examples appear in Sections D.2.1–D.2.2.

**Algorithm notation.** Algorithm S1 uses the campaign archive D (Section C.3), an ordered queue Q of admitted job identifiers awaiting a start, and an ordered pool M of *W* worker slots. Define

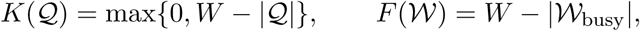

where M_busy_ contains the occupied worker slots. *K* counts positions needed to reach the target queue depth *W* ; *F* counts free workers. AF2 reservations can temporarily exceed the queue target. The campaign wall-time limit *T*_max_ is checked at each loop entry (Section C.8.3). The reporting denominator *H* is defined in Section B.3; the final qualified-cluster count *N*_SU_ follows the clustering rules in Section B.2.

The current EvidenceSummary is *E*, the campaign-state label is *s*, available HypothesisCards are ℋ, valid candidate jobs for Supervisor prioritization are J, and selection decisions are ℒ. J includes generation, redesign and permitted diagnostic AF2 refolds; required standardized AF2 evaluations use the separate reservation procedure (Section C.2.2). The Planner also returns any response warning or fallback reason *f_P_* ; *S* is the Supervisor response. The resolved selection policy P contains permitted candidate rankings, including any usable global order of the subset returned by the current Supervisor response (the *fresh global order*), adjusted **Rescue**–**Explore**–**eXploit** proportions, and each candidate’s priority and resource information. Planner fallback reasons and Supervisor validation or abstention determine which response fields can be used (Section C.7.1). The carried allocation difference **d** tracks allocation targets relative to confirmed starts and is reconstructed from archived decisions before LLM input preparation (Section C.7.3). It is included in the Supervisor’s selection context.

**Procedure conventions.** Procedures update the archive, queue and worker states in place. Planner and Supervisor procedures include input preparation, response validation and call records. Invalid, empty or abstaining Planner output supplies no new cards; retained valid cards can still be used. The Supervisor LLM is called only when cards and valid candidate jobs are available (Section C.5.4).

Before the first dispatch, a state check obtains *s* from recorded results and worker and queue information, or reuses its cached result. It appends no EvidenceSummary and calls neither LLM. DispatchReady rechecks queued jobs and records starts, deferrals or failures, with bounded retries (Section C.8.2). Running jobs continue during synchronous LLM calls; new starts are attempted at the two explicit dispatch points. State-check caching and planning-failure handling are described in Section C.8.3.

In each controller iteration, *b* is the shared AF2 reservation ceiling in jobs, and *u* counts the jobs newly reserved so far in that iteration. ReserveAF2 inserts eligible evaluations at the front of the queue, updates *u* in place and returns newly reserved identifiers; V stores only the second call’s identifiers. Both calls share *b*, with each call bounded by max{0, *b* − *u* − *n*_AF2_(M)}, where *n*_AF2_ counts running standardized AF2 jobs. Reservation limits, exceptions when one queue position is available, deep-stall restrictions and admission of evaluations to otherwise unused queue positions follow Section C.8.1.

**Algorithm S1.**
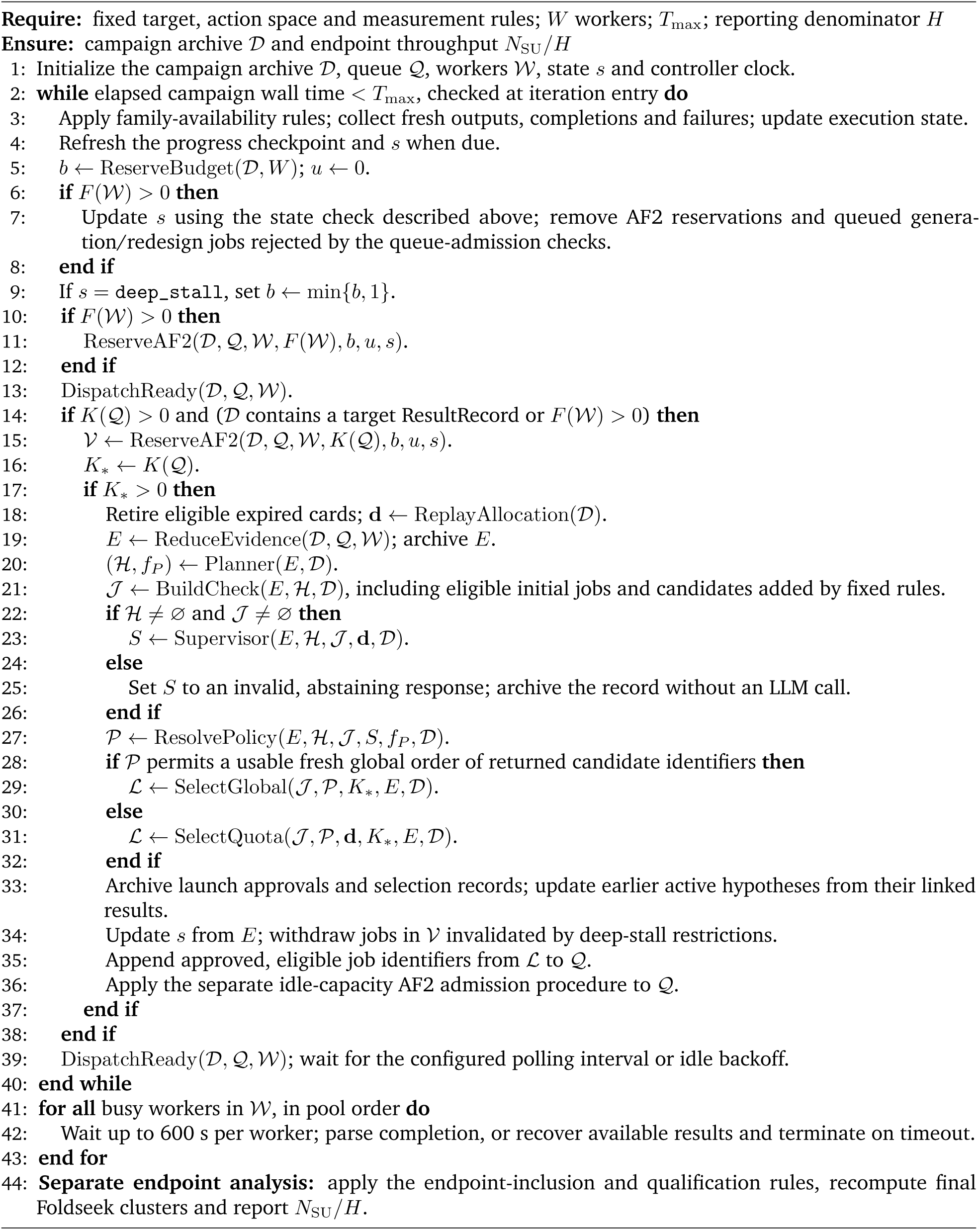
T-REX campaign control and asynchronous execution. Queue admission and worker starts are distinct. Required AF2 evaluation follows a separate admission policy; endpoint analysis is performed separately.

#### C.2. Software and permitted job configurations

##### C.2.1. Software versions

Table S4 lists the software versions, source revisions, model checkpoints and configurations used in the reported campaigns and analyses. Baseline settings and adaptive initialization are described in Section B.4.

**Table S4.**
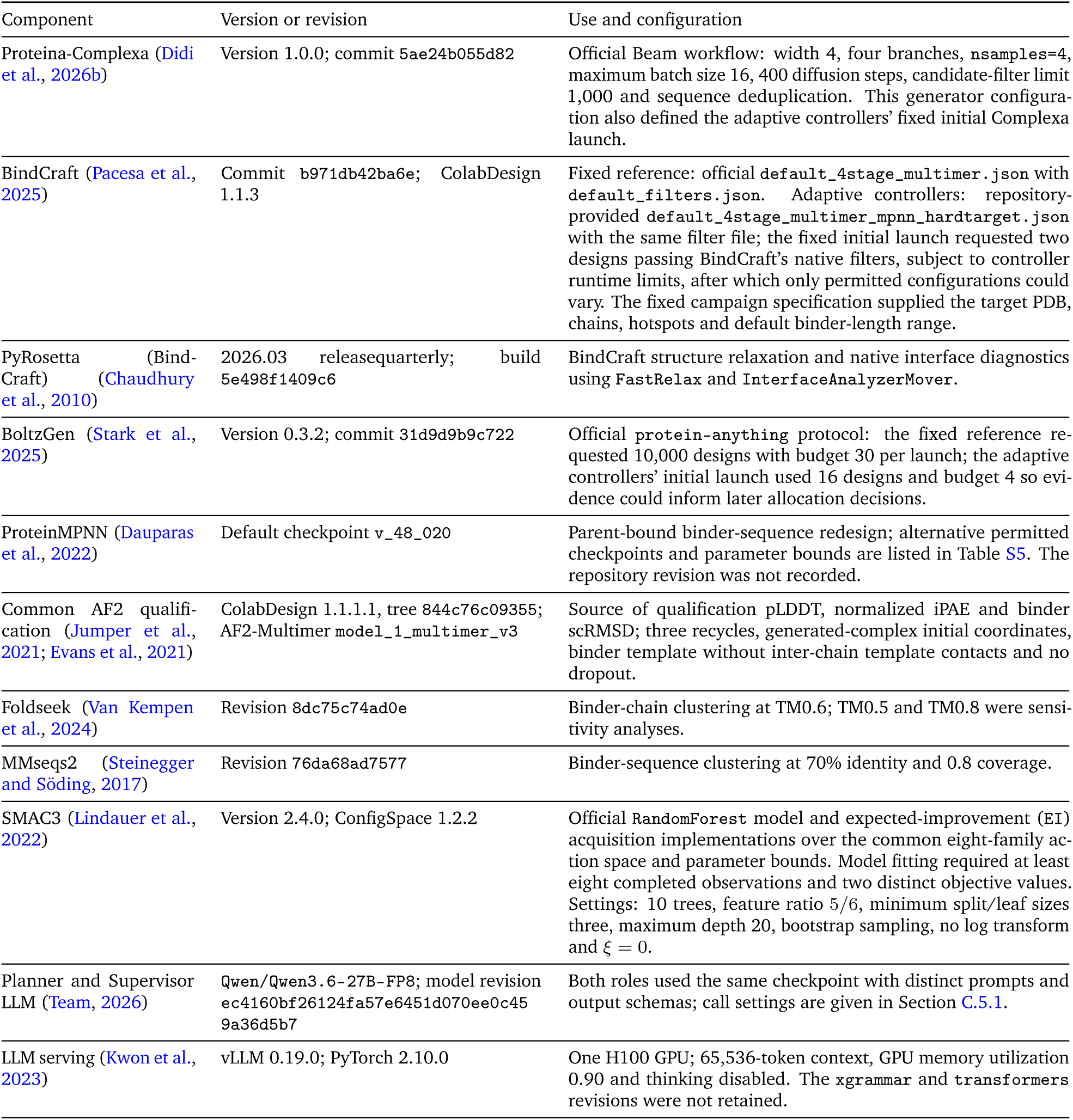
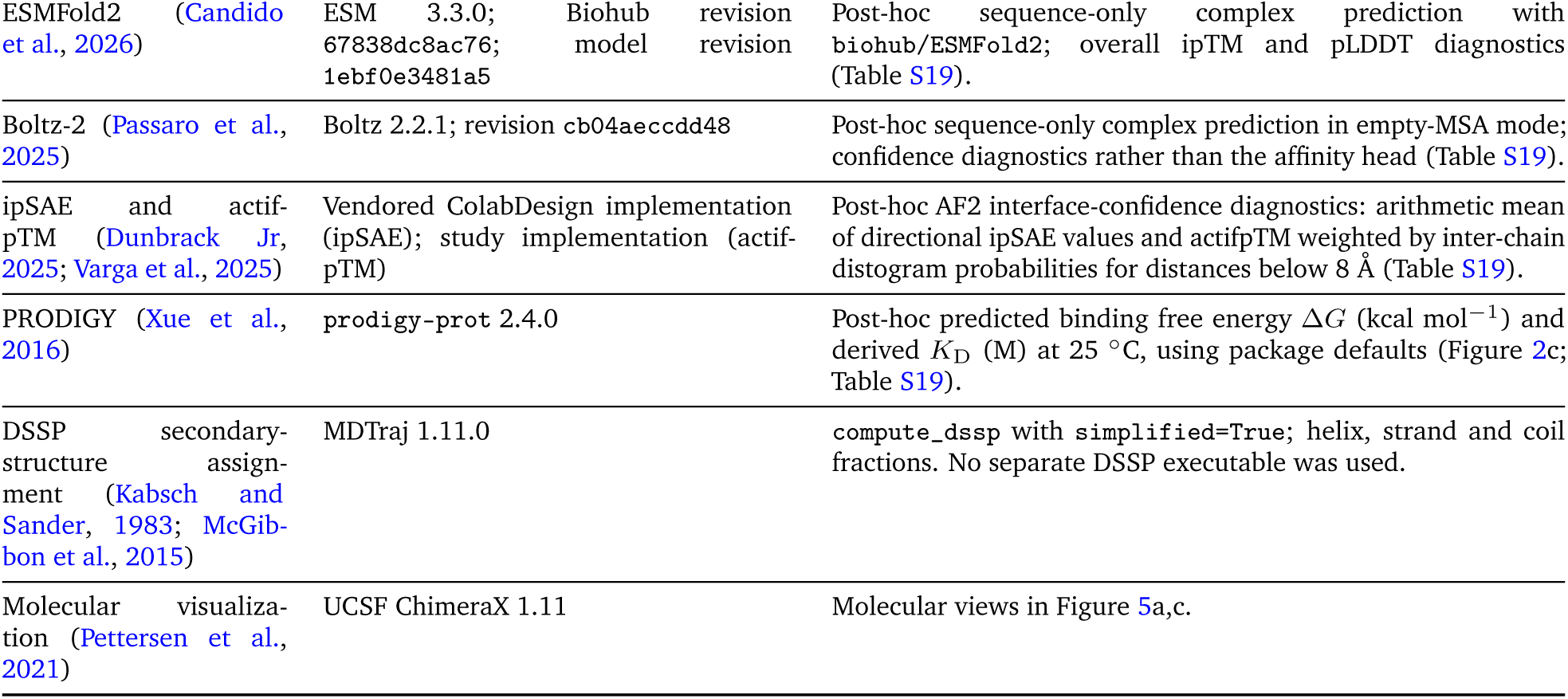
Software, model checkpoints and analysis configurations. Versions and revisions identify the primary generation, evaluation, clustering, LLM-serving and visualization tools. Post-hoc evaluator inputs, settings and score aggregation are detailed in Table S19; LLM serving is detailed in Section C.5.

##### C.2.2. Action space and parameter bounds

All adaptive controllers used the same eight action families: six generation families (Complexa beam search, best-of-*n*, Feynman–Kac steering and MCTS, BindCraft and BoltzGen), ProteinMPNN redesign and AF2 structure evaluation. An action family identifies the tool and, for Complexa, the search variant. A *generator root* groups generation families implemented by the same generator: the four Complexa variants share one root, while BindCraft and BoltzGen each form a separate root.

Each job specifies an action family, a configuration (model and hyperparameter settings) and, for redesign or evaluation, an archived parent design. A *route* groups jobs with the same generation or redesign family and configuration; redesign routes also distinguish the family and configuration that produced the parent design. For example, ProteinMPNN redesign of Complexa and BindCraft outputs forms different routes even when the ProteinMPNN settings match. Standardized AF2 evaluation is attributed to the route that produced its input; route outcomes and costs are defined in Section C.4.2.

The action families and parameter bounds in Table S5 were fixed before each campaign. The Planner and Supervisor proposed and prioritized jobs within this space, while newly available parent designs changed which candidates could be constructed (Section C.6). The space also permits diagnostic AF2 refolding with changed model, recycle or initialization settings. When eligible, these candidates follow the same selection and R/E/X start-accounting rules as generation and redesign jobs, but cannot supply common qualification measurements. Required standardized AF2 evaluation uses fixed settings and separate deterministic admission and ordering (Section C.8.1). The reported allocation analyses concern generation and redesign starts (Section D.3.2).

The per-launch limits in Table S5 constrain validated family-specific workload settings, expressed as counts or products of configuration values. For BindCraft, the controller field max_trajectories is forwarded as number_of_final_designs and specifies the requested number of designs passing BindCraft’s native filters, not a cap on attempted trajectories. Native BindCraft acceptance is distinct from standardized qualification and SU counting. Actual trajectory and output counts must be read from execution records; controller runtime limits can terminate a job before the requested count is reached. These workload settings do not measure GPU hours and are not comparable across families.

**Table S5.**
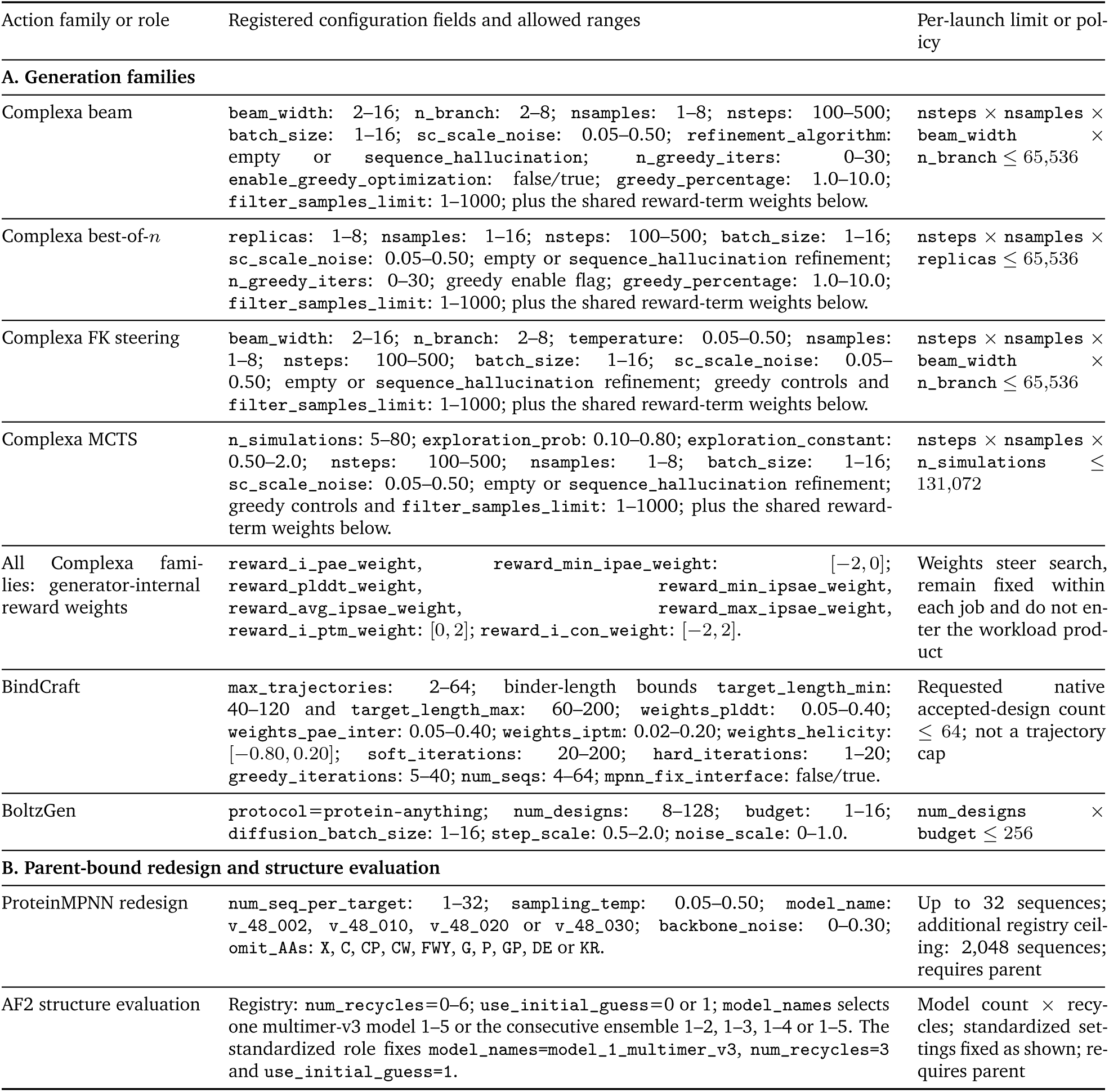
Permitted generation, redesign and evaluation action space. Numeric intervals are validation bounds rather than a discretized grid. The eight families contain 103 configuration fields: 38 integer-valued ranges, 51 continuous ranges and 14 categorical fields, counting shared Complexa controls separately for each family. Parent-bound jobs require an archived structure. The three AF2 fields permit diagnostic refolding and are included in these counts. Standardized qualification uses the fixed settings shown; diagnostic refolds cannot supply common qualification measurements. For BindCraft, the controller field max_trajectories is forwarded as number_of_final_designs: a requested native accepted-design count, not a cap on attempted trajectories.

#### C.3. Campaign archive and runtime state

The campaign archive stores job results, settings, proposals and decisions, with links to structures, score files and logs. Each primary campaign retained nine append-only JSON Lines (JSONL) streams, with one JSON object per line, grouped by role in Figure S1a. Scientific measurements in the *EvidenceSummary* come from recorded job results. The controller combines these records with information on queued and running jobs, designs awaiting AF2 evaluation, worker availability and time remaining for new launches (Section C.8).

**Results and selections.** The *ResultRecords* stream stores generation, redesign and evaluation outputs or failures, with measurements, worker cost, status and parent links. Each record has a unique identifier (result_id); one job can produce several records, including outputs collected before it finishes or a record retaining cost without a usable design (Section C.8.2). The *Panel selections* stream stores nominated design identifiers and their selection checks when emitted.

Within a *ResultRecord*, the parsers store measurements in metrics. The bins field stores annotations such as measurement sources, evaluation roles and chain labels; artifacts stores output paths and chain identifiers. A stored structure may still require standardized AF2 evaluation. Parent and action identifiers link that evaluation to the generation or redesign route that produced its input. The tick_id identifies the planning update associated with the job’s start, so its results can first appear in a later *EvidenceSummary*. Online cluster assignments can change as results accumulate; previously archived bins remain unchanged (Section C.4.2).

**Evidence and decisions.** The *EvidenceSummary* stream stores cumulative qualification and online SU counts, recent results, outcomes and costs by route, clustering status and structural duplication, and recent planning decisions at each planning update (Section C.4; JSON example in Section D.2.1). The *HypothesisCards* stream stores Planner hypotheses, proposed tests and rule-based feedback; updates append records under the same hypothesis identifier (Section C.9). *ActionCandidates* stores constructed and checked candidate jobs. *SupervisorDecisions* stores allocation proportions, an optional ranking of a candidate subset, the capacity information supplied to the Supervisor, rationale and selection checks. *LLM-call records* retain model identifiers, prompt hashes, response validation status, latency, token use and failure or fallback information, together with selected prompt fields; full prompt text is not retained.

**Launch and execution.** The *LaunchDecisions* stream records deterministic approval or rejection after selection checks. Approval permits queue admission but does not establish that a job started. The *DispatchRecords* stream separately records confirmed starts, failed start attempts, temporary deferrals and failures to collect job outputs, with worker identifiers, attempt numbers and reasons (Section C.8.2). Post-start outputs, measurements and costs are stored as *ResultRecords*. Current queue membership and running jobs are tracked separately from these event records.

**Figure S1.**
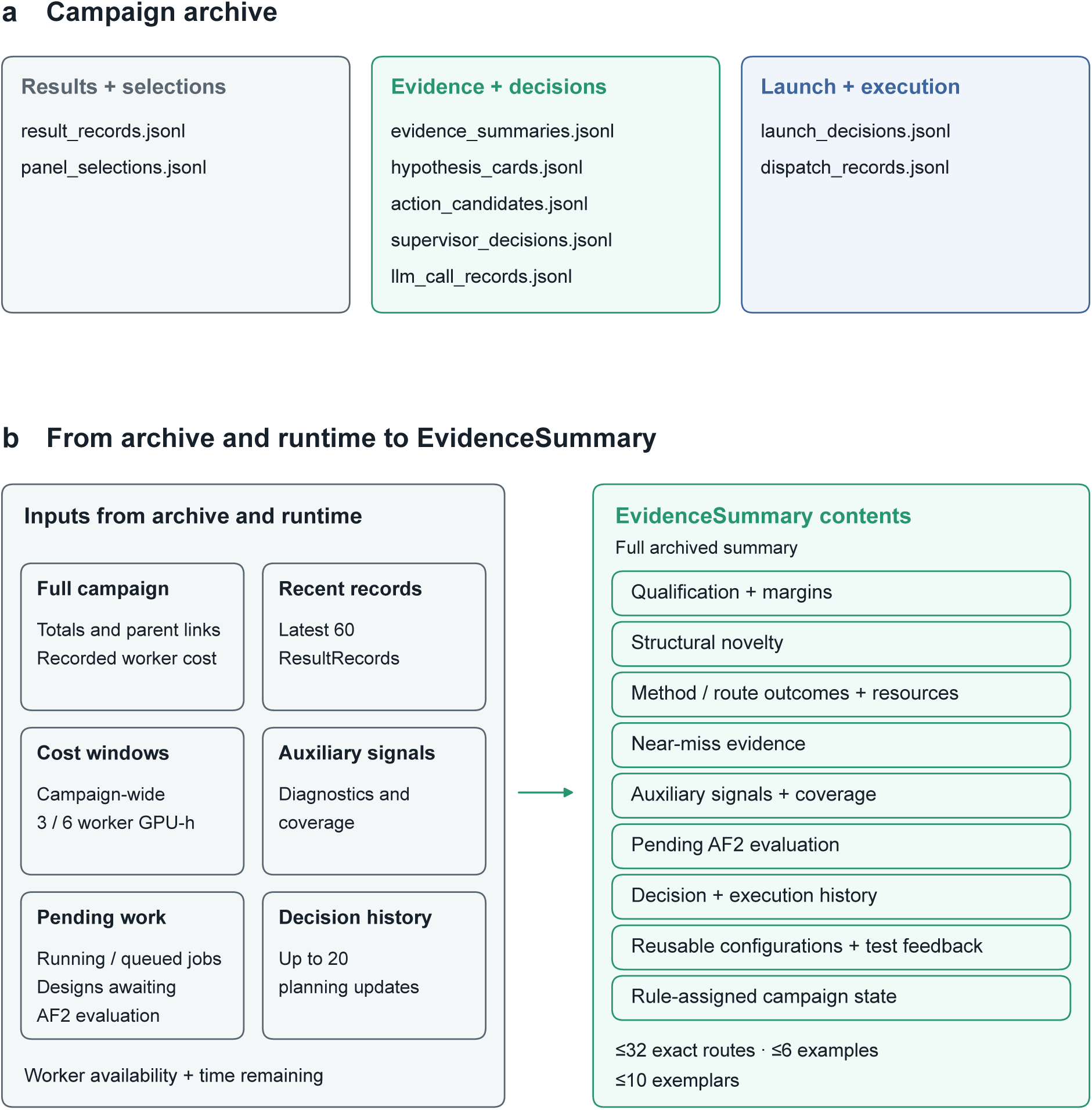
Campaign archive and *EvidenceSummary* construction. **a**, The nine append-only record streams retained in the primary campaigns are grouped by role (Section C.3). ResultRecord stores measurements, parent links, worker cost, status and output paths. Raw logs and files remain in the run directory; LLMCallRecord retains a prompt hash and compact audit rather than full prompt text. **b**, Deterministic rules combine archived records with information on current jobs and resources to construct the full archived *EvidenceSummary* (arrow). The 3- and 6-worker-GPU-h windows use campaign-wide recorded worker cost. Figure S2 shows the smaller input supplied to the LLMs. Final qualification and SU counts are recomputed separately after applying endpoint-inclusion rules (Section B.3). Gray indicates results, selections and reduction inputs; green, evidence/decision records and summary contents; and blue, launch/execution records.

#### C.4. EvidenceSummary construction and derived measures

At each planning update, the controller combines archived results with queued and running jobs, designs awaiting AF2 evaluation and available worker capacity to construct an *EvidenceSummary*. It reports cumulative qualification and online SU counts, recent results, and route outcomes and costs (Figure S1b). Section C.5.2 describes the smaller input supplied to the LLMs; Section D.2.1 provides a JSON example.

##### C.4.1. Evidence contents and record windows

**Measurements and missing values.** Qualification summaries report median measurements and median distances from the qualification cutoffs, counts of pass, near-pass and failed measurements, and paired-criterion outcomes (Section B.1). In archive field names, strict denotes computational qualification. Online Foldseek TM0.6 assignments and duplication measures include clustering status and coverage. Auxiliary confidence, interface and geometry measurements retain their source and availability. Missing measurements are not imputed or treated as failures; metric_availability identifies which measurements are present.

**Record windows and minimum observation counts.** Cumulative qualification, SU and parent-history fields use all observed results. Recent qualification summaries and combinations of criterion outcomes use the latest 60 ResultRecords (Section C.3). General auxiliary summaries combine the latest 40 records per family with the latest 60 overall, including each record only once. They require at least three finite observations per measurement; the route-level auxiliary score has a separate requirement (Section C.4.3). The 3- and 6-worker-GPU-h windows select records backward from the newest result until their summed recorded cost reaches the specified amount, including the record that reaches or crosses it. If less cost has accumulated, all records are used. Each campaign-wide window is shared across routes.

The recent duplicate fraction is

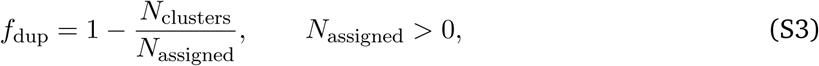

where *N*_assigned_ counts records with an online Foldseek TM0.6 cluster assignment among the latest 60 ResultRecords, and *N*_clusters_ counts their distinct clusters. The fraction is unavailable when *N*_assigned_ = 0. This recent duplication measure is separate from final qualified-binder clustering.

**Near misses and counting units.** A *deficit* is the nonnegative distance by which a measurement misses its qualification cutoff. The *near-pass tolerances* are 5 pLDDT points, 0.05 normalized iPAE and 0.3 Å scRMSD. A near miss has finite values for all three measurements and fails qualification, with at most one criterion outside its tolerance and no failed criterion more than ten corresponding tolerances from its cutoff. Near misses are clustered separately from qualified designs and receive no SU credit.

The campaign-state input near_miss_count counts distinct near-miss Foldseek clusters in the latest 60 ResultRecords, subject to the reliability checks in Section C.4.2. Method summaries count cumulative near-miss clusters; their recent near-miss yield counts clusters first observed in the latest 60 ResultRecords and attributed to each family. In contrast, near_miss_count and near_miss_recent in exact-route rows count individual near-miss ResultRecords; family rows in route_values sum these route counts. The same field name therefore has different counting units in campaign and route summaries.

**Work awaiting completion and worker availability.** The summary includes pending AF2 evaluations (diagnostic_chain_backlog), queued and running jobs (pending_family_load), failures, timeouts, evaluation-role outcomes and available workers. It also identifies usable redesign parents and parent–child lines flagged for non-improvement. The dispatch_realization and execution_realization fields distinguish proposals, selections and starts (Section C.3). Section D.2.1 illustrates how measurements, missing values and pending work appear in the summary and the LLM input.

**Method summaries and reusable evidence.** The method_health summary combines family-level attempts, completions, failures, costs and qualification/novelty outcomes to distinguish untried, delayed and unproductive methods. Its recent rates use new SU gains in the latest 60 ResultRecords. Direct rates divide those gains by the family’s recorded cost in that window. Rates attributed through downstream evaluation also include its recorded cost and required ancestor costs outside the window, counting each ancestor once within the family. These method summaries are distinct from the family aggregates in route_values; they supply the recent family rates used in selection and concurrency checks (Sections C.7.2 and C.8.2).

A previously tested configuration summary (recipes) groups results by action family, configuration and outcome category, retaining counts, median measurements and representative result identifiers. Strategy-outcome summaries (strategy_feedback) retain outcomes, costs, qualification failures, near misses and parent–child history for each strategy, including negative and partial results. These are observations for reuse, not guarantees of future success; Section D.2.3 gives worked examples.

**Routes, examples and planning history.** The route_values summary records outcomes, costs, pending evaluation, quality scores and recent or cumulative SU/GPU-h for each route, with separately labeled family aggregates. Section C.4.2 defines route cost and SU attribution. Illustrative result records (examples) link individual outcomes to qualification measurements, deficits and parents where available. Configuration-linked designs (exemplars) are selected qualified or near-miss binders with their producing configurations and parent links, providing concrete evidence for follow-up tests. Each summary retains up to twenty planning-update summaries in recent_ticks_history; the LLM input retains ten (Section C.5.2). HypothesisCards, archived separately, describe proposed tests (Sections C.3 and C.9).

##### C.4.2. Online SU attribution and compute accounting

**Route attribution and cost.** Routes group jobs as defined in Section C.2.2. Standardized AF2 evaluation is attributed to the generation or redesign route that produced its input. Each online qualified cluster adds one SU to the route that produced its first qualified member in archive order. Other routes can contribute qualified members without another SU credit. Final attribution instead follows the job that produced the fixed representative (Section B.2).

Route cost includes generation or redesign, required evaluation and the recorded costs of ancestor designs. Each ancestor ResultRecord is charged once within a route, but can be charged to several routes; summing route costs therefore does not give campaign cost. A multi-output job’s cost is divided among its ResultRecords. Recent route costs use the campaign-wide windows in Section C.4.1, including required ancestor costs even when the ancestor’s record precedes the window. Table S6 distinguishes these costs from the reporting budget and other online time measures.

**Table S6.**
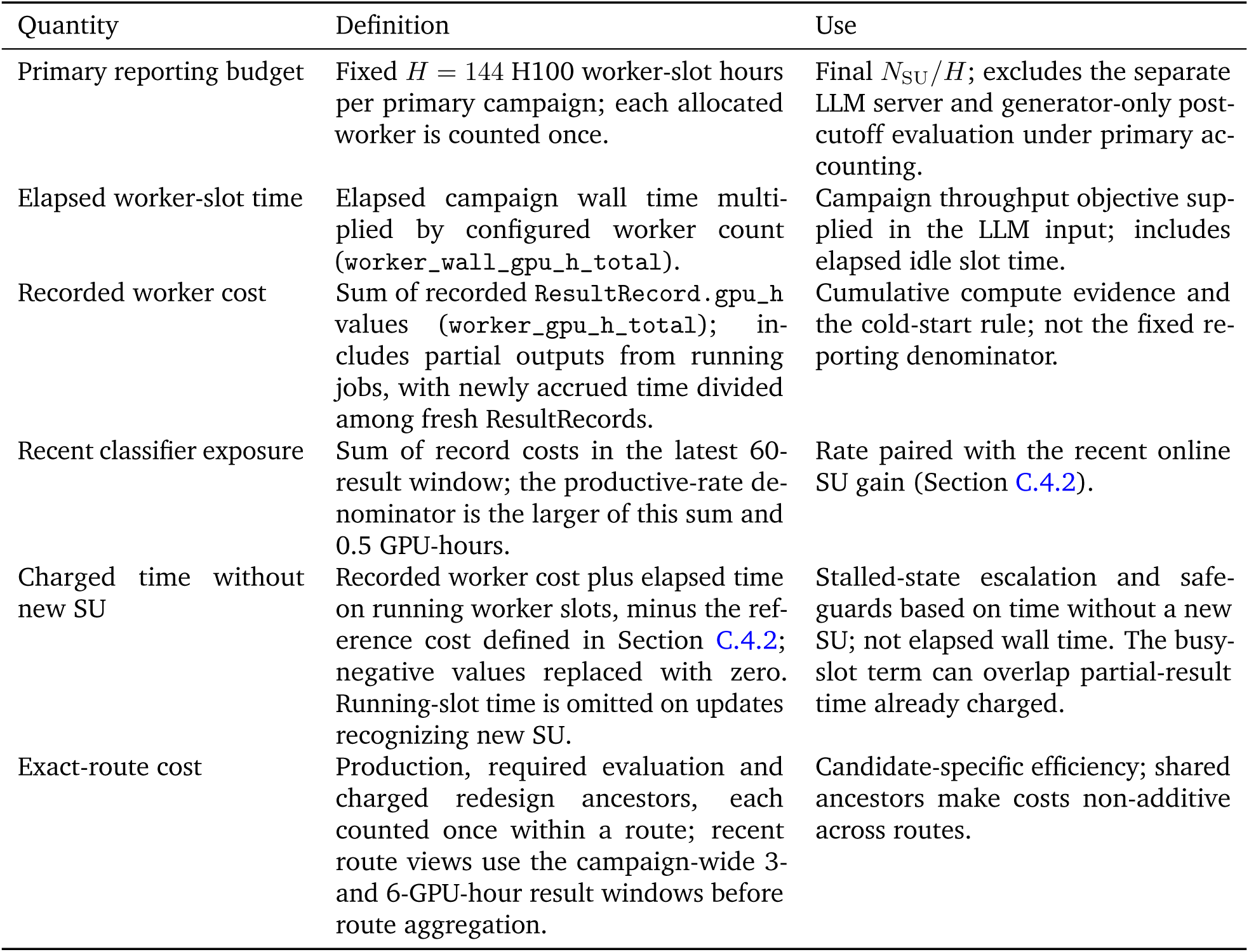
Time and cost quantities used for reporting and control. GPU-hours are time-accounting units, not measurements of GPU utilization or floating-point operations. These quantities have different definitions and must not be substituted for one another.

**New online SU and clustering reliability.** Recent gain counts online SU first represented in the latest 60 ResultRecords. At each update, the controller counts qualified clusters in all records and in records preceding that window. It subtracts the earlier-record count from the full count, replacing negative values with zero. Both counts use current cluster assignments, so the calculation does not compare summaries from different updates.

Productive-state classification requires Foldseek clustering status ok, coverage ≥ 0.999, an available duplicate fraction and no reported fallback from binder-chain to whole-complex clustering. Here, coverage is the fraction of qualified ResultRecords assigned to qualified structural clusters, not the number of clusters. Section C.4.5 specifies the additional progress and rate conditions. A positive near-miss count requires separate clustering with status ok or cached_ok and coverage ≥ 0.999 among near-miss records. Otherwise, the classifier receives zero while the original count, examples and clustering status remain available. This substitution disables the near-miss decision input; it does not establish that no near misses occurred.

**Time without new online SU.** At each update, the controller reconstructs a reference cost by scanning results in archive order. Whenever a qualified design is the first qualified member of its cluster encountered so far, the reference becomes the cumulative recorded worker cost through that result; it is zero before the first SU. If an update reports more online SU than any previous update, the reference is instead the current recorded worker cost.

The stall-time signal is current recorded worker cost plus elapsed time on running worker slots, minus that reference, with negative values replaced by zero. Running time is summed from each running job’s start and omitted on updates recognizing new SU. Partial-output costs already recorded are not deducted from this term, so time can be counted twice (Section C.8.2). This signal detects stalls using different accounting from the reporting budget (Table S6).

##### C.4.3. Online quality and auxiliary scores

**Online quality score and route summaries.** For a qualified design, the margins below measure how much better its measurements are than the qualification cutoffs. Each margin is divided by the corresponding near-pass tolerance:

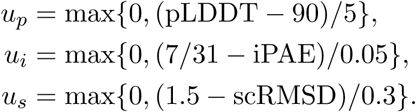

Each value contributes at most 3. Their sum is divided by 9 to give the design score *q*, between 0 and 1:

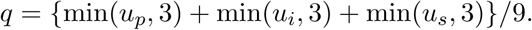

Within each route and online Foldseek cluster, the qualified design with all three measurements and the highest *q* is selected. Each cluster contributes one quality observation, even if another route received its SU credit. A later, higher-scoring design in the same cluster replaces that observation rather than adding one.

The route’s qualified-cluster count is the number of selected designs. The median and 25th percentile of their *q* values are stored as strict_quality_median and strict_quality_p25. The percentile uses inclusive linear interpolation. With one selected design, both equal its score; with none, both are missing. Criterion-specific margin summaries use these same designs. A family summary copies the quality statistics of its route with the highest 25th percentile, breaking ties by the median and then the qualified-cluster count.

These summaries include all currently observed qualified clusters in a route, not only those in a recent window. The Planner and Supervisor receive them to compare routes with similar recent SU throughput (Section C.7). An increase in the median or 25th percentile can also permit additional standardized AF2 evaluations (Section C.8.1). Final attribution, post-hoc analyses and qualification-margin sensitivity instead use *m*_min_ (Sections B.2 and D.1.4). Neither score changes qualification or gives an unqualified design SU credit.

**Auxiliary diagnostic-improvement score.** This score summarizes whether auxiliary measurements meet or approach their quality thresholds, or have improved relative to earlier results. Table S7 gives the source-specific measurements, thresholds, near-pass tolerances, weights and acceptance-reference limits. These references do not add qualification criteria.

For each route and measurement, recent observations are its finite values in the campaign-wide 6-GPU-hour result window (Section C.4.1); prior observations are its finite values in earlier records. A measurement is used in this score with at least two recent observations, whether or not they meet the quality threshold. A single recent observation is used only if it meets that threshold.

The deficit is how far a measurement misses its quality threshold, or zero if the threshold is met. Dividing it by the corresponding near-pass tolerance gives the normalized deficit. Two values are then calculated for each measurement:

*Current quality.* Among the recent observations, those meeting the quality threshold contribute one, those missing it by no more than the tolerance contribute one-half, and the rest contribute zero. The sum of these contributions is divided by the number of recent observations.

*Improvement.* The controller compares the median normalized deficit in prior and recent observations. Only a decrease contributes: the decrease is divided by the larger of one and the prior median. Without prior observations, this term is zero.

For each measurement, the larger of these two values is multiplied by its Table S7 weight: 1.0 if a registered parameter directly addresses that measurement, or 0.6 otherwise. The route score is the largest weighted value across its measurements. It is zero if observations are insufficient or if no measurement yields a positive value. Zero therefore cannot distinguish missing evidence from observed values that yield no positive score, and does not by itself establish deterioration.

Scores lie between zero and one and are stored rounded to three decimal places. An action-family summary uses the largest score among its exact routes. The score can support further evaluation (Section C.8.1) or prevent premature route deferral (Section C.7), but cannot qualify a design or replace new-SU throughput as the ranking objective.

The general auxiliary summaries are separate from this route score. They require at least three finite observations and also check the acceptance-reference limits in Table S7. Values failing a limit are counted as failures even if they fall within the near-pass tolerance of the quality threshold. A dash means that no acceptance-reference limit applies. This additional check is not used in the route score.

**Table S7.**
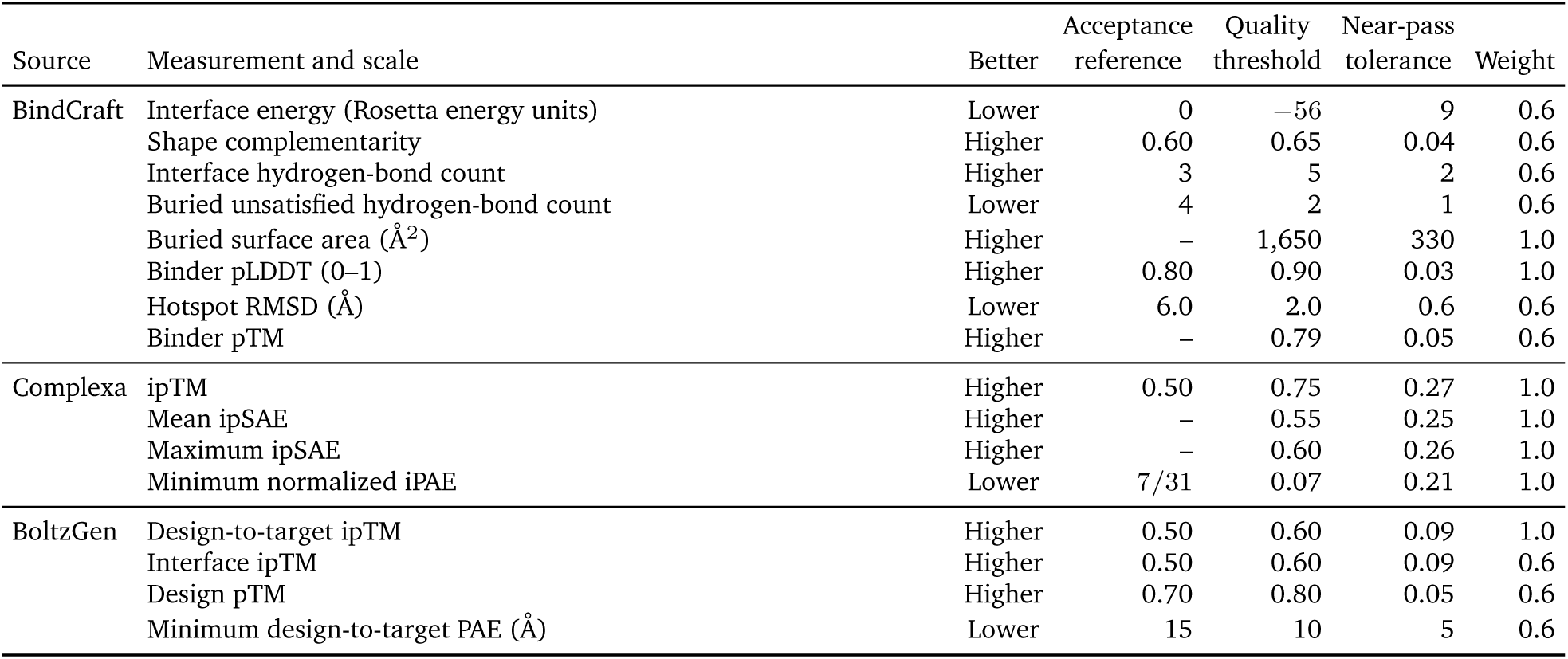
Reference values and weights for auxiliary measurements. Quality thresholds and near-pass tolerances define the auxiliary diagnostic-improvement score; a near pass misses the quality threshold by no more than the tolerance. Acceptance-reference limits additionally constrain the general auxiliary summaries, not the route score. Limits use the direction indicated in the *Better* column; a dash means no additional limit. A weight of 1.0 identifies a measurement with a registered directly relevant parameter; 0.6 denotes corroborating evidence. These source-specific references cannot establish common qualification or SU credit.

##### C.4.4. Route status rules

Each route has a general status (status) used for evaluation admission and route deferral, and a recent-productivity status (marginal_status) distinguishing continuing, delayed and declining yield. Both return the first matching rule below and identify pending evaluation separately. Family aggregates use the same rules, except that the 32-completion test is disabled and no qualified-cluster duplicate fraction is supplied. These statuses describe routes or families; Section C.4.5 defines whole-campaign states.

**Window definitions.** The *60-record window* contains the latest 60 campaign ResultRecords; the *3- and 6-GPU-hour windows* use campaign-wide recorded cost (Section C.4.1). A gain is new online SU credited to a route within the stated window. Recent near misses are individual near-miss records in the 60-record window, subject to the clustering checks in Section C.4.2. Route cost includes the applicable ancestor costs; lifetime rates use the full observed history.

**Duplicate measures.** Qualified/SU is the route’s qualified-record count divided by its credited SU count; the ratio is unavailable when the SU count is zero. The route duplicate fraction uses Eq. (S3), with *N*_assigned_ and *N*_clusters_ counted over the route’s cumulative qualified cluster-assigned records. A duplicate warning means qualified/SU ≥ 8 or duplicate fraction ≥ 0.80. Missing duplicate measures cannot satisfy their threshold tests.

**Comparison rate.** The best mature rate is the largest lifetime SU/GPU-h rate among exact routes with at least two credited SU and two route GPU-hours; zero if none qualifies. Missing rates are treated as zero only for the rule comparisons below.

**AF2 structure-evaluation rows.** AF2 evaluation rows bypass the generation/redesign classifications below. Standardized evaluation carries the archive label plumbing; the code reserves advisory for other evaluation roles, including an intentional refold of a parent model.

**General route status (**status).

1. untried: No positive accumulated route cost.
2. awaiting_score_conversion: No credited SU and at least one promising valid output awaiting standardized AF2 evaluation. Admission signals are defined in Section C.8.1.
3. collapse_risk: Either: a duplicate warning, at least one credited SU, at least eight qualified records and no 60-record or 6-GPU-hour SU gain; or qualified/SU ≥ 8, lifetime rate ≤ 0.30, and either no 6-GPU-hour gain or a positive 60-record gain.
4. defer: Either: neither a lifetime SU nor a recent near miss, with at least 3 route GPU-hours or 32 completed ResultRecords; or a best mature rate ≥ 0.50, at least 3 route GPU-hours, a lifetime rate below 35% of that best rate, and no 60-record gain, 6-GPU-hour gain or recent near miss. Completed ResultRecords here have ok or no_artifacts status; one job can produce several records.
5. observed (stale): At least one lifetime SU, but no 60-record gain, 6-GPU-hour gain or recent near miss, with at least 1 route GPU-hour in the 60-record window.
6. diversify: At least one lifetime SU and either qualified/SU ≥ 4 or duplicate fraction ≥ 0.55.
7. promote: At least one lifetime SU, a positive best mature rate and a lifetime rate at least 80% of that comparator. With no 60-record SU gain, qualified/SU ≥ 8 or duplicate fraction ≥ 0.70 prevents promotion.
8. healthy; otherwise observed: After the preceding rules, at least one lifetime SU or a recent near miss gives healthy; otherwise observed.

**Recent-productivity status (**marginal_status).

1. untried; awaiting_score_conversion: Apply the same first two tests as the general-status classification.
2. under_tested; otherwise dry_low_quality: For a route with zero lifetime SU, under_tested applies while accumulated route cost is below 3 GPU-hours and completions are below 32 ResultRecords, or while a recent near miss exists. Otherwise use dry_low_quality.
3. productive_but_duplicate: A duplicate warning and an SU gain in either the 60-record or 3-GPU-hour window.
4. delayed_productive; duplicate variant: An SU gain in the 6-GPU-hour window but none in either the 60-record or 3-GPU-hour window. A duplicate warning changes the label to delayed_productive_duplicate.
5. dry_duplicate: A duplicate warning, no 3-GPU-hour or 6-GPU-hour SU gain and recent rate decay: at least 1 route GPU-hour within the 3-GPU-hour window, with the larger SU/GPU-h rate from the 60-record and 3-GPU-hour windows below 35% of the positive lifetime rate.
6. productive: An SU gain in either the 60-record or 3-GPU-hour window.
7. dry; otherwise observed: The same recent-rate-decay test and no 6-GPU-hour SU gain give dry; all remaining routes are observed.

##### C.4.5. Campaign-state rules

The controller deterministically assigns one of seven campaign states from worker cost, recent results and new SU, duplication, near misses, criterion-specific deficits and time without new SU. State labels guide priorities and fallback allocation (Section C.7.1). low_evidence favors gathering more evidence; productive favors continuing productive routes; productive_duplicate adds diversification; and strict_duplicate_collapse favors diversification or alternative generator configurations. rescue_rich directs **Rescue** toward the limiting criterion, while stalled favors **Rescue** or **Explore** over exploitation and deep_stall strongly deprioritizes unproductive routes. Labels neither change qualification nor prescribe a particular job.

Criterion concentration is computed separately from the near-miss count. For criterion *a*, let *d_a_* be its median nonnegative qualification deficit among available measurements in the latest 60 ResultRecords (axis_stats), and *b_a_* its near-pass tolerance (Section C.4.1). The classifier uses

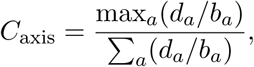

with value zero when no positive median deficit is available. Missing medians do not contribute. Thus the 0.50 cutoff below describes the concentration of scaled median deficits, not the fraction of near-miss designs failing one criterion. Scaling makes deficits in pLDDT, normalized iPAE and scRMSD comparable for this diagnostic.

**Decision order and cold start.** Cumulative recorded worker cost below 1.0 worker GPU-h or fewer than three recent ResultRecords returns low_evidence. Otherwise, the first matching rule applies in this order: structural duplicate collapse; productive; productive with duplication; near-miss enrichment, with possible deep-stall escalation; stalled or deep stall; and default low evidence. This classification rule is separate from the initial-launch policy (Section B.4).

**Structural duplicate collapse.** This state requires reliable clustering as defined in Section C.4.2, at least 40 qualified designs, at least two SU clusters, at least eight qualified designs per SU on average, cumulative SU/GPU-h no greater than 0.30 and a recent SU gain no greater than one. When available, the ratio of Foldseek TM-score 0.8 to TM-score 0.6 cluster counts within the same recent qualified-design window must also be no greater than 1.30.

**Productive and near-miss states.** Productivity requires reliable clustering, an available duplicate fraction, a positive recent online SU gain and at least 0.04 new SU per worker GPU-h. This rate divides the 60-record gain by recorded worker cost in that window, with a minimum denominator of 0.5 GPU-h (Table S6). A duplicate fraction below 0.60 gives productive; otherwise it gives productive_duplicate. Near-miss enrichment requires at least three near-miss clusters in the 60-record window and *C*_axis_ ≥ 0.50. If reached, this branch returns deep_stall after at least 12.0 worker GPU-h without new SU, and rescue_rich otherwise.

**Stalled states.** If no earlier rule applies, the stalled branch is entered when recent online SU gain is zero, clustering is unreliable, or the largest cluster contains at least 75% of assigned structures in the 60-record window without a productive rate. It returns deep_stall after at least 12.0 worker GPU-h without new SU and stalled otherwise. Missing stall-time information cannot trigger escalation; Section C.4.2 defines this signal. The 12.0-worker-GPU-h test does not override the earlier cold-start, collapse or productive rules. These fixed rules were calibrated as described in Section B.4; archived state labels describe their outcomes in the reported campaigns.

#### C.5. Language-model inputs, outputs and call handling

##### C.5.1. Model and call settings

Both roles used a local Qwen3.6-27B-FP8 checkpoint served by vLLM on one H100 GPU; Table S4 specifies the model revision and software versions. The context limit was 65,536 tokens and GPU memory utilization was 0.90, with enable_thinking=false and --trust-remote-code. Calls used max_tokens=3072, timeout_s=90 and OpenAI-client max_retries=1. Planner temperature was 0.2 with prompt_variant=default; Supervisor temperature was 0.0. Zero temperature did not ensure identical responses; Table S15, part B, reports controlled reruns.

Confidence below 0.55 was recorded without rejecting an otherwise valid response. Supervisor confidence below 0.65 could activate allocation bounds (Section C.7.1); this was a separate use of confidence. Each call recorded the model identifier, a 16-character SHA-256 prompt-hash prefix, output-format version, parse status, failure reason when present, and input/output token counts. These records distinguish LLM calls from deterministic advisory checks (Section C.5.4); Section C.3 describes their storage.

**Figure S2.**
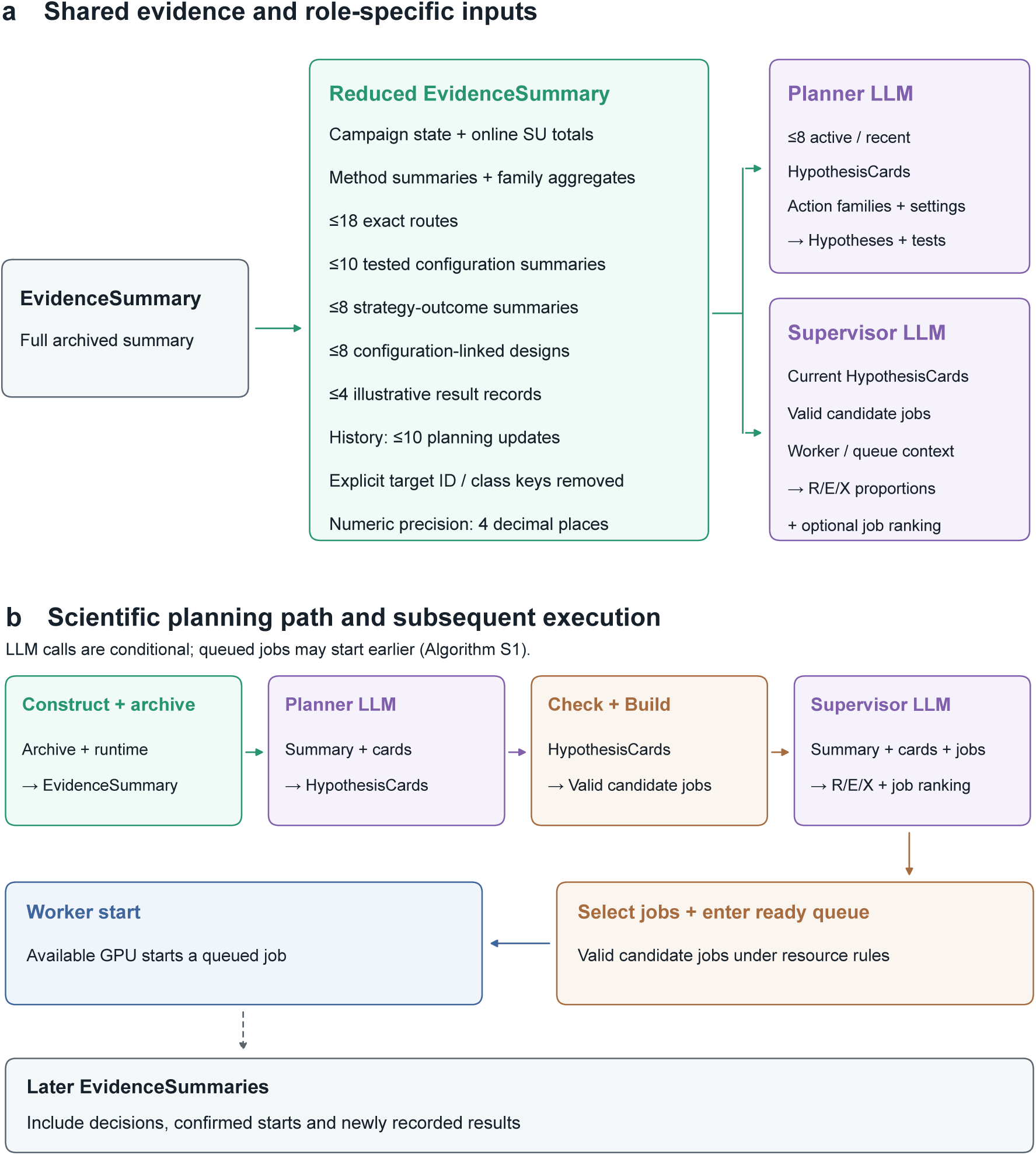
LLM inputs and the order of planning and execution. **a**, The reduced *EvidenceSummary*, including campaign state, is shared by both LLMs (limits: Section C.5.2; route and collection definitions: Sections C.2.2 and C.4.1). The Planner additionally receives up to eight active or recent HypothesisCards, available action families, permitted settings and workload limits; the Supervisor receives current HypothesisCards, valid candidate jobs and selection context. Explicit target-identifier and target-class keys are removed from the evidence view. **b**, Purple denotes LLM proposals and prioritization, brown deterministic candidate construction and selection, and blue worker execution. Each EvidenceSummary is archived before conditional LLM calls (Algorithm S1). Queued jobs may start earlier; queue entry does not guarantee an immediate start. Later summaries include decisions, confirmed starts and newly recorded results. Read-only evidence refreshes before a job starts append no EvidenceSummary and call neither LLM.

##### C.5.2. Input contents and requested outputs

**LLM input contents and size limits.** An archived *EvidenceSummary* retained up to 32 exact-route rows and 20 planning-update summaries. A second reduction supplied both LLMs with cumulative online TM0.6 SU totals, all method summaries and family-level route rows, and at most 18 prioritized exact routes, ten tested configuration summaries (recipes), eight strategy-outcome summaries (strategy_feedback), eight configuration-linked designs (exemplars), four illustrative result records (examples) and ten update-history entries (Figure S2a; definitions in Section C.4.1). The Planner also received up to eight active or recent HypothesisCards, available action families, permitted settings and per-job workload limits. The Supervisor received current cards, valid candidate jobs and selection context, including available worker slots and recent starts.

Exact-route rows preceded family summaries. The reduction replaced the full parent-identifier list with a count and availability flag; concrete identifiers remained available in selected examples and for candidate construction. It removed explicit target-identifier and target-class keys recursively, omitted top-level missing scalars and empty strings, and retained empty evidence collections such as recipes and joint_patterns and method summaries with zero attempts. These omissions did not impute measurements.

Both dynamic prompts began with a short evidence digest followed by role-specific JSON. Section D.2.1 shows an archived EvidenceSummary excerpt and a reconstructed reduced-input excerpt, rather than a saved complete prompt. Neither role read arbitrary archive files or wrote measurements through this interface. Figure S2b places input preparation, calls and subsequent records within the planning update.

**Requested outputs.** Both roles requested one JSON object without server-enforced JSON formatting. Planner fields were abstain, confidence, rationale and cards, with optional fail_reason. A non-abstaining Planner response was requested to contain one to four cards, each citing evidence and specifying a claim, R/E/X affinities (relative weights used as priority hints), predicted changes in one or more qualification measurements, preservation constraints, proposed families and optional settings. Predicted changes shared the same ordered baseline references; Section C.9 defines their later evaluation. Section D.2.2 provides an archived card and its constructed candidate job.

The Supervisor returned a mode_mixture and a ranked candidate_decisions list, with abstention, confidence and rationale fields. Each returned candidate belonged to one priority, with unique global ranks from 1 to the number returned and unique positive ranks within each priority. Allocation values were finite, nonnegative and summed to one. The JSON below illustrates the requested fields for one candidate; bracketed strings are placeholders and the numerical values are illustrative. Section D.2.2 gives an actual prioritization example.

**Illustrative Supervisor response (JSON).**

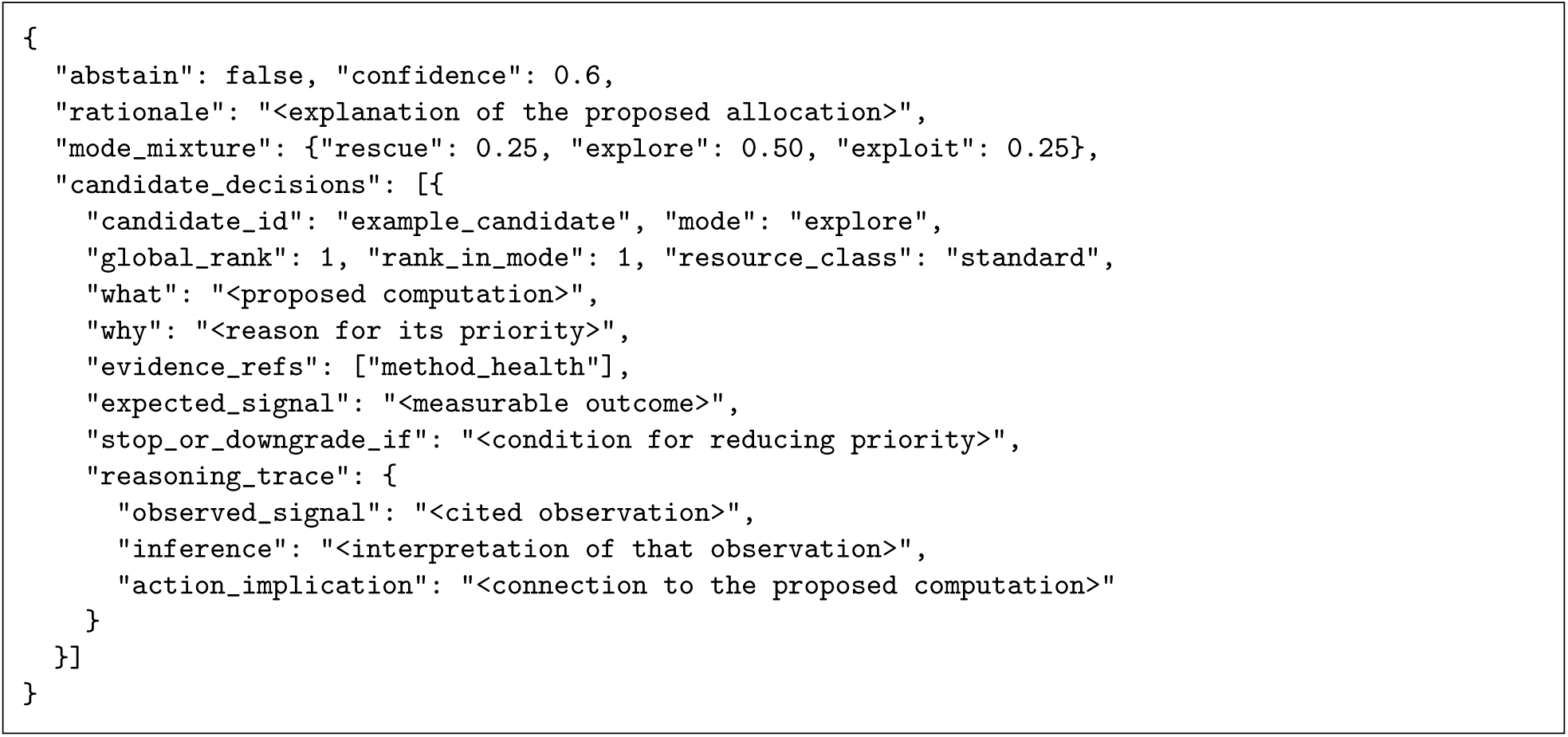

Each card and decision requested three short explanation fields in reasoning_trace: observed_ signal, inference and action_implication. These were reported explanations, not records of private model reasoning. A Supervisor response could rank only a subset of candidates, including none. Its resource classes supported auditing and tie-breaking; allocation bounds and execution remained deterministic (Sections C.7.1 and C.8.2).

**Parsing and validation.** Parsing extracted the first JSON object and normalized recognized field variants. Planner validation checked field types, evidence references, measurement directions and priority affinities; unavailable families were removed and candidate construction enforced setting bounds. Supervisor validation checked candidate/evidence references, allocation values, priority assignments, ranks and resource classes. Missing Supervisor explanation fields could be filled from returned references and explanation text. Planner repairs were deterministic. Only Supervisor errors involving duplicated candidates across priorities, invalid global ranks or unknown candidate identifiers permitted one corrective LLM call. Limited deterministic repair could then remove unknown candidate rows. Repaired outputs were revalidated; unusable responses followed Section C.7.1.

##### C.5.3. Prompt instructions

Each role receives a system prompt defining its task and a dynamic user prompt containing the current evidence and role-specific inputs (Figure S2a). The boxes below summarize the retained campaign implementation: system instructions are condensed, input bodies are schematic, and the corrective message is a template excerpt. They are not complete historical prompts, which were not archived. In the input schematics, bracketed strings stand for structured records or text supplied at runtime. Section C.5.2 specifies the requested output fields; Section D.2.2 provides archived responses and linked candidate jobs.

###### Shared scientific instructions (condensed)

Base decisions on supplied outcomes and constraints, without assumptions about target identity, target class or which method should perform best. Maximize new TM0.6 SU per elapsed worker-slot hour. for method and configuration comparisons, use recent exact-route outcomes and complete route costs before family summaries; use longer windows to account for delayed feedback.

Qualified duplicates add no new SU. Pending evaluation, unstarted jobs and missing measurements are unresolved or unavailable evidence, rather than measured failures. Auxiliary measurements can suggest follow-up tests but do not establish qualification or SU; keep them specific to their measurement source and do not average differently calibrated measurements into one reward.

Cite supplied evidence identifiers, use only available action families and permitted settings, and return one JSON object. Explain the observed signal, its interpretation and the proposed follow-up in three short reasoning_trace fields. These are reported explanations, not private model reasoning.

The campaign-level prompt objective used elapsed worker-slot time, including idle slot time; route comparisons used recorded full-route costs (Table S6). Before interpreting absence of a new SU as poor performance, the prompts suggested roughly 0.5 worker GPU-hours for low-cost, directly scored Complexa routes and 1.5–2 for delayed or higher-cost routes, including BindCraft, BoltzGen, ProteinMPNN redesign and Complexa MCTS. These were cost and feedback-delay guidance, not measured runtimes or hard timeouts.

###### Planner system instructions (condensed)

Propose one to four testable HypothesisCards from the latest evidence. Each card states a claim, cites evidence, proposes action families and settings, predicts qualification-measurement changes and specifies measurements to preserve. R/E/X affinities describe the purpose of the follow-up test; they do not assign a fixed method to each priority.

Predict changes only in pLDDT, iPAE or binder scRMSD: pLDDT increases, whereas iPAE and binder scRMSD decrease. Use the same ordered baseline references for all predicted changes in a card. Parent-bound jobs need concrete result identifiers; family summaries cannot substitute for parent designs.

Match the test to the observed weakness: sequence redesign or sequence hallucination for binder scRMSD; interface or backbone search for iPAE; and regeneration or pLDDT-directed search or refinement for low pLDDT with a plausible interface. Use past test outcomes and negative evidence to avoid repeating an unchanged, unproductive configuration. Preserve the other qualification measurements and obey registered setting bounds and workload limits.

The Planner’s L0–L4 ordering was a preference, not a required sequence: L0, obtain initial coverage of generator roots (Section C.2.2); L1, change search or sampling settings within a family; L2, address a near-miss design or an unproductive parent–child line through parent-bound redesign or a justified same-family alternative; L3, adjust scalar reward weights with criterion-level evidence; L4, expand higher-cost search when recent productivity or repeated near misses justify it. L1/L2 changes tested a different scientific hypothesis rather than only increasing workload or repeating reward adjustments. The system prompt also explains action-family roles and diagnostic-to-setting relationships; the dynamic input supplies current availability and bounds.

**Planner dynamic user prompt (schematic).**

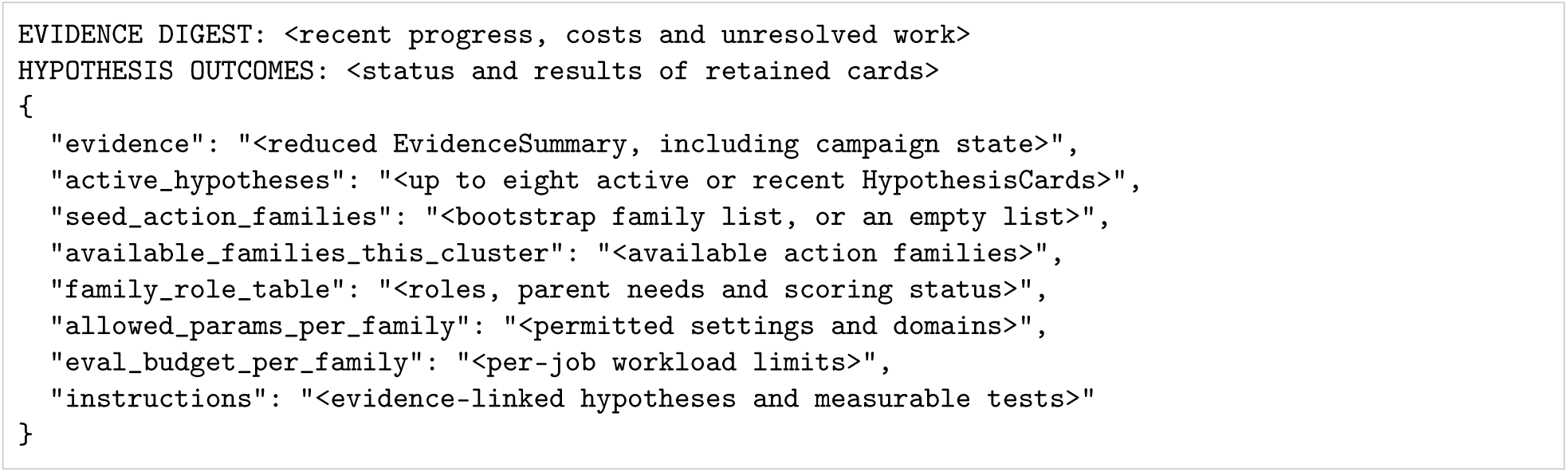

When provided, bootstrap families guide initial coverage. Prior deterministic advisory flags can be appended as recent_critic_flags_last_tick; despite the field name, these are not an additional Critic LLM call (Section C.5.4). Their presence depends on the call context. The prompt is an input for proposals, not permission to modify the archive or execute jobs.

###### Supervisor system instructions (condensed)

Use the current evidence, HypothesisCards and valid candidate jobs to propose R/E/X allocation proportions and optionally rank candidate jobs. Allocation values must be finite, nonnegative and sum to one; they concern future generation and redesign starts, not GPU-time fractions. Rank only supplied candidate identifiers. Each returned candidate belongs to one priority and has a unique global rank from 1 to the number returned; rank 1 expresses the immediate next-worker preference across priorities.

Prefer recent new SU per full exact-route cost. Among routes with reliable, approximately comparable recent efficiency, prefer the higher lower-quartile qualification-margin score, then its median (Section C.4.3). Treat sparse counts cautiously; quality does not compensate for a substantial efficiency disadvantage. Inspect source-specific diagnostics to select an evidence-supported follow-up, without counting native generator acceptance as qualification.

Keep proposed proportions consistent with immediate job priorities. Use campaign state, structural duplication, time without new SU, pending work and confirmed starts to interpret current needs. A job selected but not started is pending execution, not a failed scientific test. Required AF2 evaluation follows deterministic rules and is outside the LLM’s R/E/X allocation.

**Supervisor dynamic user prompt (schematic).**

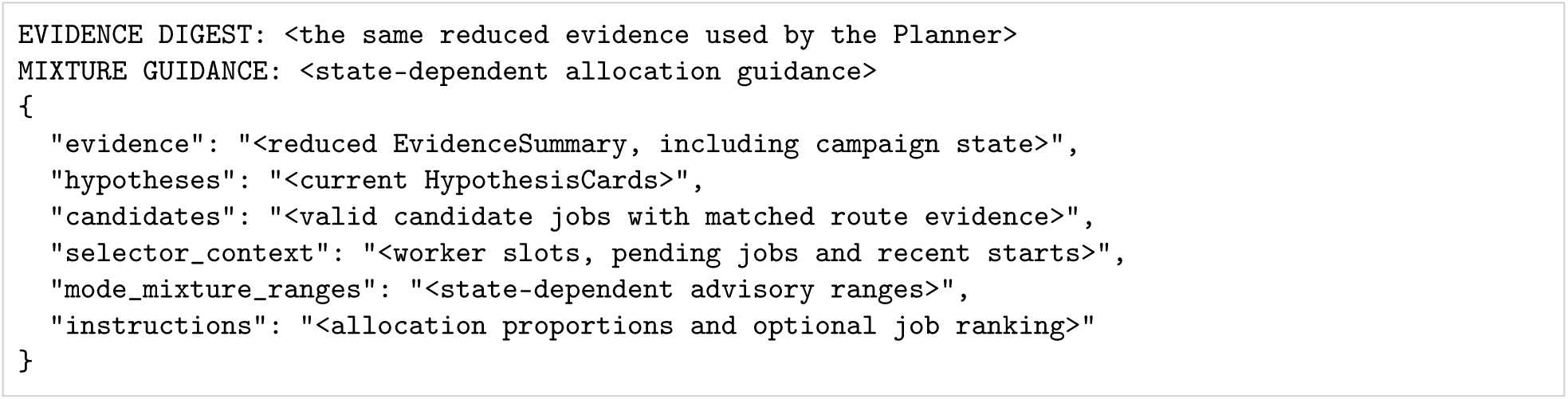

**Conditional Supervisor corrective message (template excerpt).**

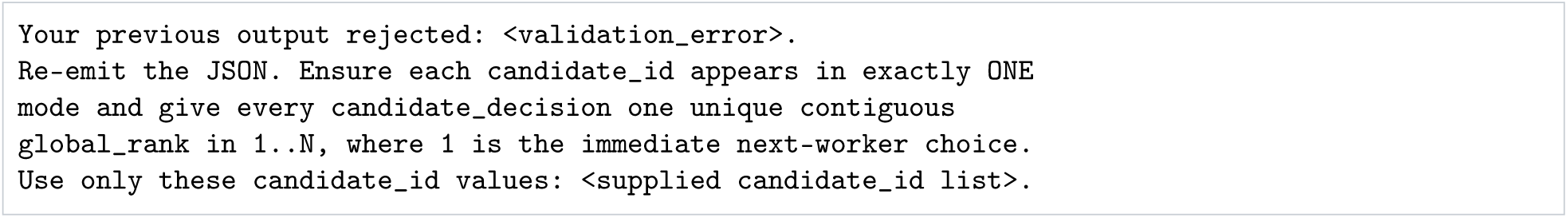

This corrective message can be sent once for the specific ranking/reference errors described under Parsing and validation in Section C.5.2. It follows the original user prompt and rejected response, with the same Supervisor system prompt. Other unusable responses follow deterministic repair or fallback; the primary campaign controller uses the Planner and Supervisor LLM roles, rather than a separate Critic LLM.

**Prompt guidance and execution rules.** Quality and cost preferences are specified in Section C.7, using the scores in Section C.4.3. Campaign-state labels inform proposals, while candidate eligibility, allocation bounds and worker starts follow the rules in Sections C.4.5, C.6, C.7.1 and C.8. Prompts impose no fixed generator hierarchy and do not guarantee that a proposed job starts. Archived proposals and decisions record the responses; prompt instructions alone do not establish compliance.

#### C.5.4. Call conditions and advisory checks

The Planner’s eight-card memory used the latest record for each card, prioritizing active cards and then recent supported or refuted cards; retired cards were excluded. Invalid, empty or abstaining output supplied no new cards, so construction used retained cards and candidates introduced by fixed rules. Refuted or retired cards could not generate candidates. Section C.9 defines card status and lifetime. The Supervisor was called with a non-empty card list and valid candidate pool; otherwise the controller recorded why it skipped the call. Invalid or abstaining responses used fallback (Section C.7.1).

**Deterministic advisory check.** When enabled, the check ran on valid, non-abstaining Planner output containing cards, before fallback to retained cards. It flagged proposed families whose prespecified runtime estimate exceeded the remaining wall time. It also flagged a configuration matching a retained failure recipe from the same family on at least two shared settings, including one beyond workload controls. All shared values had to match, the recipe needed at least two failed descendants, and its hash could not also appear among retained qualified recipes.

The check left current cards, candidates, selection and starts unchanged. Flags were stored as a zero-token, zero-latency LLMCallRecord and could inform a later Planner call; a subsequent clean record cleared earlier flags. Table S15, part A, reports these deterministic checks separately from model calls.

#### C.6. Candidate construction and validation

##### C.6.1. Validation of proposed jobs

Check + Build converts HypothesisCards into candidate jobs by resolving action families, proposed settings and required parents against the action registry and archive. Validation retains permitted settings, removes unsupported entries, clips numeric values to registered bounds and reduces oversized workloads when repair is possible (Table S5). Unrepairable jobs remain ineligible. Parent-bound redesign and evaluation require a resolvable structure with valid target and binder chains. Deterministic rules can then add or replace settings as described below; executor defaults fill only fields left unspecified. Section C.8 describes the subsequent queue and start-time checks.

##### C.6.2. Complexa configuration additions and replacements

**Measurement-directed defaults and refinement.** The *primary measurement* is the measurement named by the first predicted change in the HypothesisCard (Section C.5.2); it defaults to iPAE if no change is listed. A card with predicted changes but no suggested settings receives sequence hallucination for primary binder scRMSD, or adaptive sampling noise for primary iPAE (sc_scale_ noise=0.30 if no adaptive value is available). A proposed configuration emptied by validation does not receive this default. Greedy optimization enabled, a positive greedy-iteration count or a specified greedy_percentage selects refinement_algorithm=sequence_hallucination. For nonempty configurations without reward weights, primary iPAE or binder scRMSD also selects this refinement in stalled, deep-stall, near-miss-enriched, productive-with-duplication or structural-duplicate-collapse states, or when trusted near misses are present. State and clustering-reliability rules are defined in Sections C.4.5 and C.4.2.

**Reward-weight changes.** For reward checks, a *substantive search change* is a specified sampling-noise, steering-temperature, beam-width/branch, sample-count, replica-count, MCTS-search or sequence-refinement control. Values equal to explicitly registered workload defaults do not count; diffusion steps, batch size, filter limit or reward weights alone are insufficient. The builder uses the largest observation count in the primary measurement’s group, across qualification and auxiliary summaries. The groups are fold confidence for pLDDT (qualification pLDDT, auxiliary binder pLDDT and design pTM), interface confidence for iPAE (iPAE, minimum iPAE, mean/maximum ipSAE, ipTM and design-to-target ipTM), and geometry for binder scRMSD (scRMSD, shape complementarity, contact density, hotspot RMSD and buried surface area). Other primary measurements use their own counts.

Below 32 observations, reward weights are removed unless the proposal is Complexa MCTS with a substantive search change already present. If removal leaves no such change, the ordered additions below are attempted. With at least 32 observations, reward weights remain, but a proposal lacking a substantive search change receives one when available. Section D.2.2 shows a proposed weight removed by this check. Separately, Complexa best-of-*n* configurations with more than four replicas have batch size reduced to eight if it exceeds eight, retaining the requested sample and replica counts to limit memory use.

**Adaptive sampling noise.** Numeric additions use a fraction *f* of the registered range [*L*, *U*], giving *L* + (*U* − *L*)*f*. Values are rounded to the nearest integer, with ties to even, when both bounds are integral and *U* − *L* > 1; otherwise to four decimal places. The resulting configuration is checked against the same domains and workload limits. For sc_scale_noise, *g* denotes fold confidence, interface confidence or geometry, using the measurements listed under Reward-weight changes. Within each group, *F_g_* is the largest fraction of observations labeled fail for any measurement; *S_g_* is the largest fraction labeled pass or near-pass. Each fraction uses that measurement’s observation count as its denominator. The two maxima can come from different measurements and need not sum to one. Qualification summaries take precedence over auxiliary summaries for the same measurement; missing or zero-count summaries contribute zero.

Let *r*_MPNN_ be the largest auxiliary diagnostic-improvement score among ProteinMPNN redesign rows in route_values, or zero if absent (Section C.4.3). These substituted zeros disable evidence contributions rather than recording failures. If any *F_g_*, *S_g_* or *r*_MPNN_ is positive, the first matching row sets the initial fraction:

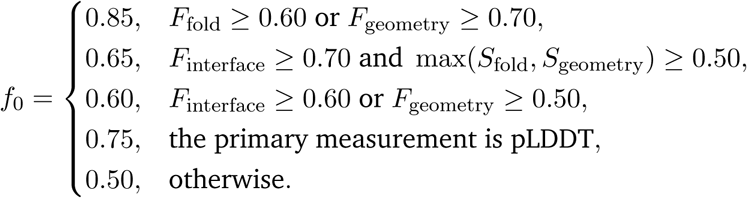

The fraction is capped at 0.50 when *r*_MPNN_ ≥ 0.50, or at 0.60 when 0.35 ≤ *r*_MPNN_ < 0.50. If max*_g_ S_g_* ≥ 0.70 and max*_g_ F_g_* < 0.50, it is additionally capped at 0.45. Without a positive measurement or redesign signal, the fraction is 0.55, increased to 0.65 only in deep stall or after at least 12 worker GPU-hours without a new SU, using the stall-time signal in Section C.4.2.

**Ordered search-setting additions.** When the reward check requires a search change, the following rules add permitted settings that are still unspecified, without overwriting accepted values. Duplicate pressure means a productive-with-duplication or structural-duplicate-collapse state (Section C.4.5). In steps 3 and 4, a beam-width addition also attempts to add n_branch at fraction 0.33; step 5 changes beam width alone.

1. For Complexa MCTS, add n_simulations, exploration_prob and exploration_constant at fractions 0.20, 0.57 and 0.33, respectively, then stop.
2. For FK steering, add temperature at fraction 0.45 under duplicate pressure and 0.30 otherwise; for primary iPAE or binder scRMSD, also add the adaptive noise above, then stop.
3. For the remaining Complexa families under duplicate pressure, first try beam width at fraction 0.43, then replicas at 0.60; stop after the first successful addition.
4. For primary iPAE, try beam width at fraction 0.43, then adaptive noise. For primary binder scRMSD, try sequence hallucination and adaptive noise together. For primary pLDDT, try beam width at fraction 0.35, then sampling noise at fraction 0.35. Stop if the applicable branch adds a setting.
5. If none applies successfully, try beam width at fraction 0.35, replicas at 0.50 and sampling noise at 0.45 in that order, stopping after the first addition.

##### C.6.3. Additional candidates generated by fixed rules

**Prespecified MCTS probe.** The builder can supplement card-derived jobs with the candidates below. These additions remain subject to validation and selection; availability does not force a launch. Outside the low-evidence state, it can add a Complexa MCTS configuration adapted from Proteina-Complexa’s inference-time search setup for difficult targets (Didi et al., 2026b). The rule requires at least six worker GPU-h without a new SU and either (i) a design that fails pLDDT but passes iPAE, or (ii) failure of at least half of standardized pLDDT or auxiliary binder-pLDDT observations, together with a pass-or-near-pass rate of at least 25% for standardized iPAE or auxiliary minimum iPAE, ipTM or mean ipSAE. It uses the stall-time signal in Section C.4.2.

The probe uses 400 diffusion steps, four samples, batch size 16, 20 MCTS simulations, exploration probability 0.5, exploration constant 1.0 and filter limit 100. Its iPAE and pLDDT reward weights are −1.0 and 1.0; sequence_hallucination uses greedy optimization, 15 iterations and greedy_percentage=5. The same configuration and trigger apply across targets. Additions stop after at least 3 route GPU-h and two ResultRecords with ok or no_artifacts status if the MCTS summary has no SU, no recent near miss, auxiliary diagnostic-improvement score below 0.35 and no pending standardized AF2 evaluation. This check uses the family summary, or the highest-cost exact route if that summary is absent.

**Previously evaluated configurations.** If no valid candidate offers parent-bound ProteinMPNN redesign or AF2 refolding with changed model, recycle or initialization settings, the builder can reintroduce up to three archived generation configurations. It uses exact-route rows, excluding untried, evaluation-only and collapse-risk routes and configurations already represented. Routes with SU or near-miss evidence precede those with only auxiliary progress. Their SU/GPU-h rate is the first available rate from the 3-GPU-hour, 6-GPU-hour or eligible 60-record window, followed by the lifetime rate. Without recent SU or near misses, a route needs at least 0.10 SU/GPU-h and is excluded after at least 1 route GPU-hour in any recent window. After 12 worker GPU-hours without a new SU, it is also excluded if only lifetime evidence remains, or its rate comes from the 60-record window without a rate or positive SU count from either cost-based window. Window and route definitions are in Sections C.4.1 and C.4.4.

A route lacking SU and near-miss evidence can instead be reintroduced when its outputs require separate AF2 evaluation, none remains pending, its auxiliary diagnostic-improvement score is at least 0.35 and accumulated route cost is at most 6 GPU-hours. Its recent-productivity status must not be dry_low_quality, dry_duplicate or collapse_risk. General-status defer routes are eligible only through this auxiliary-score rule. For both paths, 60-record rates are used only when the route has neither recorded downstream AF2-evaluation cost nor an AF2 qualification role.

**Candidates from other generator roots.** The *dominant root* is the generator root (Section C.2.2) with the most accumulated generator-route GPU time, using exact-route rows when available and family summaries otherwise; the lexicographically larger root name breaks a tie. For this candidate-availability rule, current SU evidence means a positive SU count or rate in any route or family summary for that root in the 3- or 6-GPU-hour window. A 60-record count or rate also qualifies before 12 worker GPU-hours without a new SU, but only for routes without downstream AF2-evaluation cost or an AF2 qualification role. These windows and costs follow Sections C.4.1 and C.4.2.

When feasible, at least two other roots are represented among valid candidate jobs and queued or running jobs if any of the following holds:

1. The state is low evidence or stalled, cumulative and recent SU counts are zero, and at least 0.10 recorded worker GPU-h and one ResultRecord have accumulated; a dominant root exists and lacks current SU evidence.
2. The state is deep stall and the dominant root lacks current SU evidence.
3. The state is stalled or structural duplicate collapse, at least six worker GPU-h have passed without a new SU, and the dominant root lacks current SU evidence.

In either productive state, the smaller requirement is one other root when recent SU gain is positive, at least four of the preceding ten planning updates were deep-stall updates with zero recent SU gain, and cumulative throughput is at most 0.25 SU per worker GPU-h. Recent gain uses the 60-record window (Section C.4.2). An additional root must have an untried or under-tested family, a family awaiting AF2 evaluation, or a family with SU, recent near misses, positive SU/GPU-h or auxiliary diagnostic-improvement score at least 0.35. The time-without-new-SU conditions use the stall-time signal, whereas cumulative throughput uses recorded worker cost (Table S6).

#### C.7. Supervisor prioritization and deterministic selection

**Supervisor ranking instructions.** The Supervisor ranks candidate jobs across **Rescue**, **Explore** and **eXploit** and proposes their desired allocation proportions. It is instructed to compare exact routes by recent SU/GPU-h, including required evaluation costs. When the lower of two rates is at least 85% of the higher, it should prefer the higher 25th percentile, then median, of the online quality score *q* (Section C.4.3). The prompt asks for caution with few observations or little compute; it imposes no additional minimum sample count for this comparison. Outside that rate range, quality does not override efficiency. A lower-cost route should precede a more expensive route when it yields comparable or greater SU/GPU-h with adequate quality.

The prompts also request more diversification after six worker GPU-hours without new SU and discourage treating near misses as sufficient progress after twelve. Following limited recovery from sustained deep stall, they request continued consideration of another generator root. These are instructions to the LLM; Sections C.6.3 and C.8.2 specify the separate candidate-availability and concurrency rules. Section D.2.2 shows an archived prioritization example.

##### C.7.1. Response handling and allocation bounds

**Response handling.** Table S8B distinguishes a usable Supervisor global order, valid allocation proportions without a global order, and fallback. A propagated fallback reason replaces the Supervisor proportions with the state-dependent values in part A and disables its global order. Retained cards can still supply candidates. If the response retains individual candidate decisions, their within-priority ranks and resource classes can still enter the fixed ordering described below. Candidate-specific failures affect that candidate alone. Confidence recording follows Section C.5.1.

**Allocation bounds.** Desired proportions are normalized and adjusted using either the basic or conditional bounds in Table S8C. Conditional bounds apply on fallback, in stalled or structural-duplicate-collapse states, when structural concentration is at least 0.50, when reported Supervisor confidence is below 0.65, or when at least 30% of the latest four available LLM-call records indicate fallback or non-OK parsing. Missing confidence and deep stall alone do not activate them. These bounds constrain desired shares of subsequent confirmed starts; global ordering and candidate availability can change realized start shares and GPU-time fractions.

Structural concentration is the larger available dominant-cluster fraction in the latest 60 ResultRecords: among all records with an online structural assignment, or among qualified records with a qualified-cluster assignment. The qualified fraction is used only with at least four assigned qualified records; these are records, not four distinct SU. Under conditional bounds, concentration ≥ 0.50 in either productive state sets the **eXploit** ceiling to 0.60 and the **Explore** interval to 0.20–0.40, retaining other bounds. In the near-miss-enriched state, concentration ≥ 0.70 raises the **Explore** ceiling to 0.50 while retaining the **Rescue** floor. These thresholds differ from the 0.75 stalled-state test in Section C.4.5.

**Table S8.**
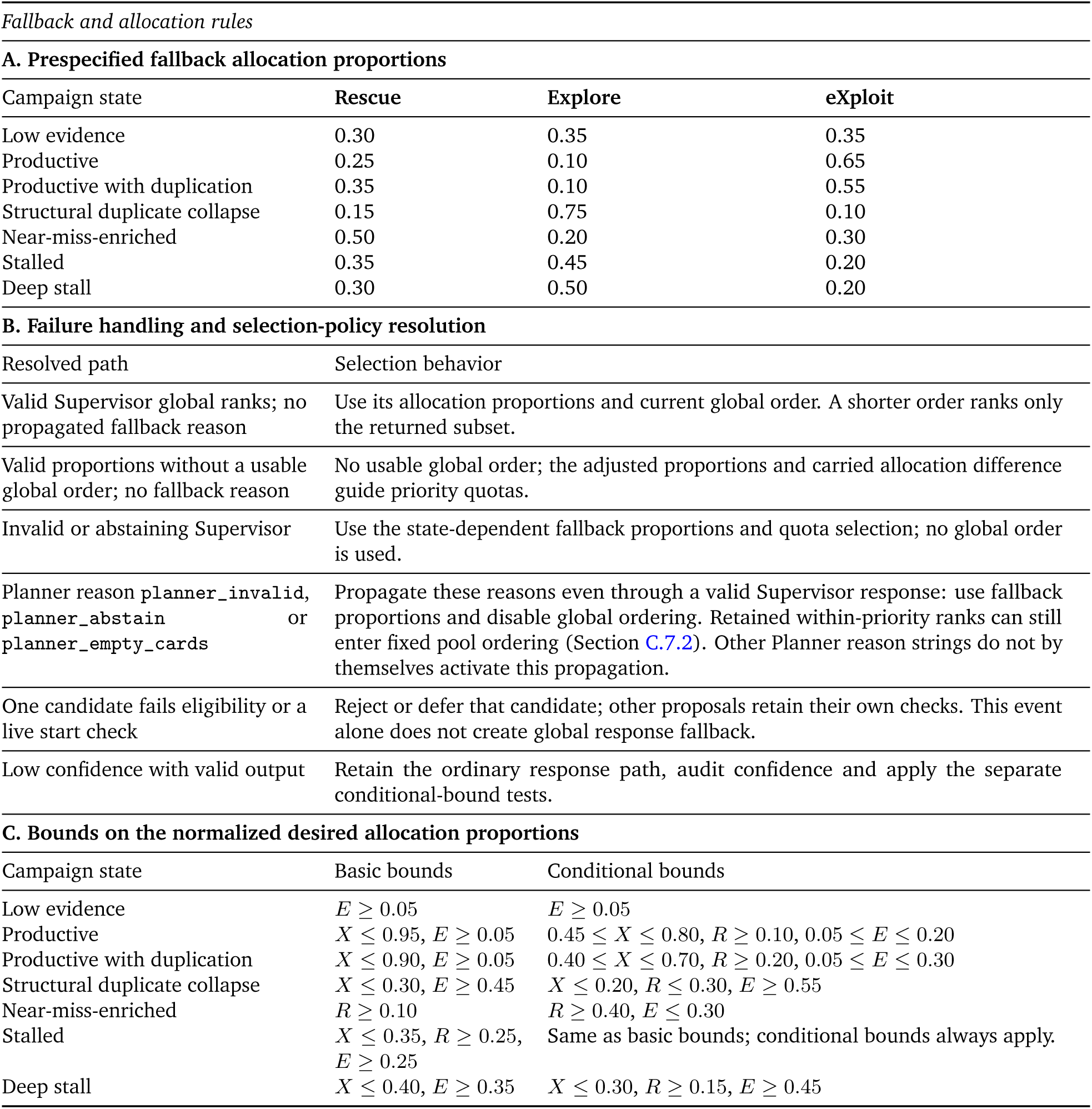
Fallback and allocation rules. Part A gives the initial desired R/E/X shares when selection takes the fallback path; Part B resolves the selection policy from Planner and Supervisor outcomes; Part C gives the basic and conditional bounds. The normalized shares satisfy *R* + *E* + *X* = 1. Shares refer to confirmed starts under R/E/X control (Section C.2.2), not success probabilities or guaranteed start-time or GPU-time fractions.

**Pending AF2 jobs and the Rescue-floor waiver.** For an update with *K_*_* additional queue positions, this check uses capacity 4 max(1, *K_*_*). It counts distinct selected or started AF2 candidate jobs without a linked ResultRecord. Recorded dispatch or parsing failures remove candidates, except deferrals caused by the high-cost running-job limit. At a count/capacity ratio ≥ 0.95, the **Rescue** lower bound is waived. This count is separate from ready-queue length and the unevaluated-design count (Section C.8.1).

The controller clips out-of-range proportions to their bounds and redistributes the remaining share proportionally among priorities not held at a bound; a zero remaining input weight gives equal shares. It repeats the correction for at most five passes. These adjustments change allocation targets without changing candidate validity. Queue capacity and worker availability remain separate quantities (Algorithm S1).

#### C.7.2. Candidate pools and ordering

**Global order and remaining positions.** A usable global order assigns unique positive ranks to the candidate identifiers returned by the current valid Supervisor response; it need not include every candidate. The selector considers that subset in global-rank order, then stable identifier order, subject to validation and high-cost limits. Family/route penalties and quotas do not reorder this subset. Remaining queue positions can be filled from other valid candidates using the fixed ordering below. Without a usable global order, allocation quotas determine how many positions each priority receives. Candidate priorities use Supervisor assignments when applicable and otherwise HypothesisCard affinities; candidates without an assignment can fill otherwise unused positions. Selection determines queue admission; ordering before worker starts follows Section C.8.2.

**Family penalties.** Family penalties place candidates later in quota selection and filling of remaining positions. They require qualified-structure clustering status ok with coverage ≥ 0.999; reports of no qualified designs or no structures also permit the check. Standardized AF2 evaluation is excluded. A positive recent family SU/GPU-h rate, direct or attributed through redesign/evaluation, removes the penalty; these 60-record method-summary rates are defined in Section C.4.1. Otherwise:

1. At least 3 recorded worker GPU-h with no direct or attributed SU and no trusted near miss gives a strong penalty.
2. At least 8 worker GPU-h gives a strong penalty when the best eligible comparison rate is at least 0.50 SU/GPU-h and the family’s lifetime rate is below either 0.25 SU/GPU-h or 35% of that comparison rate. A trusted recent near miss reduces this to a weak penalty.
3. If neither stronger rule applies, at least 6 worker GPU-h without recent SU or a trusted recent near miss gives a weak penalty.

An eligible comparison family has at least two credited SU and effective cost *g* ≥ 2 GPU-hours. Its unadjusted rate is the largest direct or attributed, recent or lifetime SU/GPU-h rate; *g* is its largest reported direct or attributed SU count divided by that rate. Multiplying the rate by *g*/(*g* + 2) reduces the influence of limited observations; the highest adjusted rate is the comparator above. Near-miss reliability follows Section C.4.2.

**Route penalties and deferral.** An exact route receives no penalty when its recent-productivity status is productive, productive with duplication or either delayed-productive variant. Otherwise, general status diversify gives a weak penalty; defer, collapse_risk, or recent-productivity status dry, dry_duplicate or dry_low_quality gives a strong penalty (Section C.4.4). On the quota path, a strong family penalty moves its candidates out of the initial pool if another candidate remains eligible for that pool. Strong route penalties then defer candidates when an alternative in the same priority remains. Under conditional allocation bounds, **Explore** candidates remain in the initial pool in both checks. Unfilled positions are considered first for remaining candidates, then family-deferred candidates, then route-deferred candidates, subject to the evaluation-backlog restriction below.

**Fixed ordering within the pools.** The ordering first compares recent family overrepresentation, family penalty and route penalty, in that order. A family is overrepresented when it supplies at least 70% of the latest ten available confirmed generation/redesign starts and either the state is structural duplicate collapse or it is productive with duplication with *f*_dup_ ≥ 0.75 (Eq. (S3)). After these penalties, unranked candidates use descending exact-route SU/GPU-h before the within-priority rank comparison; candidates with a Supervisor rank use route rate only to break equal ranks. Resource class, estimated cost class and stable identifier resolve remaining ties; cost classes are ordered low, diagnostic, standard and extended.

The route rate is the first available rate from the 3-GPU-hour, 6-GPU-hour or eligible 60-record window, then the lifetime rate (Section C.4.1). The 60-record rate is eligible only without downstream AF2-evaluation cost or an AF2 qualification role. For an unranked candidate with absent or zero route rate, a trusted recent family rate can substitute. Quota selection and filling of remaining positions initially limit a family to two candidates, increased to *K_*_* under basic bounds in productive or low-evidence states. This limit can relax to fill unused positions; the separate high-cost limits in Section C.8.2 still apply.

##### C.7.3. Allocation across starts and quota safeguards

**Carried allocation difference.** For adjusted proportions **p** = (*p_R_*, *p_E_*, *p_X_*), the controller reconstructs a carried allocation difference **d** from archived decisions and starts. At each archived update, it adds that update’s proportions multiplied by the larger of its approved generation/redesign launch-intent count and linked confirmed-start count, then subtracts one for each confirmed start from the corresponding priority. Each component is clipped to [−3, 3] after each update. An approved or queued job that never starts therefore does not count as a realized allocation. Global-order updates contribute their proportions and starts to the same reconstruction, although quotas do not control their immediate selection.

For a quota update with *K_*_* positions, temporary allocation differences are *d_m_* + *K_*_p_m_* for *m* ∈ {*R*, *E*, *X*}, clipped to [−3, 3]. Each position is assigned to an eligible priority with the largest difference; ties after rounding to six decimals favor the larger *p_m_*, then **eXploit**, **Rescue**, **Explore**. Each assignment subtracts one from that temporary difference. These provisional subtractions allocate the current quotas; subsequent reconstruction uses confirmed starts. Priorities without eligible candidates receive no quota, and unused positions can be filled through Section C.7.2.

**Maintaining productive computation.** On the quota path, one **eXploit** position can be preserved in productive, productive-with-duplication or near-miss-enriched states when stall time is at most 6 worker GPU-h, prior **eXploit** allocation has not been exceeded by more than one start, and an eligible candidate has positive route or recent family SU/GPU-h. Reused configurations labeled dry_duplicate without a positive 3-GPU-hour rate or a 3-/6-GPU-hour SU gain cannot qualify on route rate alone. Structural duplicate collapse prevents this safeguard. Stall time follows Section C.4.2.

With at least three positions, stall time at most 3 worker GPU-h and either campaign recent throughput ≥ 0.50 SU/GPU-h or an SU gain of at least two, the quota can preserve two **eXploit** positions. The gain is the increase above the previous highest cumulative SU count, or the 60-record gain when that increase is zero or unavailable. Both leading **eXploit** candidates must be outside the high-cost family, have low, diagnostic or standard estimated cost, avoid the reused-configuration exclusion above and have route or recent family throughput ≥ 0.50 SU/GPU-h. Required AF2 evaluations cannot supply these positions.

**Retaining a cross-generator candidate.** After ordinary selection, the quota path can retain one candidate added by the cross-generator rule in Section C.6.3. It fills an unused position or replaces a candidate that lacks a Supervisor within-priority rank of one. Such a candidate from BindCraft, BoltzGen or ProteinMPNN is not restored from the route-deferred pool when its family has at least 32 source ResultRecords without a linked standardized AF2 result and at least four with one, with no family or matching-route SU or near miss and no promising pending structure. This restriction applies both to the cross-generator safeguard and to ordinary filling from that pool. Start-time validity and capacity checks still follow Section C.8.2.

#### C.8. Evaluation admission and worker execution

Generation, redesign and standardized AF2 evaluation share the three worker GPUs. Evaluation admission determines which designs enter the common qualification protocol; queue management determines when their jobs start. Figure S2b and Algorithm S1 show these separate steps. Designs awaiting evaluation remain pending, without being counted as qualification failures.

##### C.8.1. Standardized AF2 evaluation: admission and capacity

**Eligible outputs and admission signals.** Section B.1.1 specifies which outputs can receive standardized AF2 evaluation. The controller schedules these evaluations alongside generation and redesign, using native measurements and previous route outcomes to prioritize inputs. These signals affect admission and ordering; qualification and SU credit still require the common evaluation rules. An *evaluation-source family* is the generation or redesign family that produced an evaluation input. A *native-like* BindCraft or Complexa output has generator-native pLDDT ≥ 90 on the 0–100 scale, normalized iPAE ≤ 7/31 and binder RMSD or scRMSD < 1.5 Å. A weaker *proxy-promising* signal is defined separately for each tool:

- **BindCraft:** a generator-accepted output, or an output with all three native measurements and at most one criterion outside the near-pass tolerances in Section C.4.1.
- **BoltzGen:** at least two of design ipTM ≥ 0.80, design-to-target ipTM ≥ 0.65, interface ipTM ≥ 0.65, minimum design-to-target PAE ≤ 7 Å, structure confidence ≥ 0.75 and native-refold RMSD < 2.0 Å.
- **ProteinMPNN:** global or local per-residue negative log-likelihood ≤ 1.20, with sequence recovery—the fraction of parent residues retained—either unreported or between 0.15 and 0.85, inclusive.

An input receives the highest evaluation priority if it is native-like or its exact route has status promote, a recent SU gain, a positive selected route rate, or a recent near miss together with status healthy (Section C.4.4). This input-priority test is broader than the route status promote. The rate follows the window preference in Section C.7.2, including lifetime fallback when recent rates are unavailable. Within each route, inputs satisfying this test precede proxy-promising inputs; the remaining inputs follow, with diversify and then defer/collapse_risk given lower priority. Native-score ranking and stable identifiers resolve ties. Eligible route groups are visited in turn, taking at most one input from each per pass.

**Cumulative allowance for each route.** Each exact upstream route initially receives an allowance of four standardized AF2 evaluations. The cumulative allowance can advance to 8, 16, 32 and 64, then double further as evidence improves. Advancing from four to eight permits at most four additional evaluations, subject to the capacity rules below. Native or proxy signals affect which inputs are evaluated first but do not bypass this progression.

Advancement uses the latest archived route summary’s count of returned standardized AF2 ResultRecords, including unsuccessful outcomes. At a completed boundary, the controller compares that summary with the most recent earlier summary whose count is no greater than the preceding configured boundary: zero for the four-to-eight decision, or at most four for eight-to-sixteen. At least one of six measures must increase: route-attributed SU, individual near-miss result count, qualified-cluster count, 25th-percentile or median online quality score, or auxiliary diagnostic-improvement score (Sections C.4.1 and C.4.3). Score increases must exceed 10*^−^*^6^; the diagnostic score is stored to three decimal places. Near-miss evidence requires the reliability check in Section C.4.2. Missing comparison values contribute zero to this test.

Without an increased signal, the allowance stays at the completed boundary; raw qualified-design count does not advance it. A partly completed block can finish its current allowance: five returned evaluations permit continuation to eight, not sixteen. Confirmed starts and evaluations already selected for admission are subtracted from the authorized total, preventing repeated admission from repeatedly advancing the same evidence. Above 64, the allowance continues doubling, but the comparison lookup still uses the largest configured boundary below the current count. The spare-capacity exception below can admit one further evaluation without advancing this allowance.

**Family-level admission limits.** An input is *evidence-backed* for these limits if it satisfies the highest-priority test above or is proxy-promising. Ordinary admission allows at most 24 currently unsupported evaluations per evaluation-source family, counting started and already selected jobs. A family’s total approvals under the same planning-update identifier are compared with a cap of four for an unsupported candidate or sixteen for an evidence-backed candidate; these are not separate additive quotas. Route allowances still apply to both groups.

**Limits based on recent starts.** Once at least 20 starts have been recorded, reservation compares the number of standardized AF2 starts among the latest 40 available starts with a limit of 24, reduced to 20 in productive-with-duplication or structural-duplicate-collapse states. Before 20 starts, the corresponding limits are 12 and 10. If this budget is exhausted, available native-like inputs permit an exception of at most two positions, or one in deep stall; otherwise an available input satisfying either the proxy-promising or highest-priority test permits one. These counts limit reservations, not the fraction of GPU time spent on evaluation.

**Queue reservations.** AF2 jobs can be inserted at the front of the queue before dispatch and again during queue refill. The passes use free workers *F* (M) and queue space *K*(Q), respectively; the first can temporarily extend the queue beyond its target depth. Both passes share a per-loop ceiling of one, increased to two when the pending-output count used for this ceiling reaches 32. That count uses native-like or proxy-promising outputs when any are pending, otherwise all eligible unevaluated outputs. Each pass subtracts both earlier reservations in that loop and currently running AF2 jobs from the ceiling (Algorithm S1). An AF2 job already in the queue prevents another reservation.

Each pass also checks its available positions and eligible candidate jobs. The local limit is at most half those positions rounded up and leaves one unreserved when two or more are available. Its one-to-two increase requires 32 eligible candidates in that pass, so each pass admits at most one with the three-worker setup. During deep stall, the shared ceiling is one when an eligible output still awaits AF2 evaluation; native-like evidence is not required. Excess queued AF2 reservations are cancelled when the refreshed state enters deep stall. A single free position can be used for evaluation when an unevaluated output exists; otherwise that position is left for generation or redesign. These rules reserve queue positions rather than assigning a GPU permanently to evaluation.

**Filling spare queue positions.** After generation and redesign selection, one AF2 job can be appended if queue space remains, the state is not deep stall and no standardized AF2 job is queued or running. This path first uses ordinary route allowances. If they provide no candidate, it can select one from a route whose allowance is exhausted, bypassing the cumulative unsupported-family limit as well. The per-update family limits and start checks still apply. This exception uses spare queue capacity; it neither establishes improved evidence nor doubles the route allowance.

##### C.8.2. Queue admission, starts and result collection

**Queue refill.** Candidate ordering and allocation follow Sections C.7.2 and C.7.3, with validation in Section C.6. Queue space *K*(Q) and free workers *F* (M) are defined in Algorithm S1. Planning can refill the queue while all workers are occupied after the first target ResultRecord arrives; it pauses beforehand if every worker is busy. Queued jobs can start before new planning calls and again after queue refill, at the two dispatch points in the algorithm.

**Queue order and new evidence.** Before dispatch, required AF2 jobs come first. Generation and redesign jobs then follow their recorded annotations: recent SU or near-miss evidence, diagnostic improvement, ordinary proposals, and finally reused configurations supported only by lifetime evidence. Within a group, jobs with selection records precede those without them; newer selection updates come first, with existing queue order breaking ties. This ordering uses annotations in candidate and selection records rather than recomputing route scores. New results or a state change alone do not remove ordinary queued jobs. The exception is jobs selected in low evidence before any result existed: once the first result arrives, these queued jobs are removed so planning can use the new evidence.

**BindCraft concurrency limit.** BindCraft is the high-cost family, with an ordinary limit of one running job. When selection has more than one queue position to fill, at least one position is withheld from BindCraft; this bound applies even if no alternative is available. The three-worker pool likewise permits at most two simultaneous BindCraft jobs.

For this concurrency check, current route evidence means a positive SU count in the 3- or 6-GPU-hour window, a positive 3-GPU-hour rate or a trusted recent near miss. Rows with neither downstream AF2 cost nor an AF2 qualification role can also use the 60-record SU count and, when the 3-GPU-hour rate is missing, its rate (Section C.4.1). The limit can increase from one to two under any of these conditions:

1. **Family evidence:** a positive recent direct or attributed SU/GPU-h rate or a trusted recent near miss in the method summary (Section C.4.1); alternatively, lifetime SU or near-miss evidence with a lifetime rate ≥ 0.25 SU/GPU-h, provided that evidence has not become stale.
2. **Route evidence:** the family’s highest-rate route or family summaries provide at least one SU, at least 1.0 route GPU-h and current SU or near-miss evidence. Their rate, taking the larger current or lifetime value, must be at least 0.10 SU/GPU-h and 80% of the best current rate among competing summaries with current evidence. For tied highest-rate summaries, the largest count and cost and the presence of current evidence are used.
3. **Testing an alternative during a stall:** at least 32 ResultRecords, at least 0.75 worker GPU-h without new SU, and a best recent non-evaluation-family rate below 0.25 SU/GPU-h, in low evidence, stalled, deep stall or structural duplicate collapse.
4. **Repeated prioritization with delayed starts:** at least six BindCraft entries in archived Supervisor candidate decisions, a start-to-entry ratio ≤ 0.35, and at least one capacity-deferred or selected-but-unstarted job. The state must be low evidence, stalled, deep stall or structural duplicate collapse, or the campaign must have fewer than four SUs. Lifetime evidence must not be stale.

Lifetime evidence is stale when neither family nor route summaries retain current SU or trusted near-miss evidence and a previously productive summary has accumulated at least 1.0 recent route GPU-h. Recent windows follow Section C.4.1; route rates follow Section C.7.2. SU-based promotion requires the clustering check described there, and near misses use Section C.4.2. Conditions 2–4 are blocked after at least 6 recorded family GPU-h and more than one timeout without direct or attributed SU or trusted lifetime/recent near misses. Raising the limit permits another start but does not select a job; queued jobs do not count as running jobs.

**Starts, deferrals and failures.** Approved LaunchDecisions permit queue admission; DispatchRecords record starts and execution events (Section C.3). Before a start, the controller resolves required inputs and rechecks current constraints. A concurrency deferral returns the job to the queue without consuming a retry. Other transient start failures are retried at most three times, for four attempts in total, after all currently free workers have been considered. Permanent failures and exhausted retries remove the job. DispatchRecords retain terminal start failures and distinct capacity-deferral reasons; intermediate transient retries need not each produce a record. A later runtime or output-collection failure does not erase an earlier confirmed start.

**Incremental and final result collection.** Running BindCraft jobs are checked for new outputs at the interval in Section C.8.3. Only previously unarchived ResultRecords are appended. Worker time since the preceding addition of results is divided among the new records; final parsing collects any remaining outputs. Thus a running job can already contribute recorded results and cost. Re-reading archived outputs does not append them again.

If final parsing yields no new design after earlier results, or no output at all, a synthetic completion record retains an uncharged remainder ≥ 0.005 GPU-hours or a failed outcome. It represents cost and outcome rather than another design. Successful re-reading of outputs already archived through another path adds no second charge. A successful, empty parse below that threshold instead records an output-collection failure without a ResultRecord. Final-parsing exceptions are recorded separately; after at least 0.005 elapsed GPU-hours, a failure record retains the full elapsed job cost, which can overlap earlier partial charges. Table S6 distinguishes these records from other compute-accounting quantities.

##### C.8.3. Worker time limits and failure handling

**Polling and planning failures.** Active controller iterations normally wait 5 s between polls; an iteration with no planned, queued, dispatched or running jobs waits 120 s. A cached campaign-state calculation can be reused for up to 60 s when result counts, worker availability, running-worker cost rounded down in 0.25-GPU-hour increments, and queued/running job counts by family are unchanged. A failed state check retains the previous state. A full planning-update exception preserves already appended records, waits 60 s and restarts the loop before further admission or dispatch. Incremental BindCraft collection runs at a default 600-s interval, bounded below by 300 s.

**Progress checkpoints.** A separate checkpoint saves elapsed campaign time and monitoring evidence at a default 60-s interval, including when all workers are occupied. When native-like or proxy-promising designs await standardized AF2 evaluation and a standardized AF2 job is running, the checkpoint recomputes current evidence and can update the campaign state. Otherwise, it reuses the latest archived summary and updates elapsed time. Checkpoints neither append an EvidenceSummary to the JSONL history nor call an LLM.

**Job timeouts.** Limits are elapsed seconds on one worker slot. Complexa Beam, best-of-*n* and FK jobs have 5,400-s ceilings; MCTS has a 9,000-s ceiling. Standardized AF2, ProteinMPNN and BoltzGen ceilings are 2,160, 900 and 10,800 s. When a limit is exceeded, the controller attempts to collect available outputs and terminates the unfinished job. These ceilings limit overruns rather than specifying expected runtimes or selection costs.

BindCraft instead uses a progress watchdog. After a 5,400-s grace period, the first check starts a no-progress timer. An increase in scoreable output count resets it; more than 5,400 s without such progress triggers termination. Growth of intermediate files alone does not reset the timer.

**Disabling repeatedly unsuccessful families.** A family becomes unavailable after at least two timeout ResultRecords and 3 recorded worker GPU-h without qualifying or near-miss metric evidence in its attributed results, provided another available generation family not requiring a parent remains. Standardized AF2 is excluded. The controller prevents further candidates from that family and removes its queued jobs, while running jobs continue. Disablement itself records neither a timeout nor a new worker start.

**Campaign time limit and shutdown.** The 48-h controller limit is checked at loop entry and includes saved elapsed time if a campaign resumes. Planning and dispatch can finish within an iteration that has already begun. Section B.3 defines the separate reporting denominator of 144 H100 worker GPU-h.

At shutdown, busy workers are visited in pool order and each receives up to 600 s to finish. Normal completion is parsed before visiting the next worker; a timeout triggers attempted output recovery followed by termination. The scheduler can end the allocation before all waits finish. Final endpoint inclusion follows Section B.3 rather than these runtime waits.

#### C.9. Hypothesis feedback

Hypothesis feedback tests the measurement changes predicted in *HypothesisCards* (Section C.5.2). At the end of each planning update, deterministic rules evaluate previously active cards using results available at the start of that update. Updated cards are appended under the same hypothesis identifier for subsequent planning (Section C.3), without changing qualification or SU credit. These tests assess numerical predictions, not causal explanations. In the worked example in Section D.2.2, eight descendants qualified but none supported the card’s specific predictions.

**Results and comparison baselines.** A result is linked to a card through the candidate job that produced it, including downstream evaluation jobs carrying the same hypothesis link. Each candidate uses one comparison baseline shared by all predicted measurements: the first resolvable result among the card’s ordered baseline references, followed by its cited evidence. Downstream evaluation can inherit this baseline; otherwise, a job without a recorded comparison baseline uses its execution parent. Thus the comparison baseline need not be the design supplied as input to the job. Results without a resolvable baseline are excluded. Counts are recomputed from all linked result–baseline pairs, rather than adding the same evidence again at each update.

**Predicted improvement.** Let *d*_baseline_ and *d*_result_ denote the measurement’s nonnegative qualification deficits (Section C.4.1). For *d*_baseline_ > 0.01, the relative deficit reduction

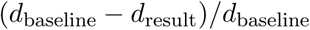

must meet the minimum specified in the card. When *d*_baseline_ ≤ 0.01, support instead requires the specified absolute improvement in the favorable direction, defaulting to 1.0 in that measurement’s units if the value is absent or zero. The 0.01 tolerance is applied in each measurement’s units.

**Result-level evidence.** A result supports a card when every predicted change meets its improvement requirement and no checked limit on worsening measurements is violated. For each such limit, a baseline deficit ≤ 0.01 must remain ≤ 0.01; otherwise, the result deficit must not exceed the baseline deficit multiplied by one plus the card’s permitted relative increase. A limit is checked only when both baseline and result measurements are available. Skipping a missing comparison does not establish that the measurement was preserved.

A result contradicts a card when every predicted measurement has a baseline deficit > 0.01 and a relative deficit reduction ≤ 0.05. Both support and contradiction require the predicted direction to match the measurement’s favorable direction. Missing predicted measurements prevent either finding, with one exception: a missing baseline measurement can receive support when the linked standardized AF2 result passes the full qualification rule. Other predicted changes and preservation limits still apply. Support and contradiction are counted independently; results satisfying neither test are neutral.

**Card status.** In the reported campaigns, each supporting result contributed one point; two points marked a card as supported. Otherwise, at least three evaluated result–baseline pairs and a contradictory fraction ≥ 0.70 marked it as refuted (archived as contradicted). Neutral records remain in that denominator. Support takes precedence if both thresholds are met. A card meeting neither threshold remains active until its specified lifetime is reached, counted from creation in planning updates (default ten; allowed range one to twenty), and is then retired. Expired cards with no linked results are retired before the Planner call; other status updates follow the end-of-update evaluation above. Supported, refuted and retired states are terminal.

### D. Supplementary experiments and campaign analyses

The analyses below examine throughput and robustness (Figure 2), archived follow-up decisions (Figure 3), allocation and recovery (Figure 4), and post-hoc confidence and structural diversity (Figures 2c and 5).

#### D.1. Throughput and robustness

##### D.1.1. Repeated campaigns

Across the three independently initialized campaigns per controller on CD45, SC2RBD and CbAgo, every T-REX campaign produced more final TM0.6 SUs than every PUCT or *ε*-greedy campaign on the same target (Figure 2d; Table S9). Section B.7 specifies the replication design. Figure S3 shows the online T-REX trajectories and separately marks the final recomputed counts; online counts can change when clustering is updated (Section C.4.2).

**Figure S3.**
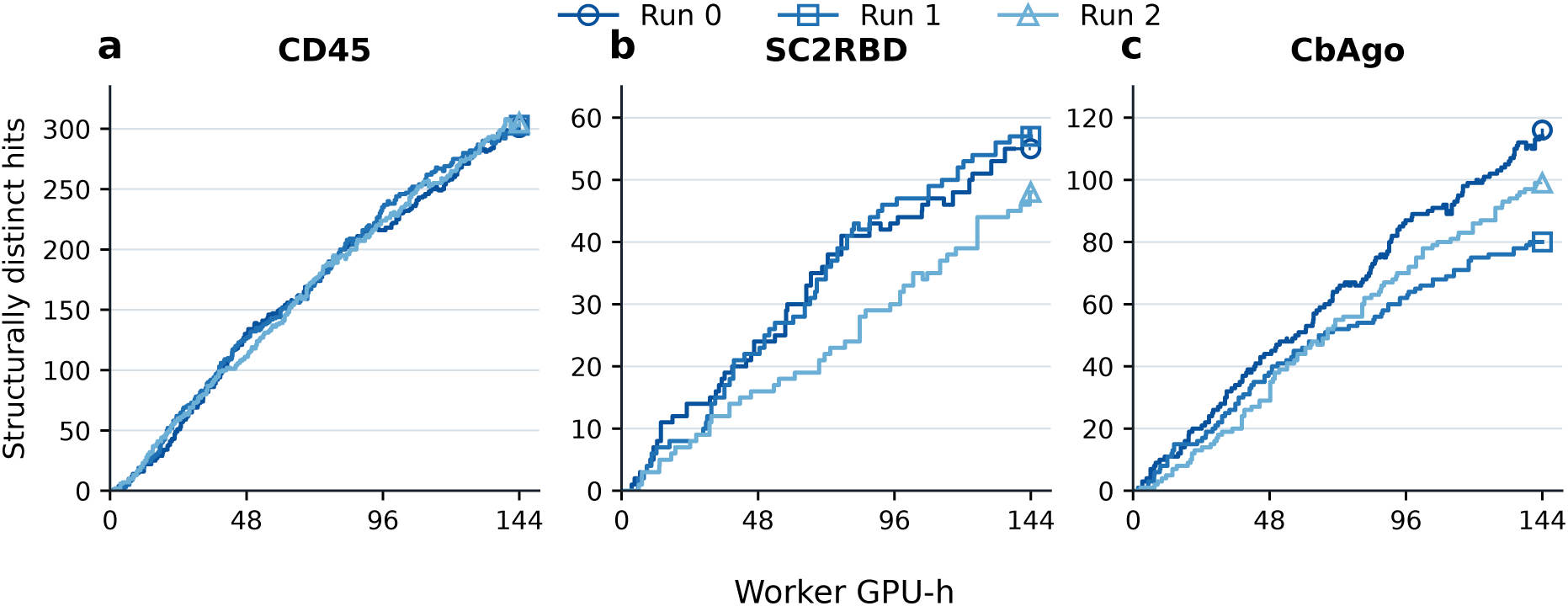
Online SU counts across three T-REX runs. Each panel shows three independently initialized campaigns for one target under the prespecified 144 H100 worker-GPU-h denominator. Solid steps show the archived online TM0.6 SU counts, which can decrease when Foldseek clustering is updated; open symbols show final recomputed TM0.6 SU counts. Figure 2d and Supplementary Table S9 compare the corresponding adaptive controllers.

**Table S9.**
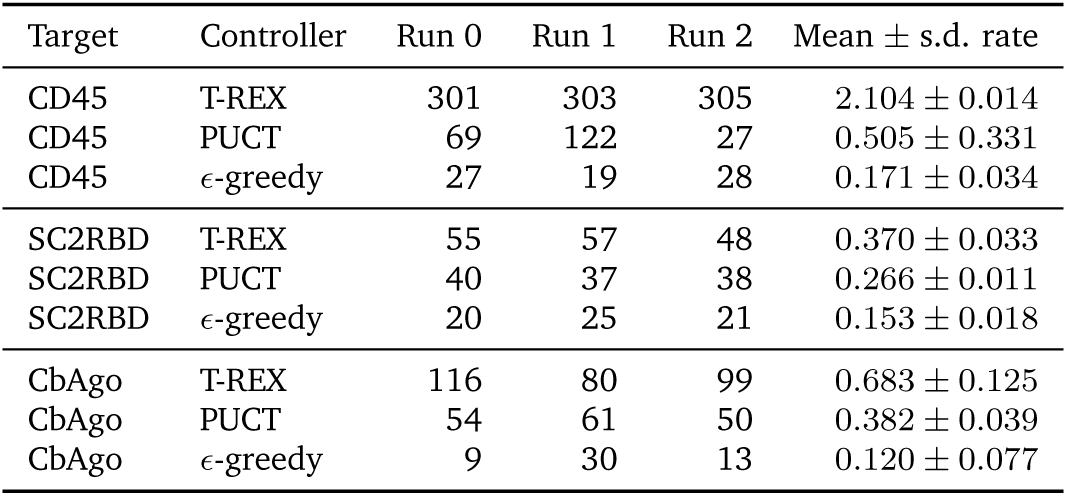
Repeated adaptive-controller campaigns. Final TM0.6 SU counts for three independently initialized campaigns of each adaptive controller (Run 0, Run 1 and Run 2 use controller seeds 0, 1 and 2, respectively) using the same campaign setup and prespecified denominator of 144 H100 worker GPU-h; the summary column is mean ± sample standard deviation (s.d.) after dividing each count by 144 worker GPU-h.

##### D.1.2. Retrospective checkpoints and shorter-budget comparisons

**Reconstructing accumulation curves.** The curves in Figure 2a use final qualification and TM0.6 cluster assignments for the designs included under Section B.3. Adaptive curves assign each cluster to the earliest archived *EvidenceSummary* containing a qualified member; generator-only curves use the completion events specified in Table S10. PUCT HER2-AAV records arriving after the last summary are assigned to the 144-worker-GPU-h cutoff. A final horizontal segment holds the last assigned count constant to that cutoff without adding clusters. These retrospectively reconstructed curves can differ from online cluster counts. Because the recorded events differ across methods, intermediate checkpoints describe accumulation rather than precise cross-method completion times.

At checkpoint *B* worker GPU-h, *N*(*B*) counts final clusters assigned an event at or before *B*, and *N*(*B*)/*B* gives the checkpoint rate (Table S10). Cluster assignments remain fixed; these are observations from campaigns run to 144 worker GPU-h, not campaigns independently stopped and finalized at *B*. At 10 worker GPU-h, T-REX had the largest count on six targets and tied BindCraft-only on HER2-AAV; it led on all seven at 50, 100 and 144 worker GPU-h. The across-target geometric-mean T-REX/PUCT count ratios were 3.94, 3.10, 2.73 and 2.43, respectively.

**Table S10.**
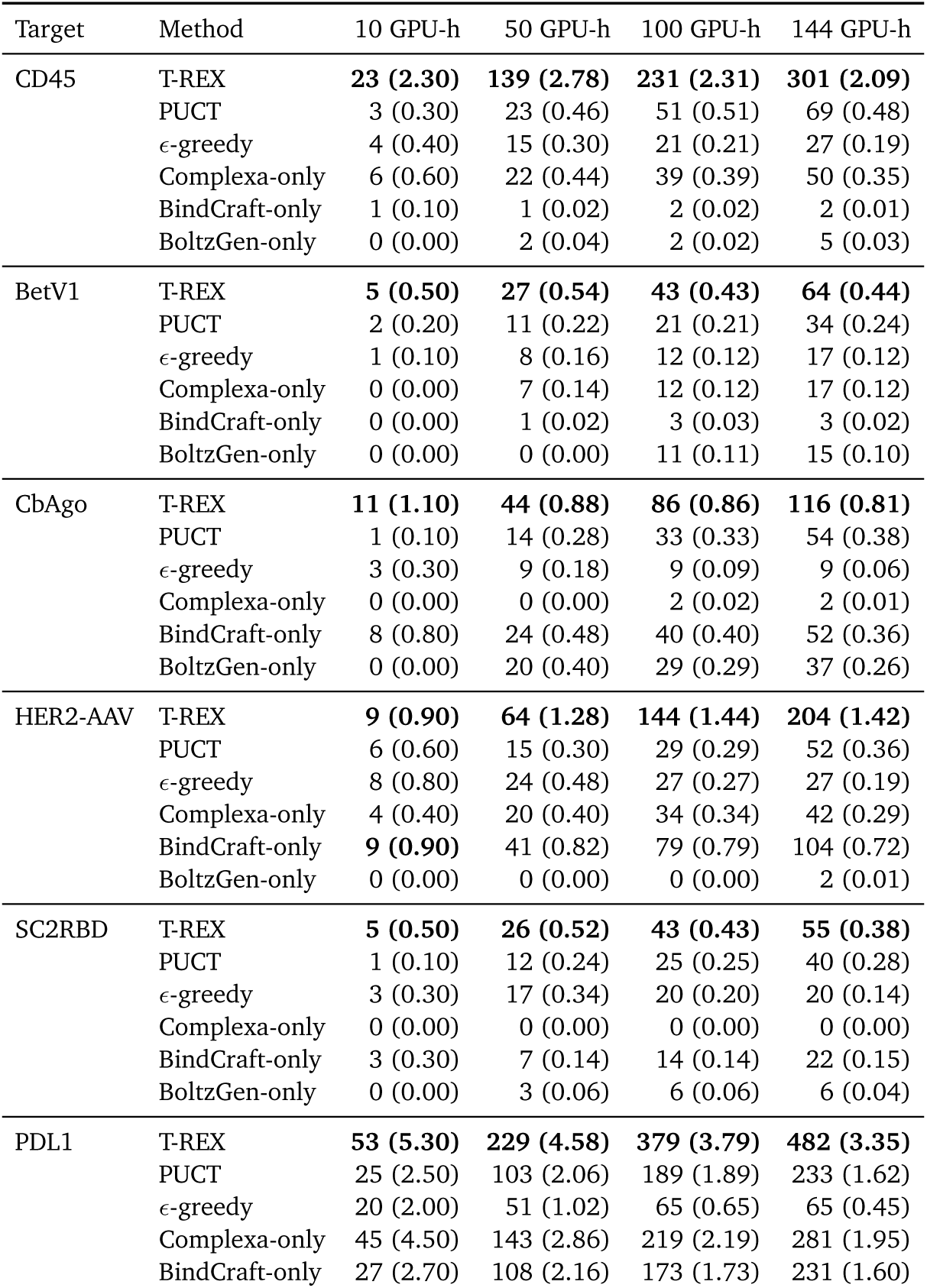
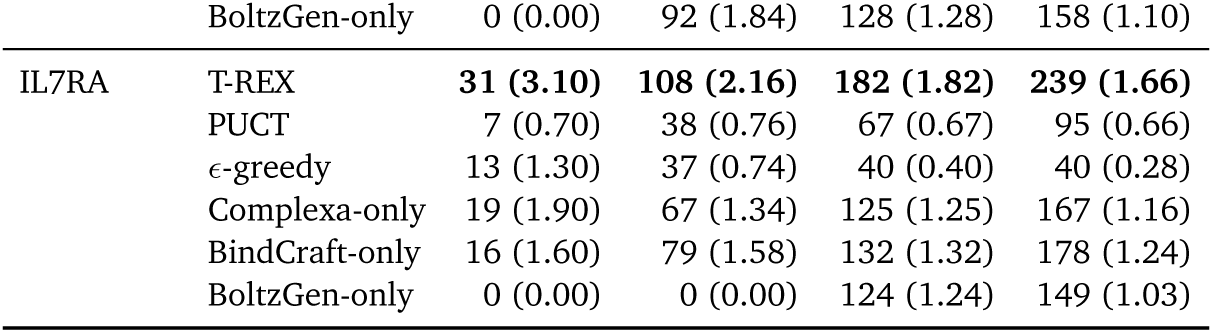
Retrospective TM0.6 structure-unique accumulation at fixed compute checkpoints. Entries give the cumulative number of final TM0.6 SUs assigned by the recorded curve event time, with SU per worker GPU-h in parentheses; bold marks the largest count within a target and checkpoint. The checkpoints are read from campaigns run to 144 worker GPU-h without new stopping, shutdown or clustering procedures at the earlier budgets. Adaptive curves use archived *EvidenceSummaries*; Complexa-only uses individual design completions; BindCraft-only uses output completions; and BoltzGen-only uses iteration completions. For the generator-only baselines, any required standardized AF2 evaluation occurred after the generation cutoff and is excluded from the generation budget. The event types differ across methods, so intermediate checkpoints do not measure comparable per-design completion times.

**Bayesian-optimization comparison.** The independently stopped 100-worker-GPU-h SMAC3 campaigns produced 329 final TM0.6 SUs across seven targets, compared with 1,108 at the retrospective 100-worker-GPU-h checkpoints of T-REX; T-REX had the larger count on every target (Table S11). SMAC3 stopped after approximately 25 h on four workers (setup in Section B.5.2); T-REX checkpoints come from three-worker campaigns planned for 144 worker GPU-h. The comparison matches worker compute but does not establish the outcome of independently stopping T-REX at 100 worker GPU-h.

**Table S11.**
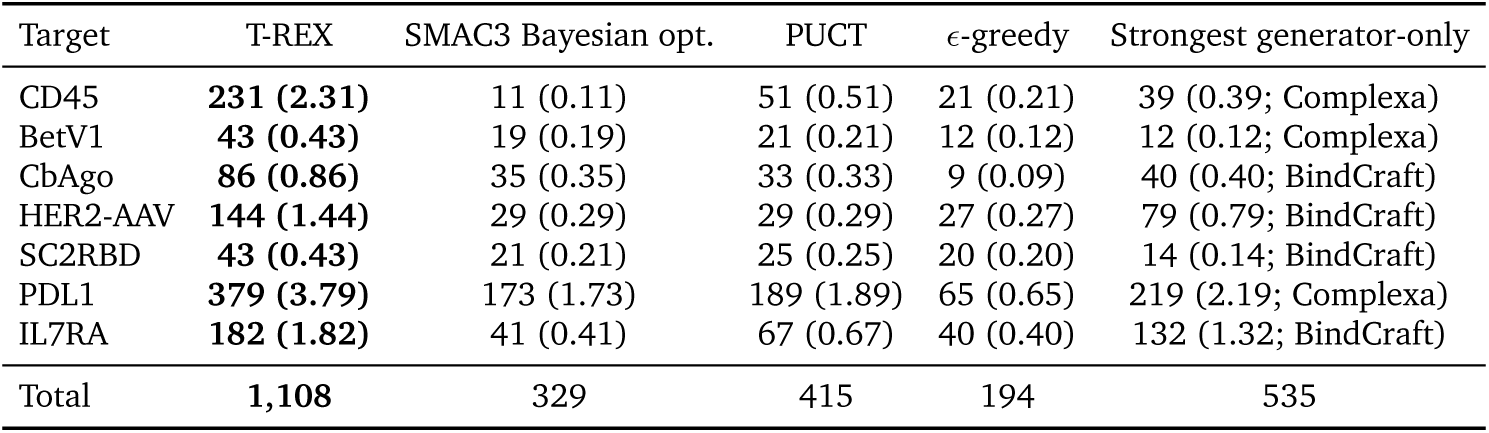
No-LLM SMAC3 Bayesian-optimization comparator at 100 worker GPU-h. Each target–method cell gives the TM0.6 SU count, with SU per worker GPU-h in parentheses. SMAC3 values are archived endpoints from independently stopped campaigns that used four H100 workers for 25 h and ended at 99.85–100.00 worker GPU-h. Values for T-REX, PUCT, *ε*-greedy and the strongest generator-only reference are retrospective 100-GPU-h checkpoints from Supplementary Table S10. The SMAC3 policy and execution setup are specified in Section B.5.2. The strongest generator-only column selects the largest of Complexa-only, BindCraft-only and BoltzGen-only at the same checkpoint, with the family shown in parentheses. The final row pools counts across seven targets. Bold marks the largest count per target.

##### D.1.3. Compute accounting sensitivity

**LLM call time and serving cost.** Across 3,125 archived model calls, end-to-end latency totalled 84,533 s (23.48 h; mean, 27.05 s per call), equivalent to 6.99% of the aggregate nominal campaign wall time (7 × 48 = 336 h). LLM serving used a separate GPU while worker jobs could continue, so this sum measures neither added campaign duration nor delays in queue refill or worker idle time. The primary worker-budget definition and endpoint-inclusion rules are given in Section B.3.

A total-compute sensitivity added the full 48-h LLM-server reservation to each campaign’s reconstructed worker time: 336 H100 GPU-h across seven campaigns, or 14.3 times the aggregate call latency. This charges approximately 192 H100 GPU-h per T-REX campaign while retaining its three-worker execution. T-REX remained ahead of PUCT on every target under this accounting (1.03–3.27-fold in TM0.6 SU per total H100 GPU-h; geometric mean, 1.82-fold; Table S12a).

A second sensitivity retained the full server charge and added archived post-cutoff AF2 evaluation time to each generator-only baseline before selecting the strongest adjusted baseline per target. Online T-REX evaluation was already included in worker time; Complexa supplied the common AF2 measurements inline and required no separate post-cutoff evaluation charge. T-REX again led on every target (1.07–4.52-fold; geometric mean, 1.90-fold; Table S12b).

**Table S12.**
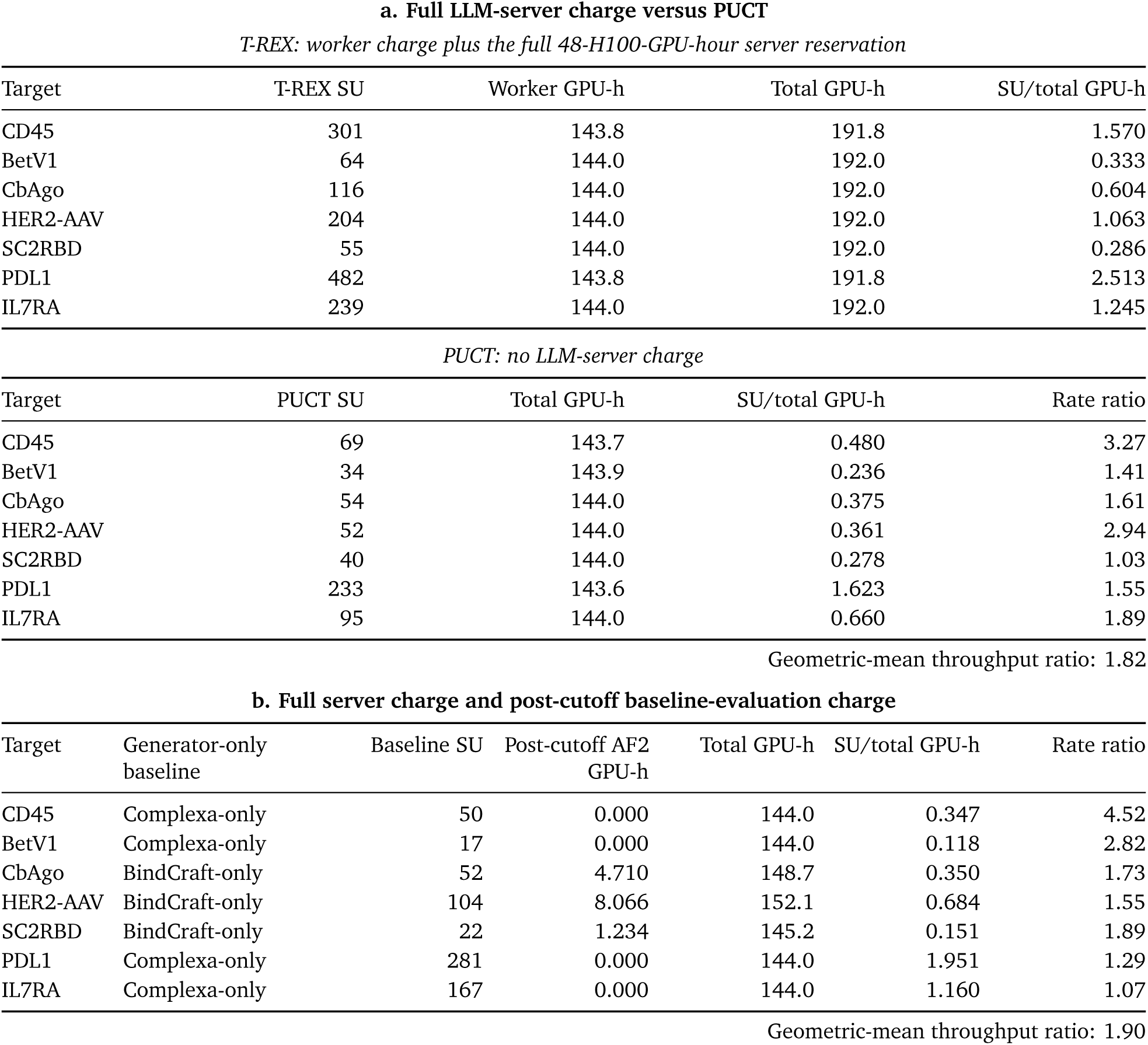
Compute-accounting sensitivities. a. , For T-REX, total charged time adds the full nominal 48 H100 GPU-h reservation of the separate vLLM server to reconstructed worker time; this is a total-compute charge, not a four-worker rerun. PUCT used no LLM server. **b**, The same full T-REX server charge is retained, and archived post-cutoff standardized AF2 evaluation time is added to every generator-only baseline before the strongest adjusted baseline is selected for each target. Online T-REX evaluation is already included in worker time; Complexa supplied the common AF2 qualification measurements inline and therefore had no separate post-cutoff AF2 charge. The T-REX counts, total charges and adjusted rates in part a also apply to part b. Rates use the primary TM0.6 SU counts. Rate ratios divide T-REX throughput by comparator throughput; values above one favor T-REX.

##### D.1.4. Diversity and qualification-margin sensitivity

T-REX retained the highest cluster throughput on all seven targets under both MMseqs2 sequence clustering at 70% identity and Foldseek structural clustering at TM0.5, TM0.6 and TM0.8 (Figure 2b; Tables S13 and S14). These analyses use the same qualified design pools and the clustering settings in Section B.2; sequence clustering is applied before structural deduplication.

**Table S13.**
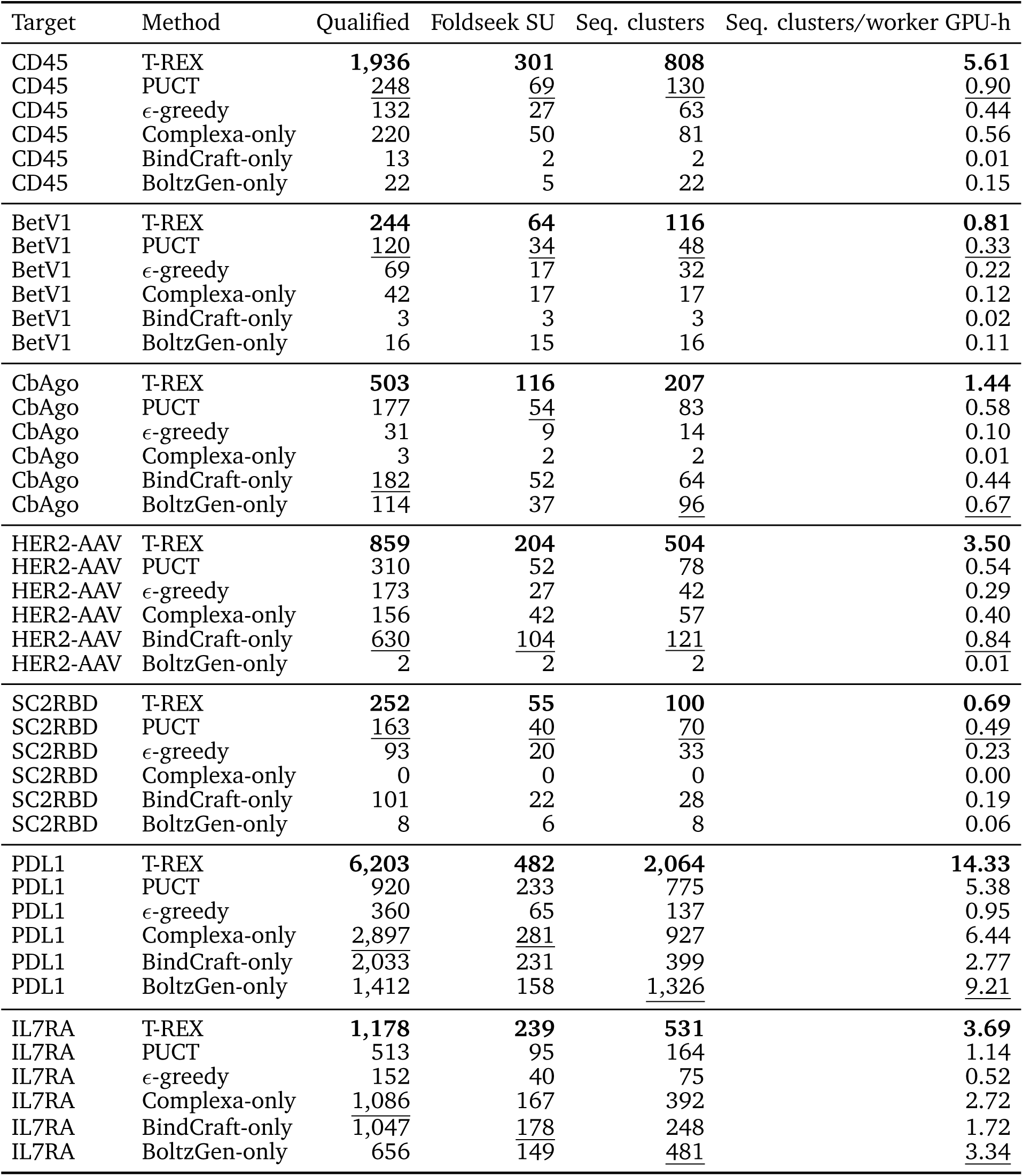
MMseqs2 sequence-cluster throughput among qualified designs at 70% identity. Target set: CD45, BetV1, CbAgo, HER2-AAV, SC2RBD, PDL1 and IL7RA. Binder-chain sequences from all designs satisfying the common qualification rule were clustered before Foldseek deduplication. Each rate divides the sequence-cluster count by the prespecified denominator of 144 H100 worker GPU-h. Foldseek TM0.6 SU is shown for structural context but does not define the sequence-cluster count. Bold and underline mark the best and second-best values within each target and numeric column.

**Table S14.**
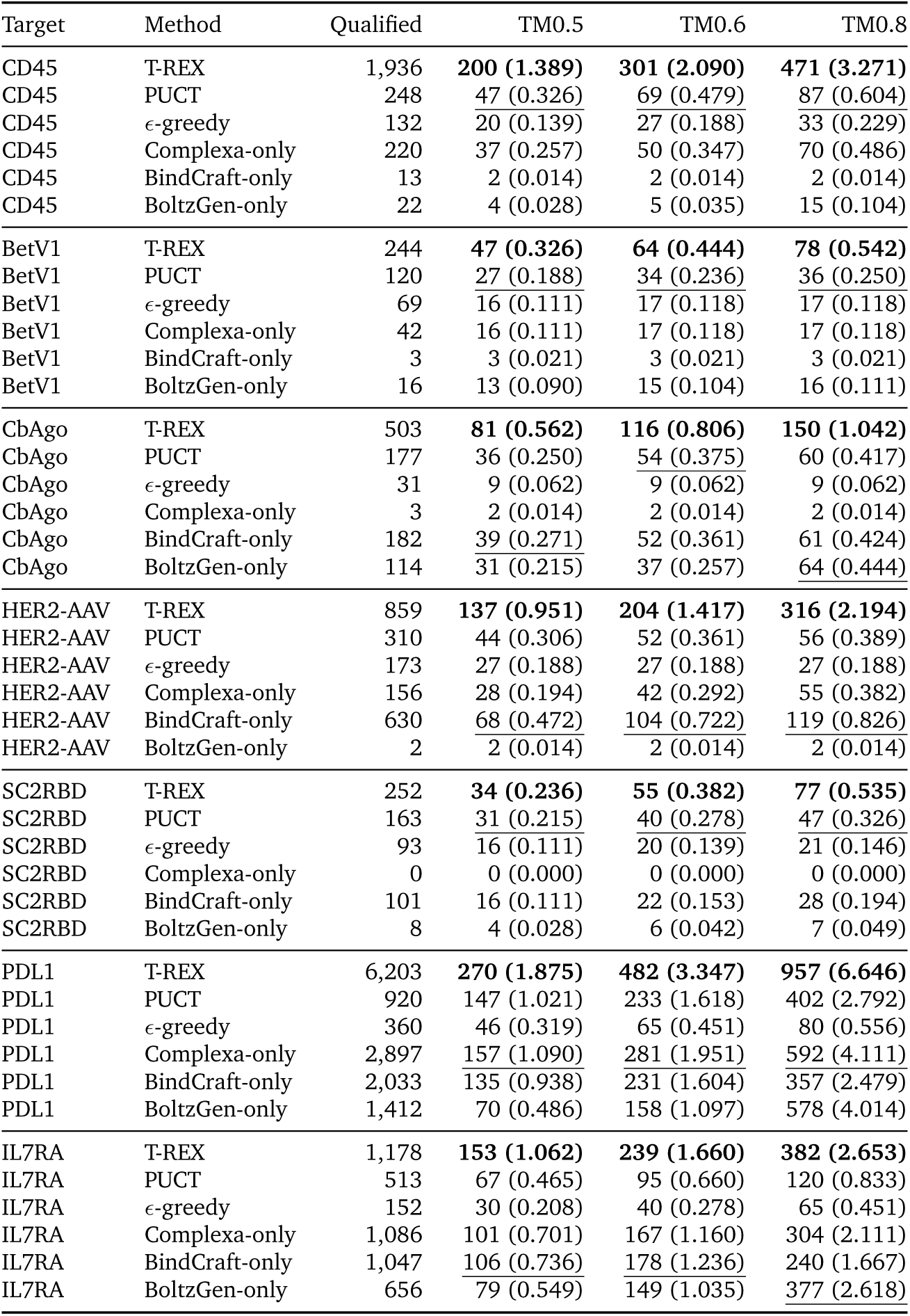
Sensitivity to the Foldseek clustering threshold. Target set: CD45, BetV1, CbAgo, HER2-AAV, SC2RBD, PDL1 and IL7RA. Each rate divides the number of qualified structural clusters by the prespecified denominator of 144 H100 worker GPU-h; Qualified is the number of designs before Foldseek clustering. Bold and underline mark the best and second-best rate within each target and TM threshold; ties share a mark.

The qualification-margin analysis instead holds the final TM0.6 clusters fixed and retains those whose selected representative has *m*_min_ ≥ *τ* (Figure S4; Section B.2). Each representative maximizes *m*_min_ within its cluster, so a retained cluster contains at least one design meeting all three tightened margins. Cluster membership is not recomputed after filtering. At *τ* = 0, the qualified pool recovers the primary count while retaining the original strict scRMSD cutoff.

Across 63 target–margin combinations (*τ* = 0, 0.05, . . ., 0.40), T-REX exceeded PUCT in 50, tied in ten, including seven zero–zero ties, and was lower in three, each by one SU. Thus, the throughput advantage persisted across most tested margin cutoffs, with ties and reversals as fewer clusters remained. This margin-retention analysis is distinct from reclustering at alternative TM thresholds.

**Figure S4.**
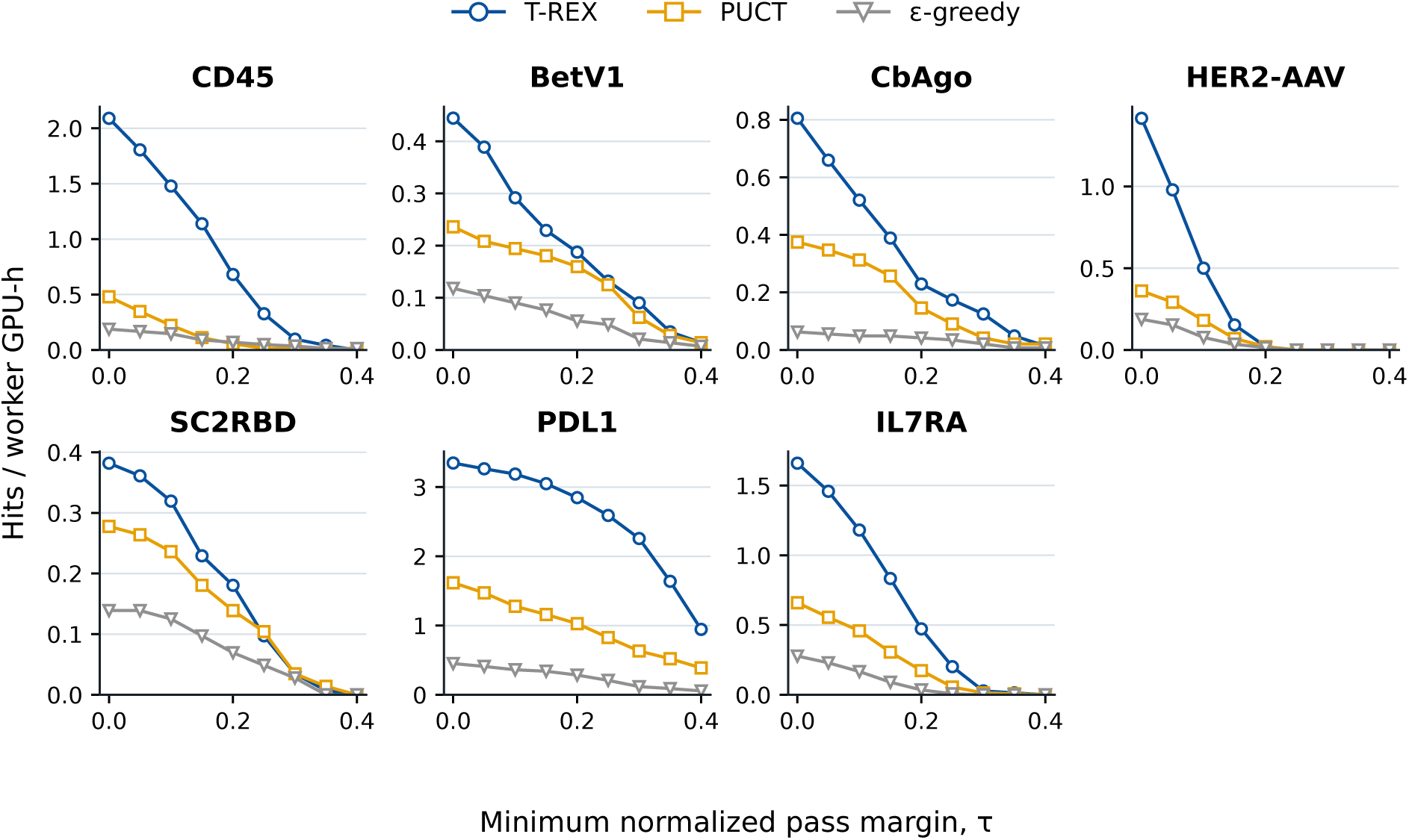
Sensitivity to jointly tightening the three qualification criteria. Fixed final Foldseek TM0.6 clusters per worker GPU-h whose selected representative meets the requirement that the normalized pLDDT, iPAE and scRMSD pass margins are all at least *τ*; *τ* = 0 recovers the primary qualification rule. Rates use the prespecified denominator of 144 H100 worker GPU-h.

#### D.2. Evidence-linked decisions and worked examples

##### D.2.1. Evidence at an archived planning update

This example follows HER2-AAV planning update 6 from archived evidence to the reduced LLM input. Update numbers identify planning steps, not elapsed time. The JSON excerpts here and in Section D.2.2 retain field names, nesting, nulls and booleans; descriptive aliases (example_design, rescue_hypothesis, rescue_candidate) replace archive identifiers. Selected measurements are rounded to four decimals; omitted fields do not imply missing measurements.

**Archived EvidenceSummary excerpt.**

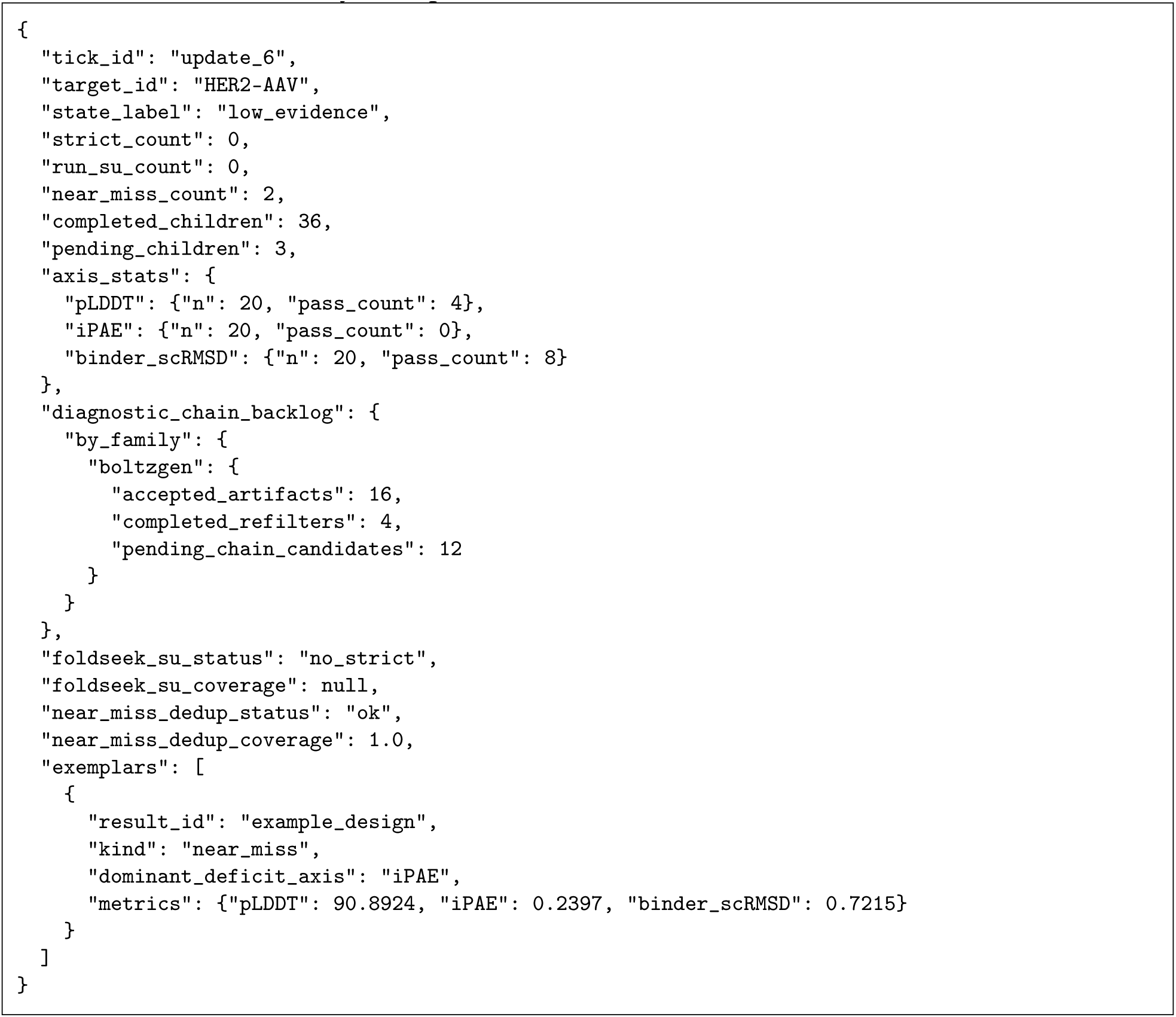

**Qualification and pending evaluation.** The campaign had no qualified designs or online SUs, two near-miss clusters and the deterministically assigned campaign state low_evidence (Section C.4.5). The illustrated near miss passed pLDDT (90.8924) and scRMSD (0.7215 Å) but its normalized iPAE (0.2397) exceeded the cutoff. The LLM later incorrectly described scRMSD as failing; qualification used the recorded measurements (Section D.2.2). Qualified-pool clustering had no structures (no_strict, null coverage), whereas near-miss clustering was complete (ok, coverage 1.0). Of sixteen BoltzGen designs, four had completed standardized AF2 evaluation and twelve awaited evaluation under the admission rules in Section C.8.1; their qualification outcomes were still unknown.

**Criterion summaries and structural duplication.** Each criterion had twenty measurements. For pLDDT, normalized iPAE and binder scRMSD, respectively, pass/near-pass/fail counts were 4/8/8, 0/5/15 and 8/2/10; medians were 85.7853, 0.3446 and 2.2067 Å. These criterion-specific categories follow Section C.4.1. Paired pLDDT–iPAE counts were zero both-pass, eight both-fail, zero pass/fail or fail/pass, and one both-near-pass. Mixed near-pass combinations are omitted from these five patterns, so their counts need not sum to twenty. Among recent cluster-assigned records, *f*_dup_ = 0.75 (Eq. (S3)) and the largest cluster contained 20% of records. This structural duplication measure includes unqualified designs and does not confer SU credit.

**Recorded results and compute.** The 36 *ResultRecords* comprised sixteen Complexa Beam outputs, sixteen BoltzGen outputs and four standardized AF2 evaluations; all had ok status. The family-level attempts and completions therefore counted records, not job launches (Section C.3). Their recorded costs were 0.3216, 0.1362 and 0.0363 worker GPU-h, respectively, totaling 0.4941 worker GPU-h. The three pending_children were running jobs. Elapsed worker-slot time was 1.0003 GPU-h, whereas the stall-time signal was 0.9673 worker GPU-h and six updates had passed without an SU. This signal combines recorded cost and running-worker time, with a zero cost reference before the first SU (Section C.4.2); it differs from elapsed campaign time.

Complexa Beam contributed both near-miss clusters. All three qualification measurements were available for Complexa Beam and AF2-evaluated outputs, but absent from native BoltzGen outputs. A Complexa Beam recipes entry grouped eight descendants under a near-miss outcome, with beam width 4, four branches and four samples; the illustrated design came from the initial Complexa Beam job. Auxiliary and alternative-model measurements provided context without affecting qualification or SU credit (Section C.4.3).

**Four outcome types within one campaign.** These records illustrate the distinctions in Figure 1b: a later Complexa FK steering job timed out without outputs or measurements; twelve BoltzGen designs awaited evaluation; and the illustrated Complexa Beam design missed the iPAE cutoff. Between updates 11 and 12, qualified designs increased from 42 to 43 while online SUs remained at four, with complete, successful qualified-pool clustering. This last case added qualification without increasing the online SU count.

**Reduced evidence input to the LLMs.** The following excerpt was reconstructed using the input-reduction rules in Section C.5.2; it is not a saved historical prompt. The target identifier is omitted, the 36 parent identifiers become a count and availability flag, and cumulative objective fields are grouped in objective_summary. Parent identifiers remained available for candidate construction. HypothesisCards and, for the Supervisor, candidate jobs were supplied separately. The excerpt shows selected evidence fields rather than either role’s complete input.

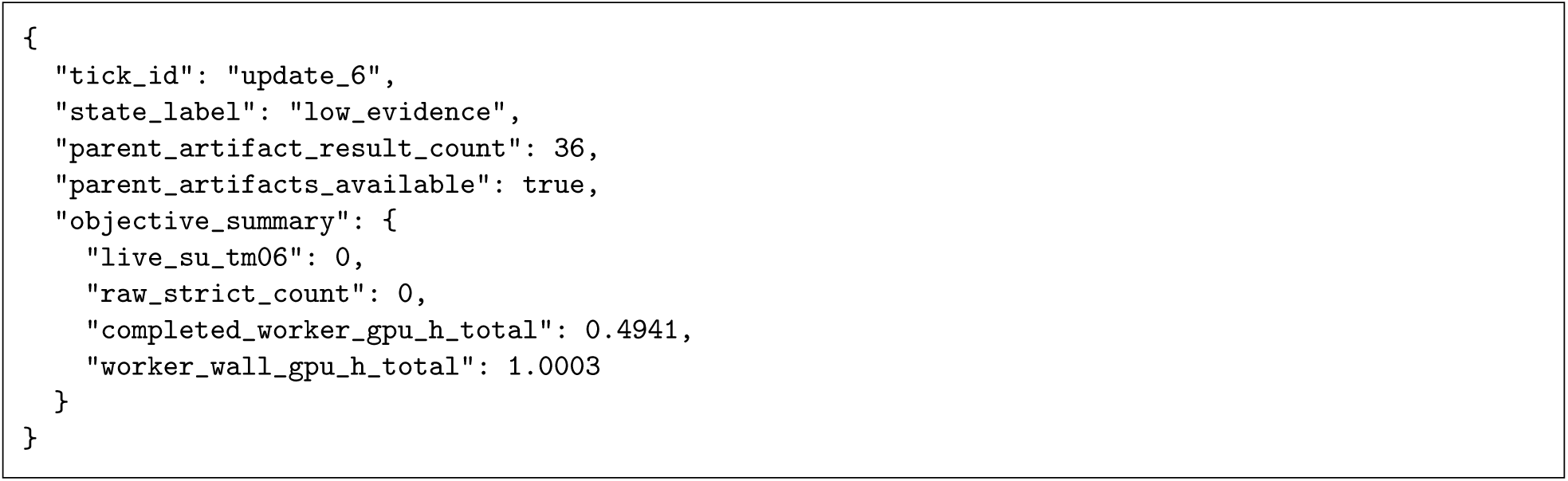

##### D.2.2. From proposal to execution and feedback

The trace continues from update 6 in Section D.2.1, following one proposal through selection, execution and feedback. JSON excerpts show archived HypothesisCard and ActionCandidate fields, using the same descriptive aliases; they are not raw LLM responses.

1. **Planner proposal.** The card proposed a **Rescue** test: decrease normalized iPAE and binder scRMSD while preserving pLDDT. Two other cards proposed BindCraft and Complexa best-of-*n* exploration. The excerpt shows the cited evidence and suggested settings; Section C.5.2 defines the Planner output.

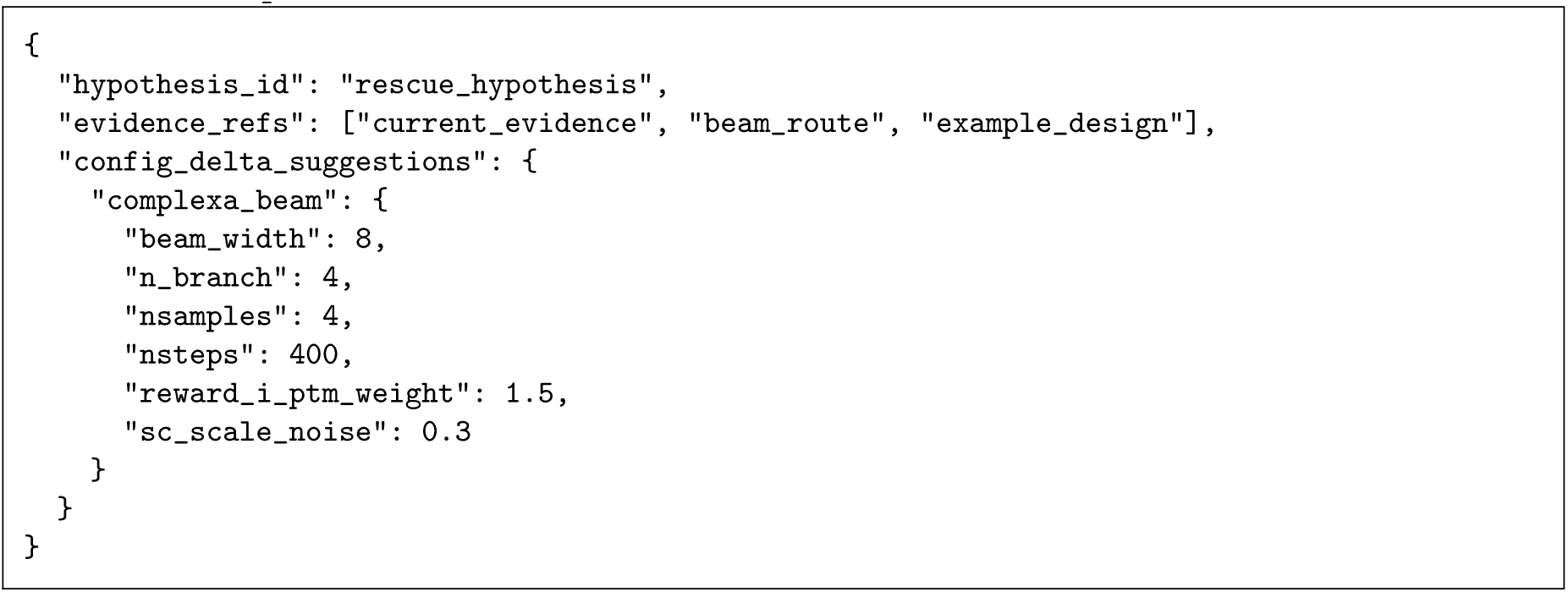
2. **Candidate construction and validation.** Check + Build removed reward_i_ptm_weight: the largest observation count among interface measurements was 20, below the required 32 (Section C.6.2). The remaining settings passed family-availability, configuration, workload and resource checks. The cited design served as the comparison baseline; it was not supplied as a parent structure to this generation job.

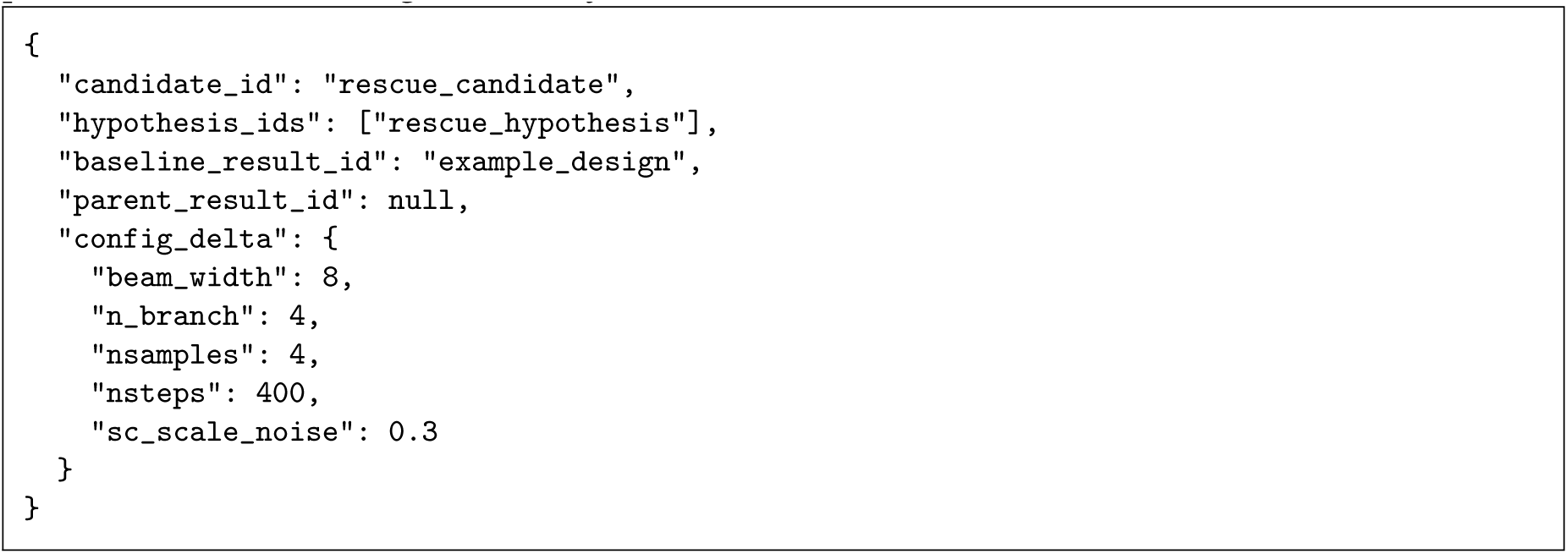
3. **Supervisor prioritization.** The Supervisor ranked Complexa Beam **Rescue** first, BindCraft **Explore** second and Complexa best-of-*n* **Explore** third. Its confidence of 0.50 activated the conditional allocation bounds, but the proposed **Rescue**/**Explore**/**eXploit** shares remained 0.40/0.60/0.00 after adjustment; no fallback was used (Section C.7.1). These are desired shares of subsequent starts, not GPU-time fractions.
4. **Queue admission.** One queue position was available for selection, although all three workers were running jobs. The selector followed the Supervisor’s current ranking across priorities and admitted the first-ranked job; the other two were not selected (Section C.7.2). The recorded launched status denotes queue admission, with execution confirmed separately.
5. **Independent evaluation and confirmed starts.** At the same update, the evaluation-admission rules admitted a previously generated BindCraft design for standardized AF2 evaluation (Section C.8.1). DispatchRecords confirmed that this evaluation and the selected Complexa Beam job passed start-time checks and started. At update 7, the campaign recorded its first qualified design and online SU, from the BindCraft evaluation. The campaign state was productive_duplicate, and recorded cost totaled 1.1468 worker GPU-h. This first SU did not come from the selected Complexa job. The BindCraft route had one standardized AF2 evaluation, cost 0.3491 route GPU-h, and general/recent-productivity statuses healthy/productive (Section C.4.4). With one qualified cluster, its quality-score median and lower quartile both equaled 0.3670; its auxiliary diagnostic-improvement score was 1.0 (Section C.4.3).
6. **Recorded outputs.** Links to the producing candidate job identified 32 ResultRecords with ok status, of which eight met all three qualification criteria. Their update identifier refers to the producing job’s start, not result arrival (Section C.3). These are result counts, distinct from jobs, SUs and hypothesis-support points.
7. **Hypothesis feedback.** At update 9, the card had 32 evaluated descendants, zero support points and zero contradiction points. Qualification did not establish the predicted changes relative to the comparison baseline (Section C.9). Because baseline scRMSD already passed and the card specified no absolute improvement threshold, support required the default 1 Å decrease from 0.7215 Å—an unattainable change. The already-passing criterion also prevented contradiction points under the rule requiring contradiction of every prediction. The card remained active and was retired at update 16, after its ten-update lifetime.

##### D.2.3. Archived decision traces and use of earlier results

The three cases in Figure 3 were selected retrospectively to illustrate follow-up tests prompted by an unsuccessful test, repeated near misses or a parent-specific failure. For every case described here, we linked the proposal, candidate job and launch decision to a confirmed job start and subsequent results. Qualification counts and displayed pLDDT, iPAE and scRMSD values came from standardized AF2 records; online TM0.6 SU gains came from the linked campaign summaries. The HER2-AAV and CbAgo panels compare route-level evidence and output (Figure 3a,b), whereas PDL1 follows a direct ProteinMPNN parent–descendant line (Figure 3c). Batch membership and route-level SU attribution were assessed separately: a job could yield multiple designs, and a route summary could include multiple jobs.

For visualization, output complexes were rigidly aligned on the target to the corresponding evidence or parent complex. The illustrated PDL1 descendant differed from its parent at 35 of 58 residues and had a target-aligned binder C*α* RMSD of 0.714 Å relative to its parent. Qualification scRMSD uses the same target-based alignment but compares a generated complex with its AF2 prediction, rather than a parent with its redesigned descendant.

**Within-campaign evidence reuse across seven targets.** These retrospectively selected cases include unsuccessful reuse and do not estimate success rates. Counts distinguish qualified designs from final Foldseek TM0.6 representatives directly linked to the job; online configuration-level outcomes are identified separately (Section C.4.2). Rates and duplicate fractions describe the online evidence available at the time of the proposal (Section C.4). Card support and refutation assess numerical predictions, not causal explanations; retirement ends a card’s lifetime without either finding (Section C.9).

**CD45: continuing a productive route and reusing it for Rescue.** A Complexa route with adjusted generator-internal reward weights had a recent rate of 2.32 SU/GPU-h, whereas the default route had no recent SU and a duplicate fraction of 0.81. The controller continued the productive Complexa Beam route with its recorded refinement and interface-related weights. A later Rescue proposal cited the same route and weight pair for an interface-limited near miss, testing whether that configuration remained useful for a related case. The continued-production job yielded 27 qualified designs and three final representatives; the later Rescue job yielded eleven qualified designs and one final representative.

**SC2RBD: increasing diversity within a productive generator.** The BindCraft route requesting eight native accepted designs had yielded 1.40 recent SU/GPU-h, but all recently clustered designs occupied one structural cluster. A route requesting sixteen native accepted designs subsequently yielded four SUs across four clusters. The proposed test increased the requested native accepted-design count from eight to sixteen while retaining the interface-weight configuration, then reused that configuration after the narrower route became redundant again. The initial job yielded nineteen qualified descendants and three final representatives; the later repeat yielded ten qualified descendants and one final representative.

**PDL1: testing an intermediate sampling setting.** A backbone sampling-noise value of 0.30 had the highest recent rate, 6.36 SU/GPU-h, but a duplicate fraction of 0.84. A value of 0.20 had lower yield and greater structural duplication, whereas the tested 0.40 configuration had reduced productivity. The controller tested 0.35, within the permitted bounds, while retaining the other route settings to examine a possible trade-off between production and diversification. The job yielded thirty qualified designs and five final representatives.

**HER2-AAV: revising a hypothesis after an unsuccessful test.** Complexa’s native metrics for FK steering indicated strong interface confidence (median ipTM 0.77) but lower fold confidence (median pLDDT 83.4). An earlier pLDDT reward-weight test yielded no qualified designs, and its card was refuted. A subsequent proposal tested sequence–backbone compatibility by retaining FK steering and switching to sequence hallucination (Figure 3a). Five designs qualified. The first outcome summary for the same route and configuration reported three online SUs; this count is not attributable to the individual job. Feedback counted all 32 evaluable designs as support under the card-specific improvement and preservation tests, which are distinct from qualification.

**BetV1: revisiting a generator with earlier structural diversity.** Complexa had become duplicate-rich. BindCraft had produced 33 SUs earlier with greater structural diversity, although it had produced none in the recent window. The controller revisited BindCraft requesting eight native accepted designs and using positive helicity, ipTM and pLDDT weights to test whether another generator family could restore novelty. The job yielded seventeen qualified designs and two final representatives. Its HypothesisCard was retired at the lifetime limit.

**BetV1: unsuccessful reuse of an earlier configuration.** An interface-confidence Rescue proposal cited an earlier configuration that had produced qualified designs. The controller reused Complexa Beam with sequence hallucination, the maximum permitted interface-confidence reward weight and backbone sampling noise 0.50 for another near miss. None of sixteen linked designs qualified. The HypothesisCard was retired at the lifetime limit. Thus, reuse of a previously productive configuration did not guarantee qualification.

**CbAgo: repeating the highest-yield active route.** An FK steering route with a negative iPAE reward weight had the highest recent rate among active configurations, 2.72 SU/GPU-h, and had previously yielded a high proportion of qualified designs. The controller relaunched that route with its recorded refinement and interface-related weights to test whether it continued to add structural clusters. This job, distinct from the one in Figure 3b, yielded eight qualified designs and one final representative.

**IL7RA: continuing production despite a high duplicate fraction.** BindCraft had recent throughput of 1.46 SU/GPU-h and a cumulative online count of 57 SUs despite a duplicate fraction of 0.80. The controller continued the route requesting two native accepted designs while other jobs tested alternatives. Both linked designs qualified and supported the card’s numerical predictions.

#### D.3. Controller behavior and intervention

##### D.3.1. LLM use and interface checks

**Model calls and output validation.** The seven primary campaigns recorded 3,125 LLM calls: 1,719 Planner and 1,406 Supervisor calls (Table S15, part A). Seven Supervisor responses failed structured-output validation, representing 0.22% of all model calls. All seven were parsed as JSON, but six cited evidence references outside the allowed set and one used an invalid resource class. These rejected responses triggered deterministic fallback; Sections C.5.2 and C.7.1 describe validation and response handling.

**Selection paths and confirmed starts.** Model-call validation and fallback during selection have different denominators. We counted unique planning updates in *SupervisorDecision* records and linked each confirmed generation or redesign start through its *LaunchDecision* to the selecting update. Of 1,719 updates, 1,398 used the current Supervisor candidate ranking and 321 (18.7%) used fallback. These comprised 313 updates without hypotheses or candidates for a Supervisor call, seven with rejected Supervisor responses and one carrying a fallback reason from an empty Planner-card list despite a valid Supervisor response. The counts include updates that started no generation or redesign job.

All 1,454 generation or redesign starts linked to a recorded selection: 1,446 to updates using the Supervisor ranking and eight (0.55%) to fallback. The 3,160 separately admitted AF2 starts were excluded from this denominator. The selection path does not establish that the LLM proposed a job: fixed construction rules can add candidates to either path (Sections C.6 and C.7).

**Controlled interface checks.** The Planner tests used five synthetic evidence situations: productive work, concentrated near misses, stall, ambiguous evidence and qualified but structurally redundant output. Supervisor tests used the same situations except ambiguous evidence, with five calls per situation for each role. Checks required valid parsed outputs; normalized Supervisor allocation proportions also had to satisfy case-specific expectations: productive, *X* ≥ 0.45 and *E* ≤ 0.30; near-miss-enriched, *R* ≥ 0.40; stalled, *E* ≥ 0.25 and *X* ≤ 0.45; structurally redundant, *E* ≥ 0.25. All output-validity and allocation checks passed (Table S15, part B).

Repeatability and sampling-sensitivity tests used one fixed productive-evidence case with five candidate jobs: two **eXploit**, two **Rescue** and one **Explore**. Each test made ten Supervisor calls, at temperature 0 for repeatability and at temperatures 0.00–0.45 in 0.05 increments for sampling sensitivity. Within each test, all pairs of valid responses were compared using within-priority first-ranked-candidate agreement, Kendall correlation and the sum of absolute allocation-share differences, as defined in Table S15, part B. Three sampling-sweep responses omitted one Rescue candidate, so the reported agreement does not establish identical full rankings. These checks assess output validity and repeatability for the tested inputs, without validating explanations or establishing biological accuracy, campaign-wide reliability or a causal benefit of LLM reasoning.

**Table S15.**
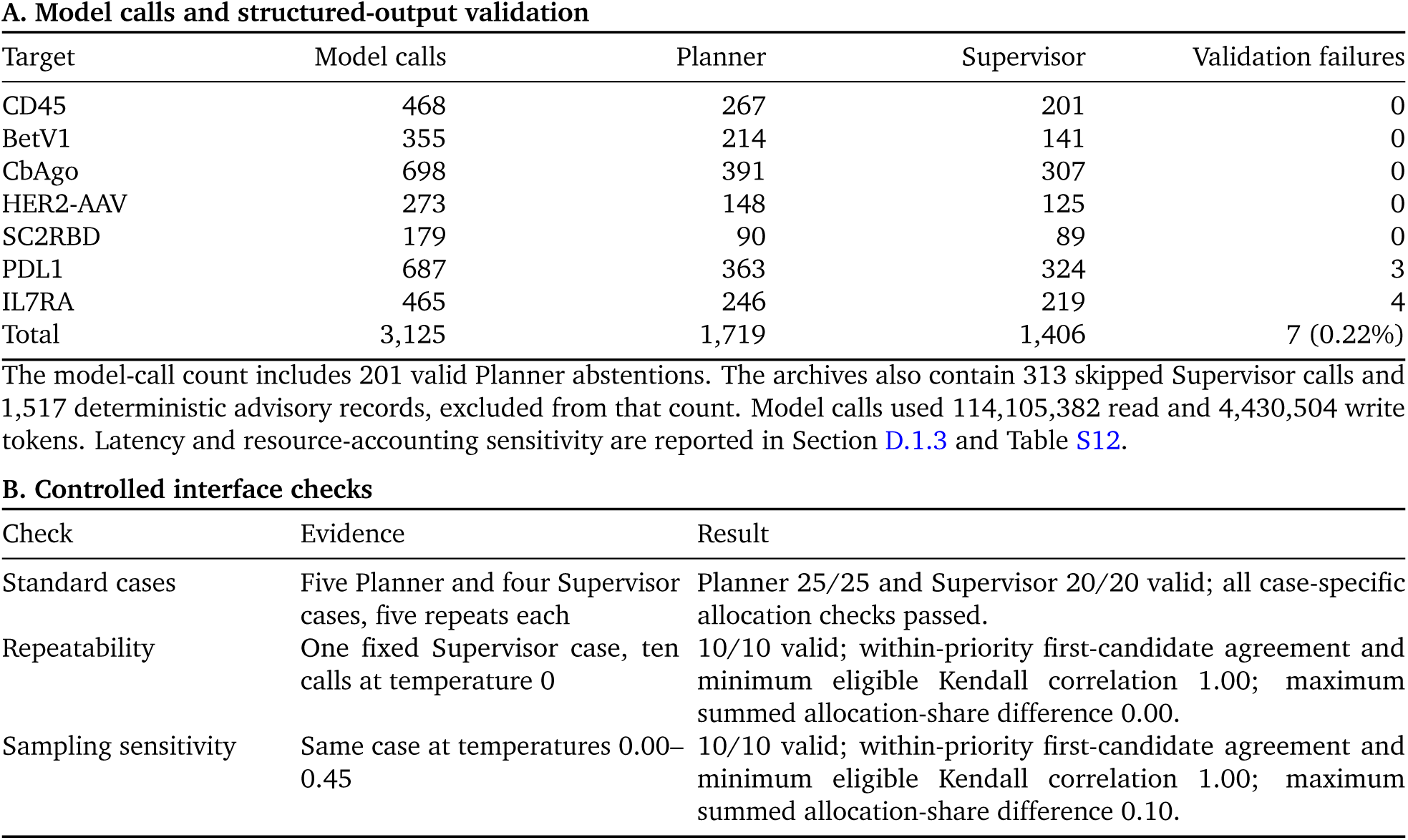
LLM calls and controlled interface checks. Part A counts model calls, including valid Planner abstentions, but excludes skipped calls and deterministic advisory records. Part B uses synthetic inputs. Agreement is the fraction of pairwise comparisons within priorities with the same first-ranked candidate. Kendall correlation requires identical candidate sets with at least two jobs. Allocation difference is the sum of absolute differences across the R/E/X shares. Fixed handling of invalid or unavailable jobs is described in Sections C.5.2, C.6 and C.8.2.

##### D.3.2. Campaign-wide allocation and realized action-space use

Figure 4a groups 1,454 confirmed generation and redesign starts by campaign-state label and recorded R/E/X priority. Within-state percentages use all generation and redesign starts in that state as the denominator; required AF2 evaluations are excluded. The main-text percentages correspond to 48/65 **Explore** starts in low evidence, 41/80 **Rescue** starts across stall and deep stall combined, and 930/1,133 **eXploit** starts in productive-with-duplication states. No generation or redesign start was recorded in structural duplicate collapse. The state rules are defined in Supplementary Section C.4.5; panels c,d instead use the retrospective annotations described below.

**Evidence-focus annotation.** Each start was linked to structured decision text by target and candidate-job identifier, retaining repeated starts as separate events. The four annotation categories were interface confidence, fold confidence, backbone consistency and structural redundancy. For linked decisions, manual annotations covering 27 planning decisions (35 starts) took precedence over text matching. Otherwise, case-insensitive matching first examined the proposed-job description (what). Only when this field matched no category did matching examine the combined evidence and rationale fields (observed_signal, inference, why and action_implication). One matched category assigned a single focus; matches to multiple categories assigned a multi-focus label. Starts with no match or no linked decision remained unassigned.

Panels c,d include the 1,305 starts assigned a single focus, excluding multi-focus and unassigned starts. The Figure 4 label *backbone consistency* denotes structural self-consistency, including wording about scRMSD, pose error or backbone drift (Section B.1). These annotations describe cited evidence, not independently diagnosed design limitations; annotation was not blinded and inter-rater agreement was not assessed.

Supplementary Table S16. reports measured qualification failures and structural redundancy separately from these annotations of decision text. Associations among campaign state, evidence focus and priority are descriptive: deterministic state rules, candidate construction and allocation bounds also influence realized starts.

**Table S16.**
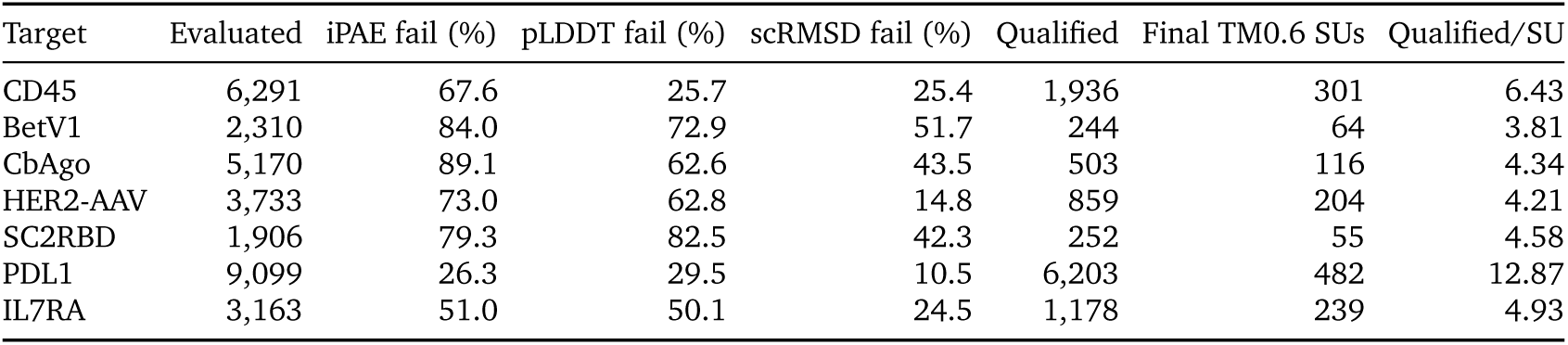
Target-level qualification and structural-redundancy profiles in the seven primary T-REX campaigns. Each percentage is the fraction of standardized AF2-evaluated designs that failed the named criterion; designs can fail more than one criterion. Qualified designs per final TM0.6 SU reports structural repetition after the common qualification rule and final Foldseek clustering.

**Follow-up job types.** For Figure 4d, jobs were classified in the following order. ProteinMPNN jobs were assigned to candidate redesign, explicit repeats of archived routes to recorded-route replay, and the first start of a generation family within each target campaign to first method use. For the remaining jobs, case-insensitive matching used the proposed-job description, or the recorded configuration when that description was absent. A match to sequence[_ -]?hallucination assigned candidate redesign; otherwise, reward, weight, penalty or objective assigned scoring-weight adjustment. All remaining jobs were assigned sampling or search adjustment. This order makes the categories mutually exclusive; the text-based categories describe proposed jobs and do not establish that every matched job changed scoring weights or sampling settings.

**Realized action-space use.** The seven primary T-REX campaigns used all six registered generation families and ProteinMPNN redesign, spanning 161 target-specific configurations and 1,454 confirmed generation or redesign starts (Supplementary Table S17). The table pools direct SU attribution across targets; Supplementary Figure S5 shows the target-level counts. Each final SU is attributed to the family that produced its selected representative, rather than all jobs that contributed to its ancestry. This attribution does not measure causal contribution or cost-normalized efficiency. Permitted configuration ranges are given in Supplementary Table S5.

**Table S17.**
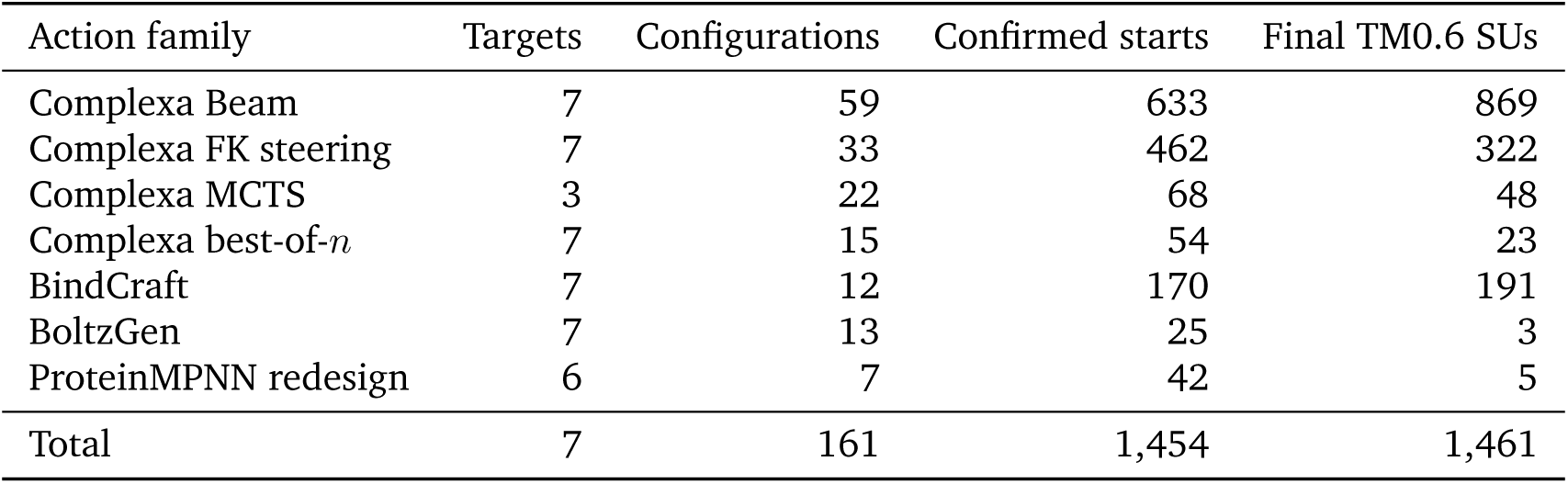
Realized generation and redesign action space in the seven primary T-REX campaigns. Distinct recorded configuration settings are counted separately for each target and action family; the same settings used for two targets therefore count twice. Confirmed starts exclude standardized AF2 evaluation. Final TM0.6 SUs are attributed to the action family that directly produced each representative after evaluation, qualification and clustering. One start can return multiple designs and can therefore receive credit for more than one SU.

**Figure S5.**
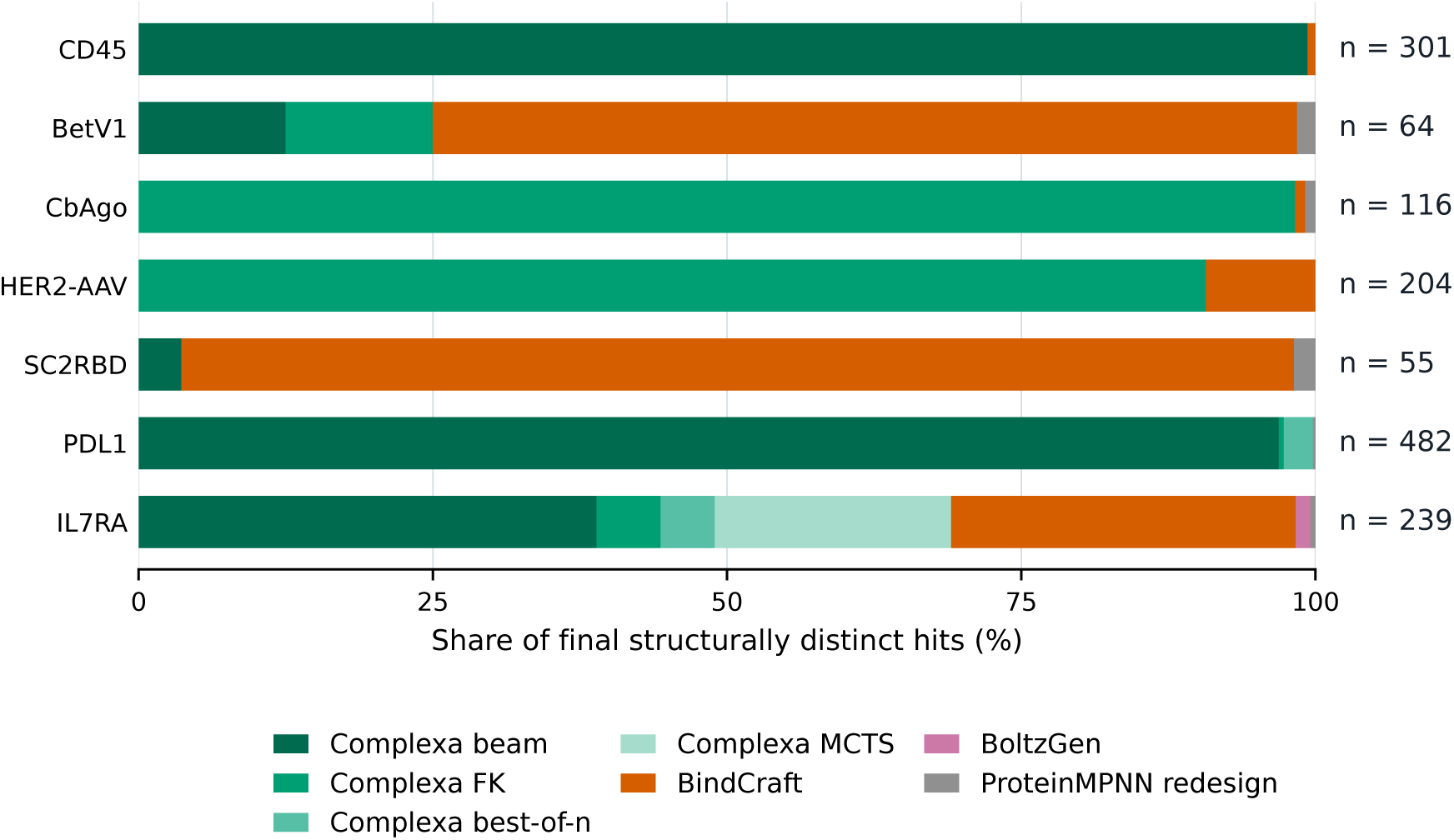
Action-family attribution of final T-REX TM0.6 structure-unique units by target. Each bar partitions the target’s final Foldseek TM0.6 clusters according to the generation or redesign family that directly produced the selected representative; labels at right give the total number of structure-unique units (SUs). Each cluster was counted once, and the standardized AF2 job that evaluated an output received no direct credit. A generation or redesign job could produce several designs and could therefore contribute more than one SU. The plot describes output attribution, not causal contribution or per-start efficiency. Supplementary Table S17 reports the corresponding counts pooled across targets.

##### D.3.3. Intervention without explicit Rescue prioritization

To test explicit **Rescue** prioritization, we ran one intervention campaign per target with a planned budget of 100 worker GPU-h. The action space, structured evidence, resource limits and campaign-state thresholds were retained. Planner and Supervisor priorities were restricted to **Explore** and **eXploit**; fallback allocation and priority accounting also used only these two priorities. Candidates assigned **Rescue** by deterministic rules were relabeled **Explore**. Rescue terminology was removed from LLM instructions, and the near-miss-enriched state was given the neutral label blocker_rich without changing its definition.

**Recovery and time in stall.** Figure 4b uses planning records within the first 100 worker GPU-h of each policy. After retaining the last record at each worker-time value, each transition into stall or deep stall began a new entry, including a transition from stall to deep stall (state definitions in Section C.4.5). Recovery was the first later planning record at which the highest online TM0.6 SU count observed so far exceeded its value at entry, even if the state had changed. Recovery times therefore have planning-record resolution. Entries without recovery by cutoff remained in the recovery-fraction denominator. Recovery-time medians used only recovered entries, without adjusting for unrecovered entries.

Full T-REX recovered in 26 of 27 entries, compared with 14 of 20 under the intervention; one and six entries, respectively, had no observed recovery. These entries are repeated observations within campaigns and can share a recovery observation: the 26 and 14 recovered entries corresponded to 21 and nine distinct target–recovery-time pairs, respectively.

To calculate stall occupancy, each state label was carried to the next planning record, with the last label extended to 100 worker GPU-h. The total duration assigned to stall or deep stall, divided by the summed window durations across targets, gave pooled occupancies of 3.8% for full T-REX and 27.7% for the intervention.

**Endpoint comparison.** Supplementary Table S18 compares final intervention SU counts with full-T-REX counts from the retrospectively reconstructed accumulation curves in Section D.1.2, rather than the online counts used for recovery. Each full-T-REX curve was read at the worker time reached by its corresponding intervention, capped at 100 worker GPU-h. Full T-REX was planned for 144 worker GPU-h, so this is a comparison at the same observed worker time, not between campaigns run with the same planned budget.

**Table S18.**
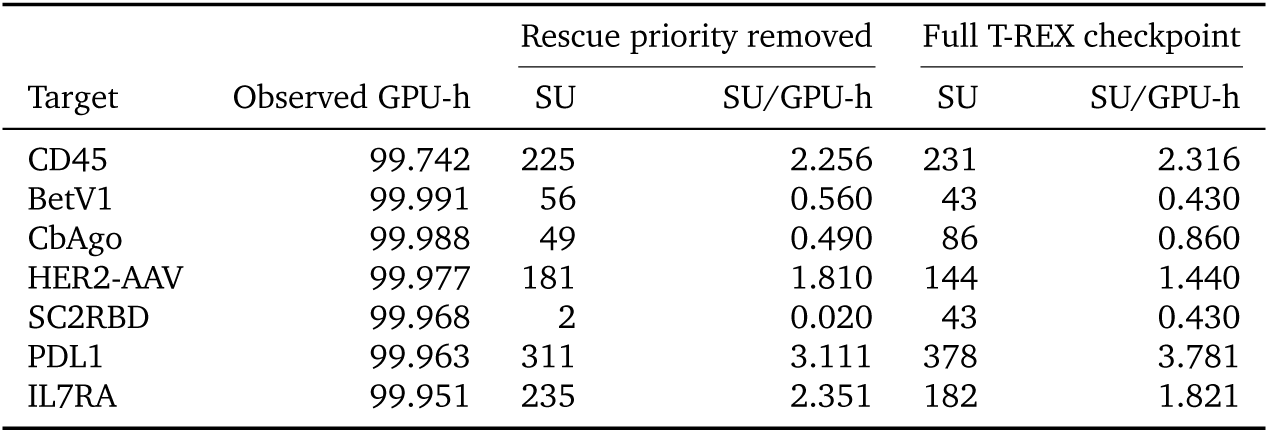
Intervention removing explicit Rescue priority. The intervention was planned for 100 H100 worker GPU-h; its columns report final TM0.6 SU counts and rates. Full-T-REX counts come from the reconstructed accumulation curves in Section D.1.2, read at the corresponding observed worker time, from campaigns planned for 144 H100 worker GPU-h. Both rates divide by the intervention’s observed worker time, using unrounded values. Each target has one campaign per condition; the comparison is descriptive.

Full T-REX had higher counts at this checkpoint for four targets and lower counts for three. The comparison thus does not establish a uniform throughput benefit. Removing **Rescue** could also change how often stalls occurred and which stalls entered the recovery analysis. The intervention jointly changed LLM instructions, priority allocation and state-label wording; it does not isolate an allocation coefficient or the contribution of LLM reasoning.

#### D.4. Post-hoc characterization of binder designs

##### D.4.1. Additional prediction models and metrics

We characterized the fixed representatives defined in Section B.2 using additional model predictions and scores. The set contained 3,766 representatives across six methods and seven targets, with one representative per final Foldseek TM0.6 cluster within each method–target pair. All had finite results for every measurement in Table S19. Harmonized AF2 re-evaluated complexes with common settings, and actifpTM used the same AF2-Multimer configuration (Evans et al., 2021; Varga et al., 2025). Both therefore used the same predictor family as qualification. ESMFold2 and Boltz-2 instead re-predicted complexes from target and binder sequences alone (Candido et al., 2026; Passaro et al., 2025). These post-hoc analyses did not guide allocation or change qualification, clustering or SU counts.

The AF2-derived interaction prediction score from aligned errors (ipSAE) is a higher-is-better interface-confidence score, distinct from the lower-is-better normalized iPAE used for qualification (Dunbrack Jr, 2025). We report the arithmetic mean of its two directional values; actifpTM weights interface confidence by predicted contact probabilities. PRODIGY supplied predicted binding free energy Δ*G* and the dissociation constant *K*_D_ derived from it (Xue et al., 2016). These are predictions of confidence or affinity, rather than experimental binding measurements. Table S19 gives evaluator inputs, settings, units and score aggregation.

Figure 2c shows selected post-hoc thresholds; Supplementary Figure S6a–d reports all 24 thresholds across harmonized AF2, actifpTM, PRODIGY, ESMFold2 and Boltz-2. Each threshold was applied separately to the same fixed representative set, and passing counts were pooled across the seven targets. T-REX retained the largest pooled count at every threshold. These pooled counts reflect both the size of each method’s qualified representative pool and the fraction passing each threshold. The thresholds summarize related measurements, including Δ*G* and its derived *K*_D_, and are not independent validation experiments.

**Figure S6.**
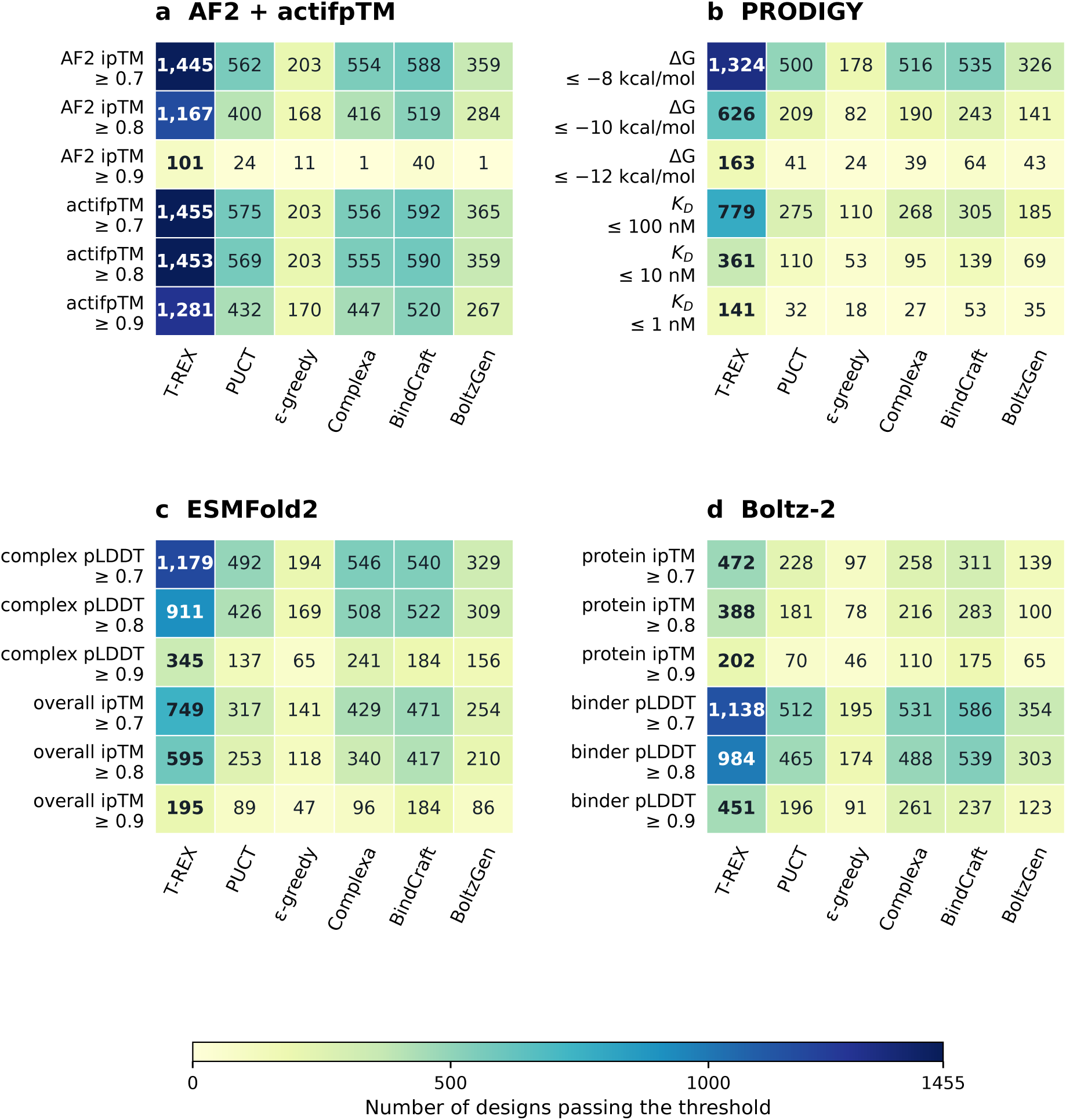
Fixed cluster representatives retained across post-hoc evaluation thresholds. Counts of fixed representatives from final Foldseek TM0.6 clusters passing thresholds for **a**, harmonized AF2 ipTM and actifpTM; **b**, PRODIGY predicted Δ*G* (kcal mol*^−^*^1^) and derived *K*_D_ (nM); **c**, ESMFold2 complex mean pLDDT and package-reported overall ipTM; and **d**, Boltz-2 protein ipTM and binder pLDDT. ESMFold2 and Boltz-2 pLDDT thresholds use the reported 0–1 scale. Counts are pooled across seven targets. Each threshold was applied separately to the same representative set. All four panels share one count scale; bold values mark the largest count at each threshold. Complexa, BindCraft and BoltzGen denote generator-only baselines. These post-hoc diagnostics did not affect allocation, qualification or SU counting.

**Table S19.**
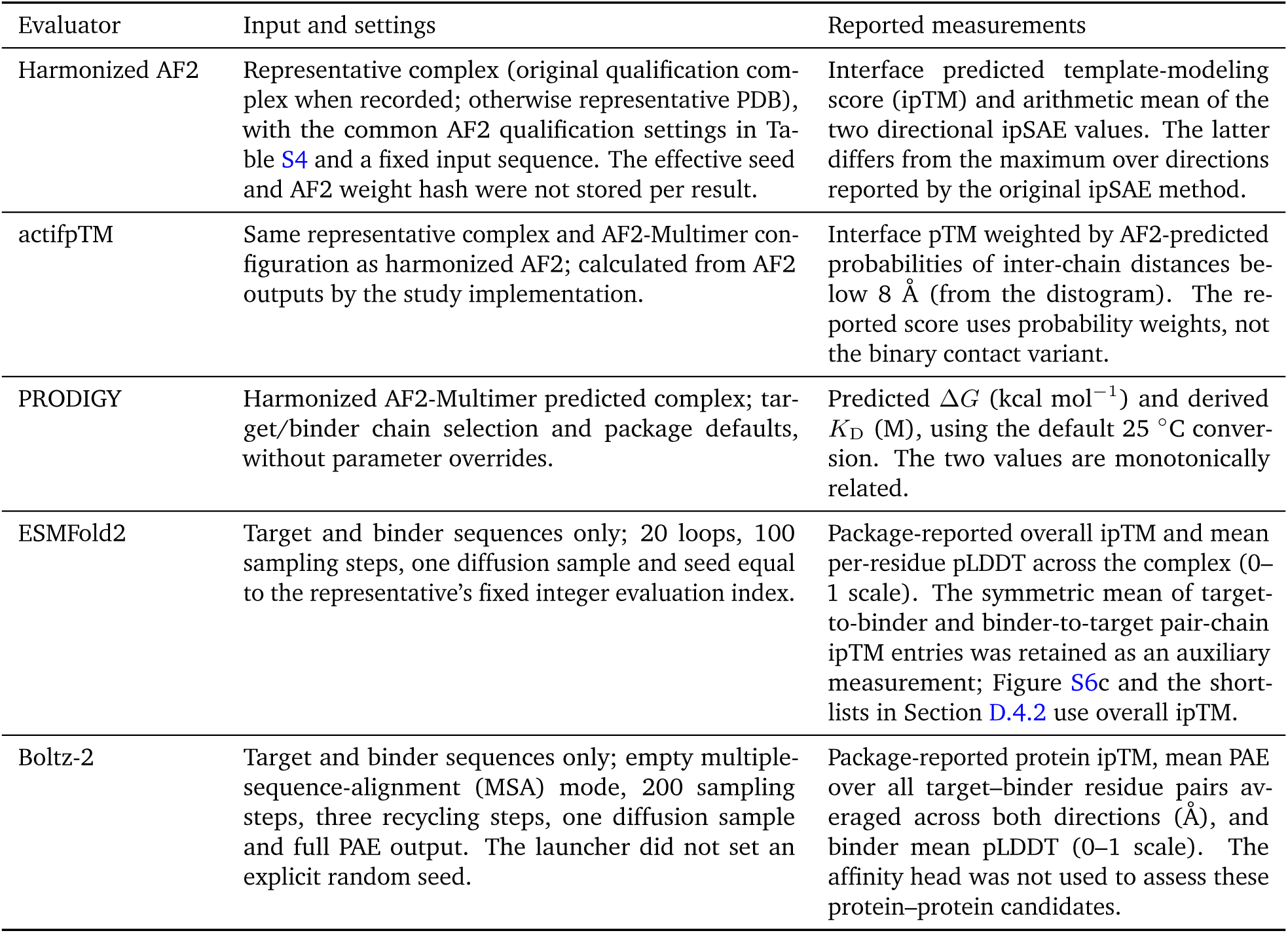
Post-hoc evaluator inputs, settings and outputs. All 3,766 fixed representatives across six methods and seven targets had finite results for every listed measurement. Software versions and model revisions are listed in Table S4.

##### D.4.2. Metric-specific shortlists

Tables S20–S22 summarize fixed TM0.6 representatives from each method–target pair under six separate rankings: minimum qualification margin, AF2 ipTM, mean directional ipSAE, ESMFold2 overall ipTM, actifpTM and PRODIGY predicted affinity. Each ranking selects its own shortlist; Δ*G* and *K*_D_ describe the same affinity-ranked shortlist. From *N* available representatives, each list retains *n* = min(10, *N*) without padding. Margins are defined in Section B.2, and post-hoc metrics in Section D.4.1. These summaries describe metric-specific selection from pools of different sizes, not one common multi-metric shortlist. Because the ranking scores also summarize the selected designs, this is not independent validation.

**Selection and tie-breaking.** The margin ranking orders designs by decreasing *m*_min_, then mean margin, then pLDDT, iPAE and scRMSD margins, in that order. Each confidence ranking uses decreasing values of its named score, then *m*_min_ and mean margin. The affinity ranking uses increasing *K*_D_, then Δ*G*, followed by decreasing *m*_min_ and mean margin. Remaining ties are resolved by ascending lexicographic order of the stable record identifier. All rankings use values before display rounding.

**Table S20.**
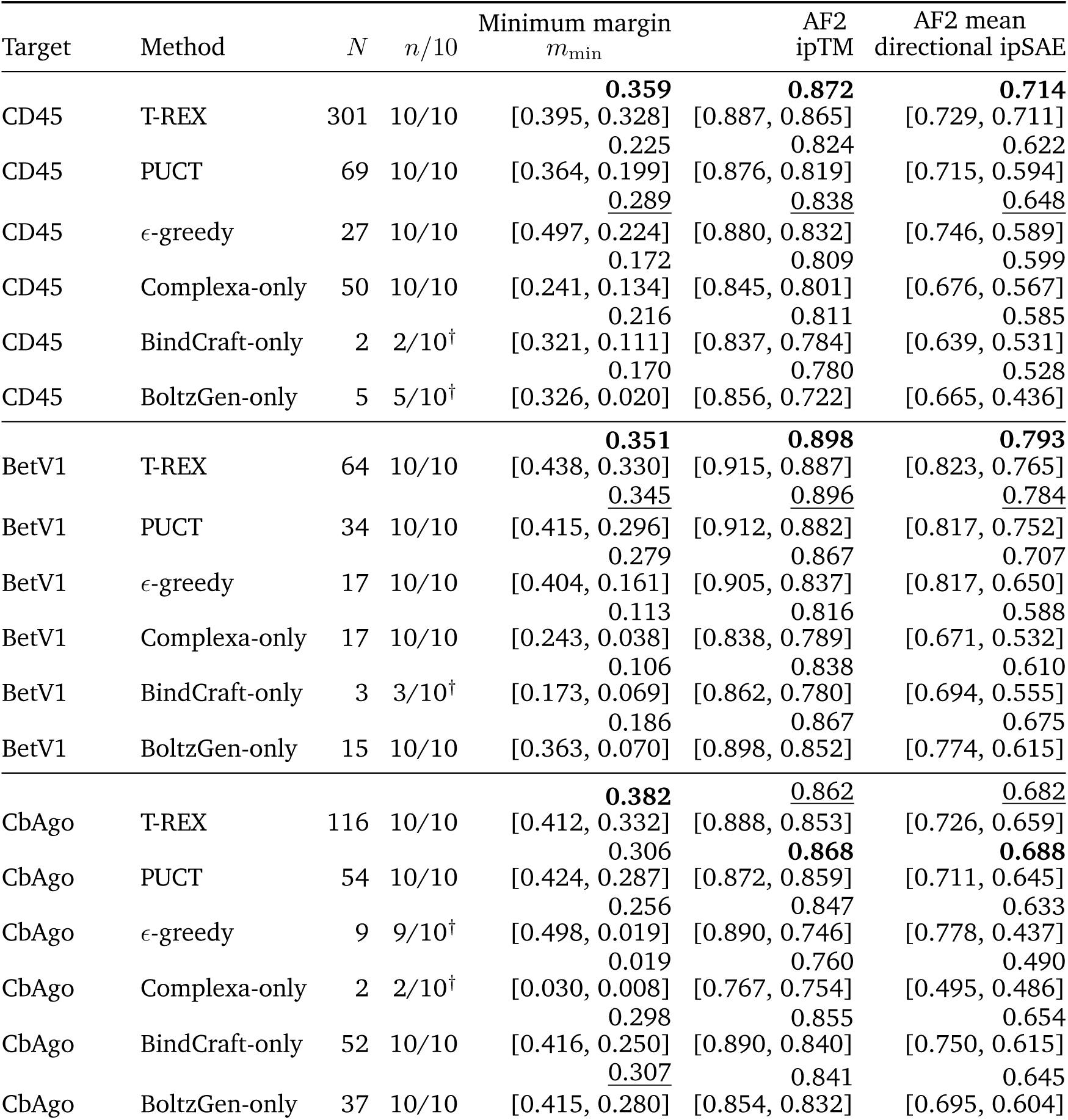
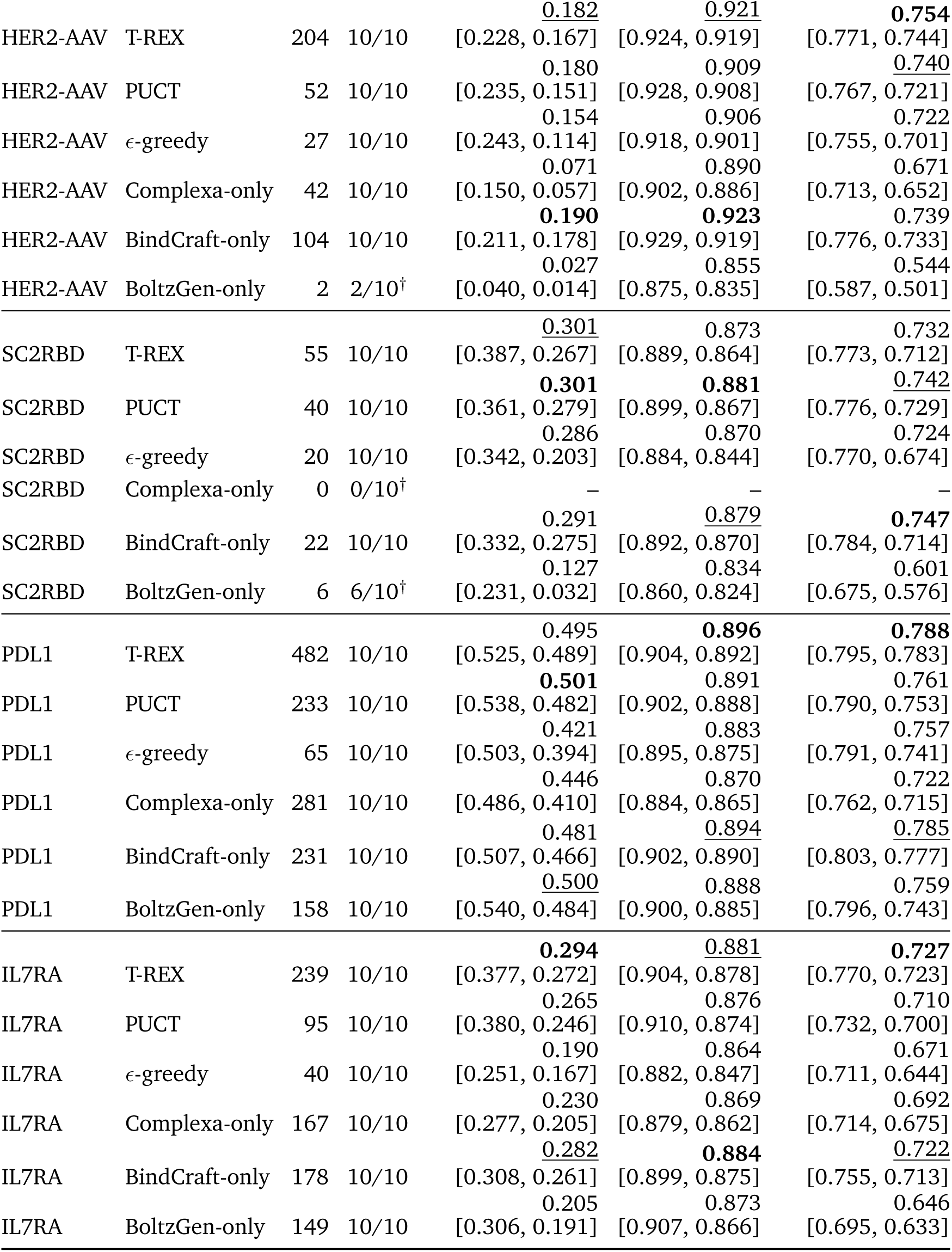
Qualification-margin and AF2-confidence shortlists. Each metric selects its own top *n* = min(10, *N*) representatives from the method–target pool of size *N* and reports median [best, worst]; larger values rank higher. Within each target and metric, bold and underline mark the highest and second-highest distinct unrounded medians among lists of ten designs; ties share a mark. Identical rounded values can carry different marks. Shorter lists carry a dagger and are excluded from highlighting; – denotes an empty pool. Selection rules are given in Section D.4.2.

**Table S21.**
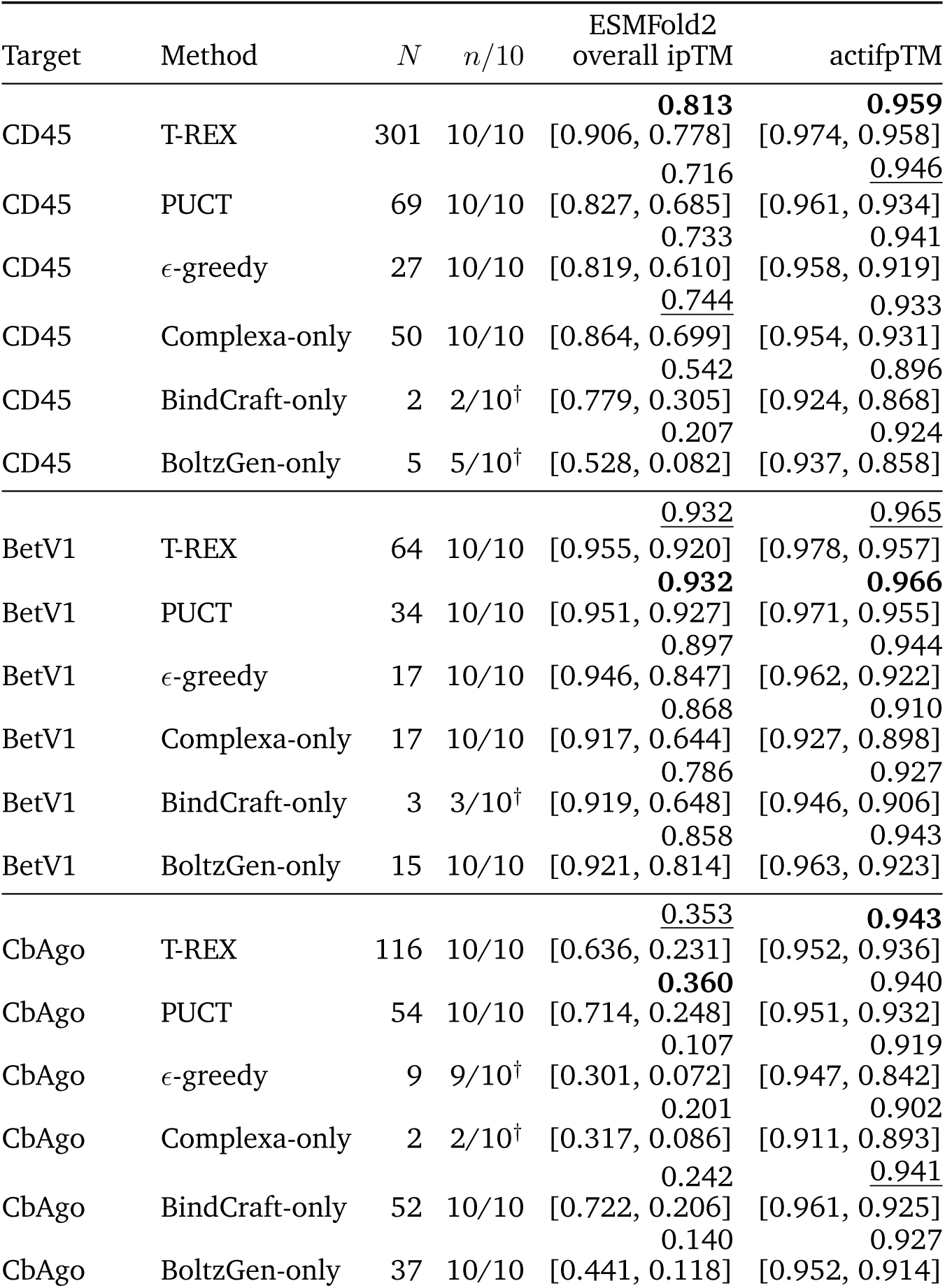
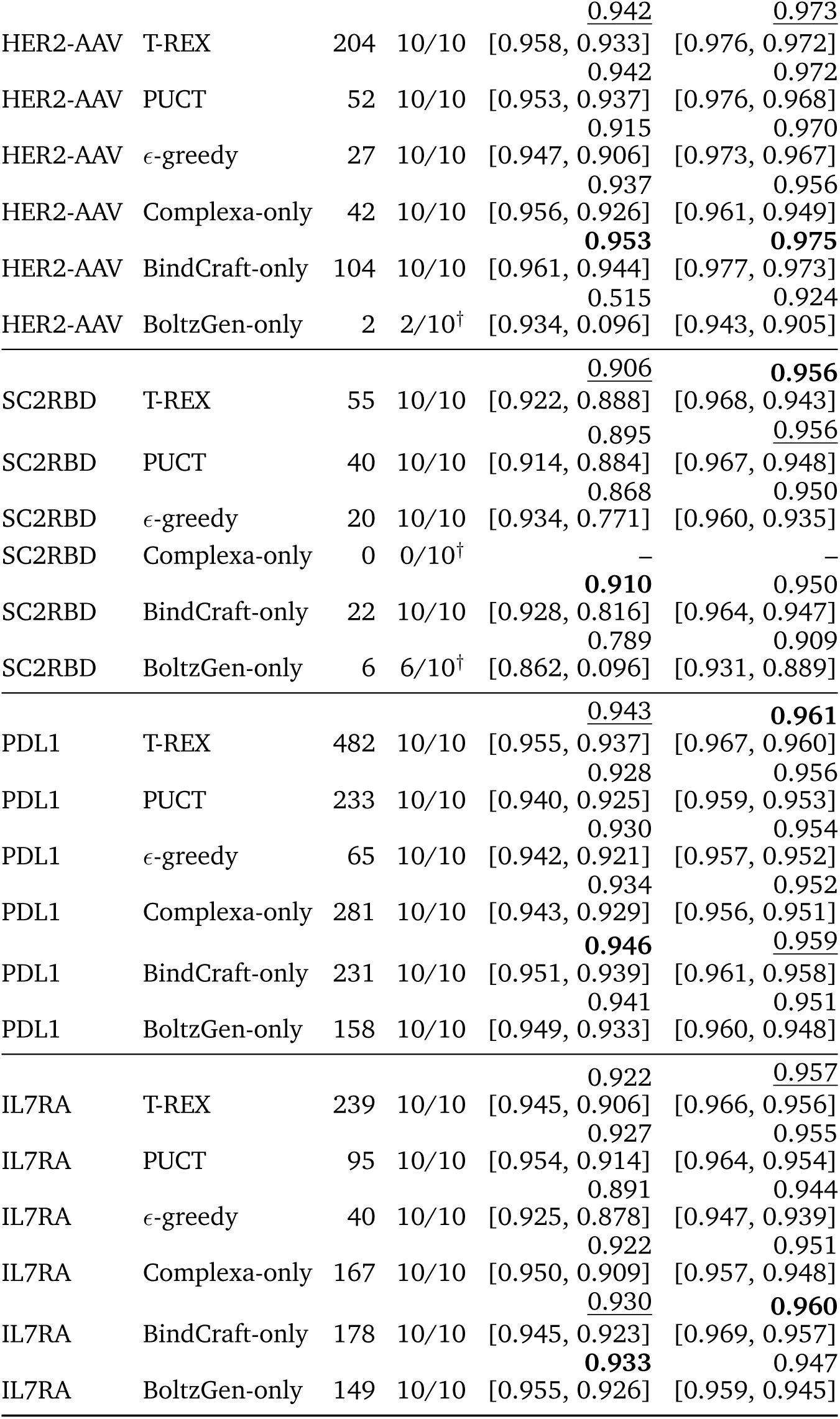
ESMFold2 and actifpTM confidence shortlists. Each metric selects its own top *n* = min(10, *N*) representatives from the method–target pool of size *N* and reports median [best, worst]; larger values rank higher. ESMFold2 uses the package-reported overall ipTM. Within each target and metric, bold and underline mark the highest and second-highest distinct unrounded medians among lists of ten designs; ties share a mark. Identical rounded values can carry different marks. Shorter lists carry a dagger and are excluded from highlighting; – denotes an empty pool. Selection rules are given in Section D.4.2.

**Table S22.**
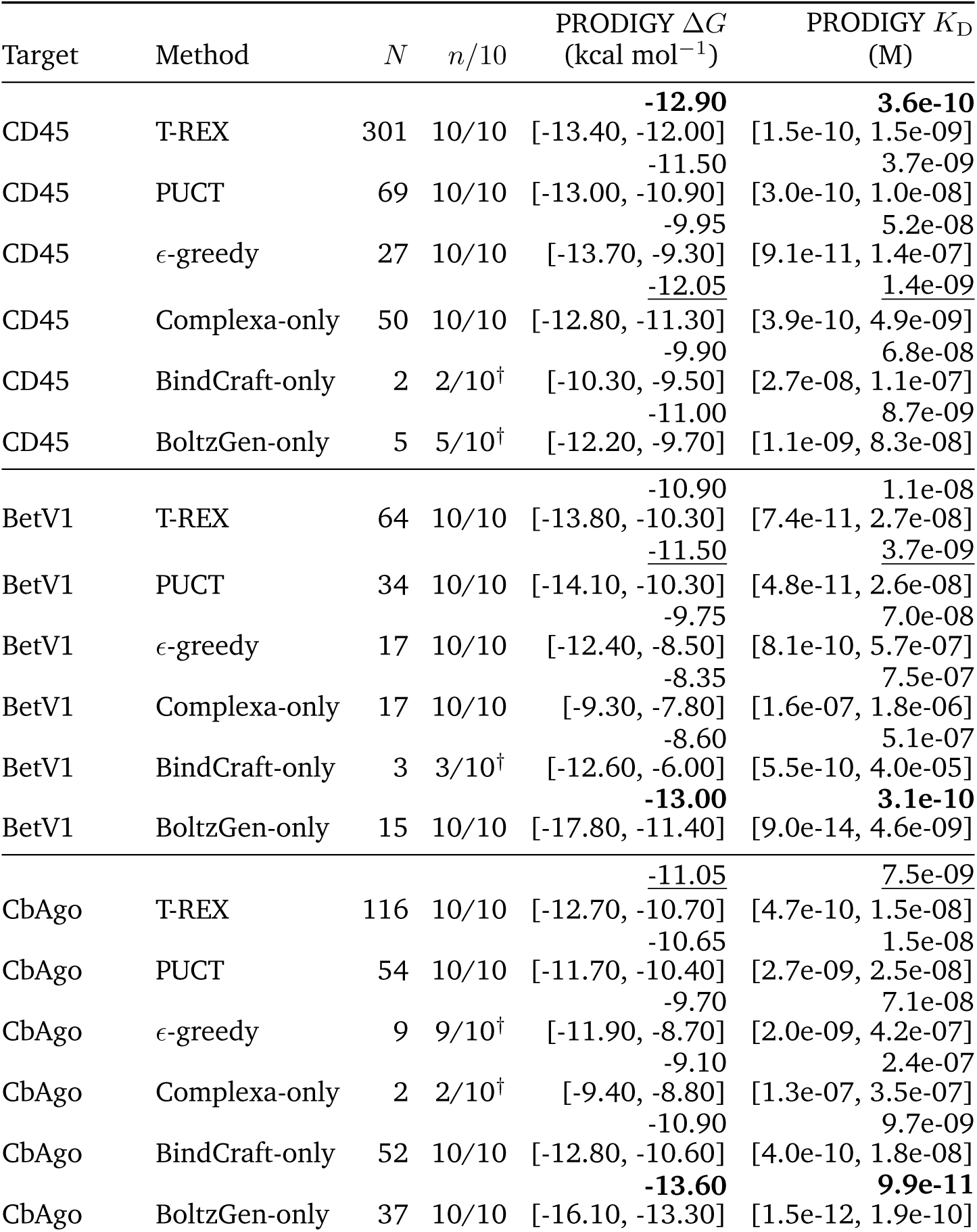
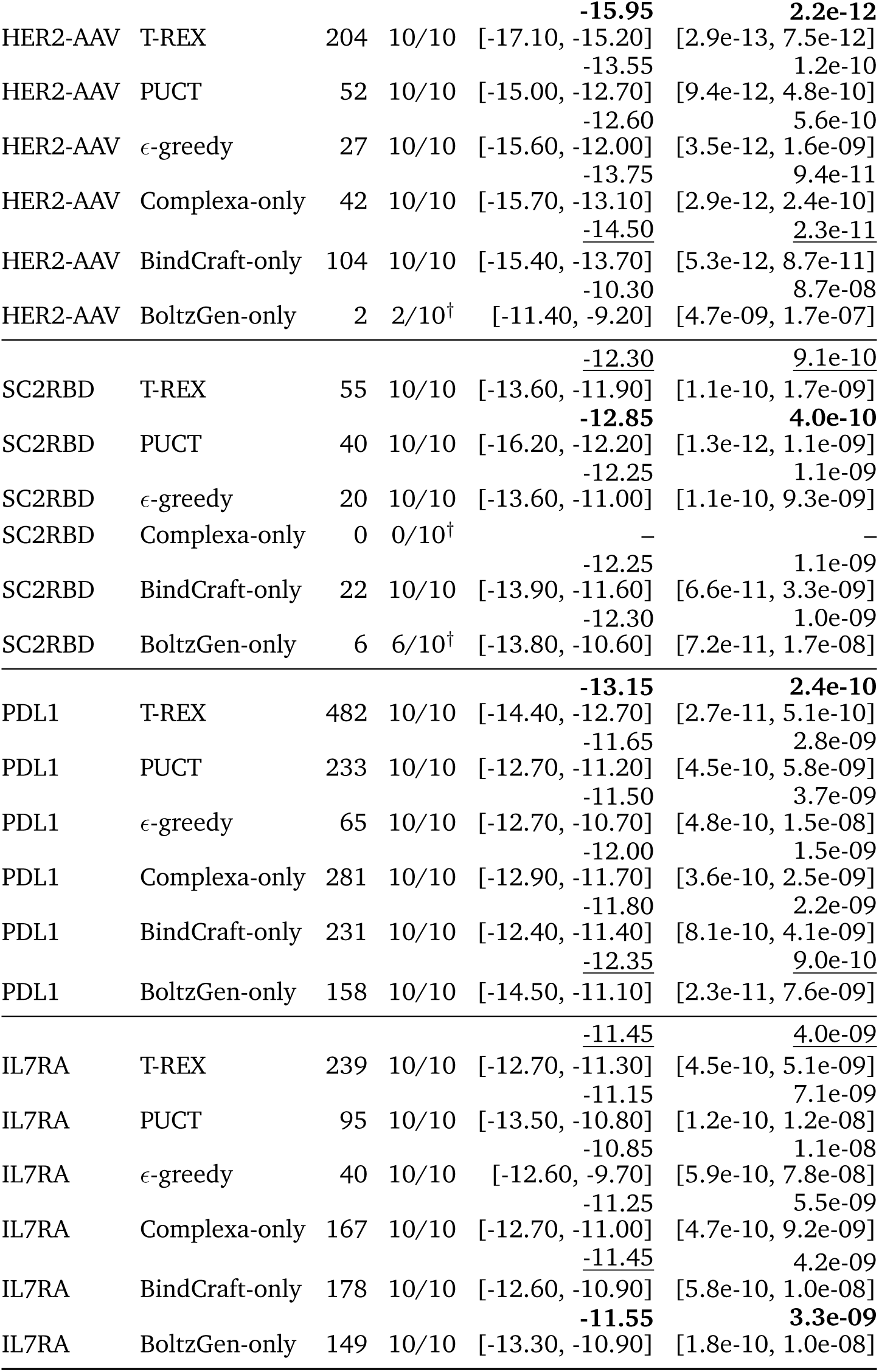
PRODIGY predicted-affinity shortlists. Δ*G* and *K*_D_ report median [best, worst] for the same *K*_D_-ranked top *n* = min(10, *N*) representatives from the method–target pool of size *N*; smaller values rank higher. These are predicted affinities. Within each target and metric, bold and underline mark the lowest and second-lowest distinct unrounded medians among lists of ten designs; ties share a mark. Identical rounded values can carry different marks. Shorter lists carry a dagger and are excluded from highlighting; – denotes an empty pool. Selection rules are given in Section D.4.2.

##### D.4.3. Secondary structure and binding-position diversity

**Secondary-structure annotation and examples.** For Figure 5a,b, we annotated binder secondary structure in the 1,461 final T-REX TM0.6 representatives with MDTraj 1.11.0 (compute_dssp, simplified=True) (Kabsch and Sander, 1983; McGibbon et al., 2015). DSSP states H, G and

I count as helix, and E and B as strand; each fraction uses all binder residues as its denominator, and remaining assignments count as other. The helix-rich and *β*-rich examples maximize helix and strand fractions, respectively, with ties favoring the smaller complementary fraction. The mixed helix/strand example maximizes the smaller of the two fractions, with ties favoring their larger sum. Remaining ties use ascending record-identifier order. Molecular views in Figure 5a,c were rendered with UCSF ChimeraX 1.11 (Pettersen et al., 2021).

To compare *β*-rich output across methods, we selected all qualified designs with at least 20% strand before Foldseek TM0.6 clustering within each method–target pool. This counts clusters within the *β*-rich subset, rather than *β*-rich members of the fixed representative set. T-REX was the only method with such designs on all seven targets and yielded 308 clusters (0.306 per worker GPU-h), compared with 95 (0.094 per worker GPU-h) for BoltzGen-only, the next-highest method (Supplementary Table S23). T-REX yielded more clusters than PUCT on six targets and more than the highest-yielding generator-only method for each target on five. These annotations did not change the primary endpoint.

**Table S23.**
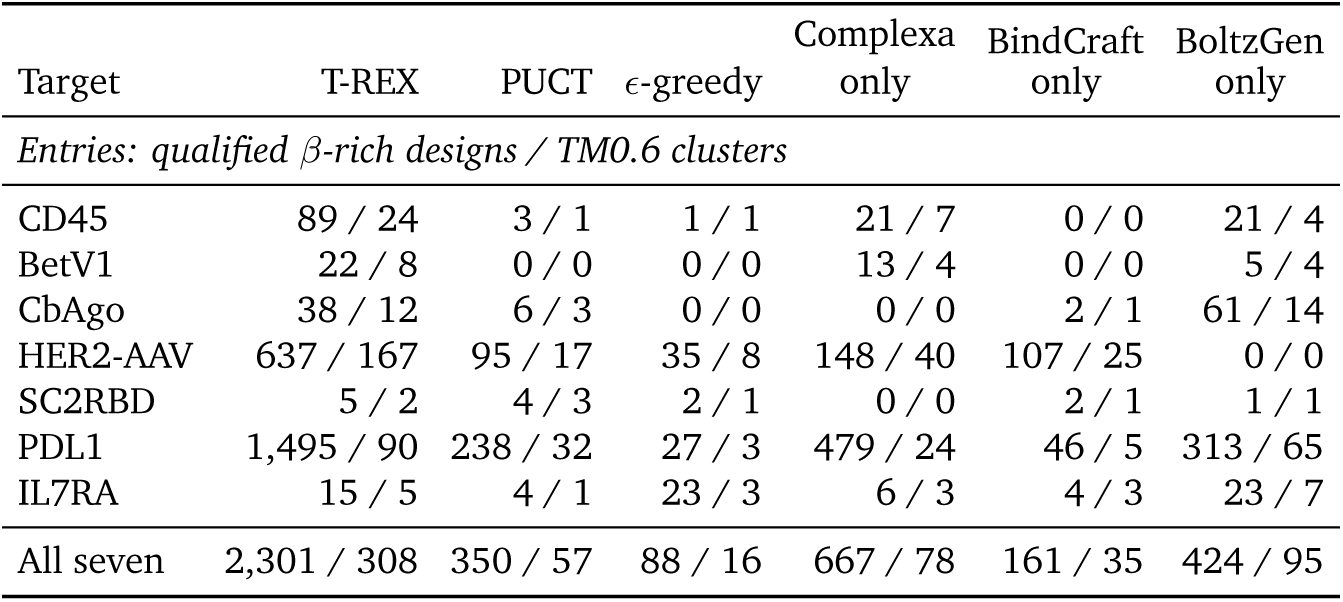
*β***-rich qualified designs and structural clusters by target.** A binder is *β*-rich when at least 20% of its residues are assigned to the strand category by MDTraj’s simplified three-state DSSP implementation. Entries give qualified *β*-rich designs / TM0.6 clusters after reclustering that subset within each method–target pool. The final row gives pooled counts across the seven targets; pooled rates in the text use the fixed reporting budget of 7 × 144 = 1,008 worker GPU-h per method. These post-hoc annotations did not affect allocation, qualification or SU credit.

**Contact-pattern dissimilarity.** To compare contact patterns, we defined a target–binder residue contact as any heavy-atom pair within 6 Å. Each contact feature combined target position with the contacting binder residue’s class: hydrophobic, A/V/L/I/M/C; aromatic, F/Y/W; polar, S/T/N/Q/H; positive, K/R; negative, D/E; glycine, G; proline, P; or other. Repeated contacts with the same position–class pair contributed one element to the design’s feature set. For two nonempty sets *F_i_* and *F_j_*, dissimilarity was 1 − |*F_i_* ∩ *F_j_*|/|*F_i_* ∪ *F_j_*|, ranging from zero for identical patterns to one for no shared features. CbAgo example selection used target residue number and identity to identify positions; the all-target comparison used the sequence alignment described below.

**Target-relative binder positions.** For Figure 5c, we first aligned predicted CbAgo complexes to a common target reference. Alignment used target C*α* atoms shared with the reference at residues contacting a binder in at least 20% of the pooled T-REX and PUCT complexes, with at least 15 matched atoms per complex. The Kabsch rotation and translation were also applied to the binder, whose center was the mean position of its C*α* atoms. PCA was fitted jointly to these three-dimensional centers for all final CbAgo T-REX (*n* = 116) and PUCT (*n* = 54) TM0.6 representatives. The plotted coordinates describe position relative to CbAgo and retain units of Å; axis directions and signs are arbitrary. They do not measure binder fold, orientation or contact-pattern dissimilarity.

To illustrate positional and contact variation among higher-scoring representatives, we selected P1–P3 from the 116 T-REX representatives ranked by the minimum of three scaled margins: (pLDDT − 90)/10, (7/31 − iPAE)/0.05 and (1.5 − scRMSD)/0.3, with scRMSD in Å. This example-selection score differs from *m*_min_ used to choose the final cluster representatives (Section B.2). We retained the upper half (*n* = 58) and chose the highest-scoring design as P1. P2 and P3 each maximized the minimum combined distance to the previously chosen examples: one-half the three-dimensional binder-center separation divided by its 95th percentile across all CbAgo T-REX pairs, plus one-half the contact-pattern dissimilarity.

**Sample-size-matched comparisons.** For the all-target comparison, we compared contact patterns at a common sample size and mapped target positions by global sequence alignment to remove PDB residue-number offsets. For each target, the reference was the longest sequence observed across T-REX and PUCT, preferring the most frequent at equal length. Alignment scores were +2 per match, −1 per mismatch and −2 per gap; ties preferred a diagonal step, then a gap in the query, then a gap in the reference. Only identical aligned residues supplied position mappings. These matches had to cover at least 70% of each sequence and constitute at least 90% of aligned nongap pairs. Contacts without a mapped target position were omitted; every retained design had a nonempty feature set. The resampled CbAgo position comparison also used this sequence mapping, with Kabsch alignment on all matched target C*α* atoms. Figure 5c instead uses the recurrent-contact residues defined above.

For each target and method, we sampled 30 final TM0.6 representatives without replacement and repeated sampling 1,000 times (random seed 20260831); all pools contained at least 30 representatives. Each sample yielded a median pairwise contact-pattern dissimilarity, or, for CbAgo positions, a median three-dimensional Euclidean distance between binder centers. We report the mean and fifth–ninety-fifth percentiles of these resampled medians. The intervals describe variation among subsets of the observed pools, not confidence intervals across independent campaigns.

The CbAgo mean resampled median separation was 31.2 Å [25.2–36.6] for T-REX and 9.97 Å [9.08–11.00] for PUCT. Contact-pattern dissimilarity was higher for T-REX on four of seven targets (Supplementary Table S24). These comparisons support broader predicted binder positions for CbAgo, while the contact-pattern comparison did not favor one method uniformly across targets.

**Table S24.**
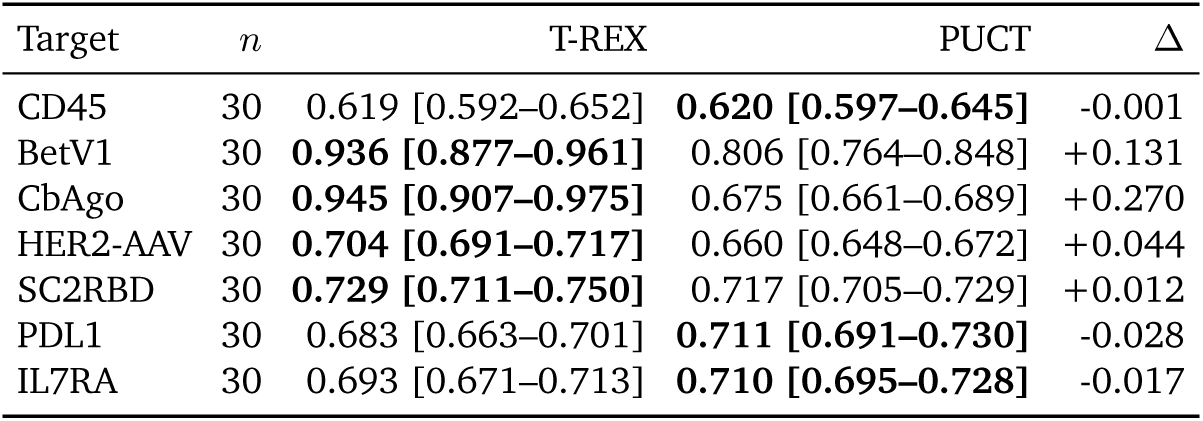
Sample-size-matched predicted contact-pattern variation. For each target, 30 final TM0.6 representatives were sampled without replacement from the T-REX and PUCT pools; both pools contained at least 30 representatives. The method columns report the mean [5th–95th percentile] of the median pairwise dissimilarity across 1,000 samples; *n* is the number of representatives per sample and method. Dissimilarity is one minus Jaccard similarity for 6-Å contact features formed from aligned target-sequence position and contacting-binder residue physicochemical class; higher values indicate more varied predicted contact patterns. Δ is the difference between the two means (T-REX minus PUCT), calculated before rounding. Bold marks the larger unrounded mean in each row. Contact patterns measure a different aspect of structural variation from the binder positions in Figure 5c. Intervals describe variation among subsets of the observed pools, not confidence intervals across independent campaigns.

